# A multiplexed, confinable CRISPR/Cas9 gene drive propagates in caged *Aedes aegypti* populations

**DOI:** 10.1101/2022.08.12.503466

**Authors:** Michelle A.E. Anderson, Estela Gonzalez, Matthew P. Edgington, Joshua X. D. Ang, Deepak-Kumar Purusothaman, Lewis Shackleford, Katherine Nevard, Sebald A. N. Verkuijl, Tim Harvey-Samuel, Philip T. Leftwich, Kevin Esvelt, Luke Alphey

## Abstract

*Aedes aegypti*, the yellow fever mosquito, is the main vector of several major pathogens including yellow fever, dengue, Zika and chikungunya viruses. Classical mosquito control strategies, mainly utilizing insecticides, have had success in controlling other mosquito vectors in recent years, but are much less useful against *Ae. aegypti*, and even these methods are threatened by rising insecticide resistance. This has stimulated interest in new mosquito control mechanisms, notably genetic systems such as gene drives. However, the development of CRISPR/Cas9 gene drive systems has faced challenges such as low inheritance biasing rate, the emergence of resistance alleles, and the possibility of spreading beyond the intended population. Here, we test the regulatory sequences from the *Ae. aegypti benign gonial cell neoplasm* (*bgcn*) homolog to express Cas9 in the germline to find an expression timing more conducive to homing. We also created a separate multiplexing (targeting multiple different sites within the target gene) sgRNA-expressing homing cassette inserted into the *Ae. aegypti kynurenine 3-monooxygenase* (*kmo*) gene to limit the consequences of resistance alleles. This creates a ‘split’ gene drive such that one part does not drive, allowing control over geographic spread and temporal persistence. When combined, these two elements provide highly effective germline cutting at the *kmo* locus and act as a gene drive. Our target genetic element was driven through a cage trial population such that carrier frequency of the element increased from 50% to up to 89% of the population despite significant fitness costs to *kmo* insertions. Deep sequencing suggests that the multiplexing design could mitigate resistance allele formation in our gene drive system.

**Significance statement:** Mosquito-borne diseases affect millions of people worldwide, with the yellow fever mosquito (*Aedes aegypti*) being the principal vector of many viral diseases. Effective measures for controlling this mosquito are sorely needed. Gene drive systems have arisen as a potential tool for mosquito control due to their ability of biasing inheritance of a trait into a target population. Here, we assess a split gene drive, based on CRISPR/Cas9 endonuclease technology driving a target element into the mosquito population. Evaluated over successive generations in a replicated cage trial, the drive successfully biased its inheritance, increasing in frequency from 50% to up to 89%. Our results are encouraging for the potential use of this type of contained gene drive system for mosquito control in endemic areas.

## Introduction

The highly invasive nature of *Ae. aegypti* and its rapid adaptation to human-commensal habitats such as densely-populated cities/towns have played a significant role in the global spread of vector-borne diseases (1, 2). With up to 40% of the world’s population at risk of infection, an estimated 390 million infections per year for dengue alone (3), and a predicted increase in the future due to climate change and urbanization (4), control of the *Ae. aegypti* vector is fundamental to reducing this burden (2). While conventional control methods have had some success in suppressing mosquito populations and the associated burden of disease (2), inherent limitations and reductions in efficacy brought about through insecticide resistance and off-target impacts have highlighted the need for research into orthogonal, effective, and environmentally friendly alternatives - including gene drives (5).

Gene drives are a means of biasing inheritance to spread a trait of interest through a target pest population (6, 7). The development of readily programmable nucleases (clusters of regularly interspaced short palindromic repeats/Cas9 (CRISPR/Cas9)) greatly facilitated the development of nuclease-based drives, notably homing-based drives, with a focus on their potential for mosquito control (6, 8). These gene drives contain a Cas9 endonuclease and at least one single guide RNA (sgRNA) that can be easily reprogrammed to recognize the genomic site of choice. The Cas9 then cuts the DNA creating a double-stranded break which then must be repaired, by the cells own DNA repair machinery. By targeting a site at which components of the gene drive have been inserted into the genome, homology-directed repair will result in conversion of the cut allele into a gene drive carrying allele though a process referred to as homing. Alternatively, the cut may be repaired by non-homologous end joining. There have been demonstrations of the remarkable efficiency of Cas9-based homing drives at biasing inheritance in a few organisms, namely the yeast *Saccharomyces* cerevisiae and *Anopheles stephensi, Anopheles gambiae* mosquitoes (9–11). However this high efficiency has not been replicated in other species such as *Drosophila melanogaster, Ae. aegypti* and mammals such as mice (12–14).

Like any pest management intervention, gene drives are expected to select for resistance in the target organism. Sequence variations in the target loci, caused either by pre-existing heterogeneity or mutations induced from cut-site repair without homing, may lead to the selection of ‘resistance alleles’ (15–17). These resistance alleles can rapidly accumulate in the population if they also maintain function of the target gene, so called ‘r1’ alleles (as opposed to ‘r2’ which are resistant to cleavage but non-functional). In some of the first gene drives tested in cage trials, it was found that resistance led to rapid inhibition of homing and drive (15, 18), a problem that remains to be overcome. Including multiple sgRNAs targeting numerous sequences at the target loci, or ‘multiplexing’ is one potential way to mitigate against this (8); pre-existing sequence variations (whether r1 or r2) or failed homing attempts must have inhibited all target sequences to fully inhibit further drive (19–24). Such a system is furthermore less likely to result in functional resistant alleles, given the multiple disruptions to the target gene and will function most effectively if non-functional mutations result in some fitness cost, as would likely be expected of most genes. Close linkage of the sites may be important for HDR-efficiency, minimizing the sequence length which must be resected, however there is a possibility that NHEJ-based repair at one site may affect the target sequence of closely linked sites, for example if a large deletion is caused (25). Here we investigate the feasibility of a multiplex design in *Ae. aegypti*.

One of the most attractive features of CRISPR/Cas9 gene drives is their potential to spread from very low initial release frequencies (6), but this efficiency is also a cause for concern. The dangers of accidental release or issues around control in the field have promoted interest in less invasive threshold-dependent gene drive systems that are more geographically confinable (26–31). Split-drive systems, where one essential component of the drive is not biased, allow for safe and straightforward optimization and comparison of the different components of the drive, and provides many of the desirable effects of CRISPR/Cas9 homing gene drives with increased control (5–8). While nonlocalized gene drives have been tested in a handful of dipteran species (10, 12), population-level assessment of confinable ‘split-drive’ designs has previously only been empirically demonstrated over multiple generations in *D. melanogaster* (32).

A split-drive system requires separating the drive into two parts, both of which are essential for the drive to function. Fortunately, CRISPR/Cas9-based drives provide a natural split, as CRISPR/Cas9 nucleases have two components: the Cas9 protein and an sgRNA that defines the target sequence. Part of the drive – the component that will home and correspondingly benefit from biased inheritance – is inserted into the target region where the Cas9 will cut guided by the sgRNAs specifically designed for that region. We selected *kmo*, an attractive target gene, for initial gene drive studies and designed a multiplex homing cassette expressing four sgRNAs targeting the *kmo* gene (hereafter referred to as *kmo*^sgRNAs^). *kmo* is required for the synthesis of ommochrome pigments in mosquitoes; homozygotes for non-functional mutant alleles display a white eyed phenotype (33, 34). The recessive eye phenotype allows easy tracking of insertional mutants (also marked by a fluorescent protein), as well as other non-functional mutations resulting from ‘error-prone’ DNA repair pathways such as non-homologous end joining (NHEJ) (33). Mosaicism observed in the eye can be a useful indication of somatic expression or deposition of Cas9/sgRNAs into the embryo (33, 34).

Indels generated in somatic tissues could result in a phenotype similar to homozygotes if cut rates are high in the relevant tissues. In drives that target haplosufficent female sterility genes this could result in females carrying the drive elements becoming sterile themselves instead of the desired heterozygous, fertile phenotype (10, 35); somatic expression of active nuclease in heterozygotes is therefore undesirable. The second element of the split-drive assessed here utilizes the regulatory elements from *Ae. aegypti bgcn* to express Cas9. *bgcn* has been identified and characterized as a regulator of cystoblast formation in *D. melanogaster* where transcripts are restricted to a few cells, including germline stem cells (35). This restricted expression pattern is favorable for confining Cas9 expression to the germline and minimizing somatic expression/cutting (36).

Each element on its own will be transmitted between generations under standard Mendelian principles and rates of inheritance. However, when these two components come together in a single organism, the desired outcome is cleavage of *kmo* in the germline, allowing the *kmo*^sgRNAs^ element to be utilized as a template for homology-directed repair (HDR) and so bias inheritance in its favor (“drive”). In simple crosses between trans-heterozygotes (*kmo*^sgRNAs^; *bgcn*-Cas9) and WT, we observed an inheritance rate of the *kmo*^sgRNAs^ element of greater than 75%. In our small cage trials, we observed highly effective germline cutting rates and the split-drive was able to bias inheritance such that, after several generations, up to 89% of a population carried the element. These results demonstrate the ability of split CRISPR/Cas9 homing drives to increase in frequency within *Ae. aegypti* populations over multiple generations and validate previous modeling work predicting the general dynamics of this type of system.

## Results

### Design and generation of split-drive elements

It has been proposed that multiplexed designs may mitigate the formation and accumulation of resistance alleles due to the use of multiple target sites. To investigate this hypothesis we designed an array of four different RNA pol III promoters, each expressing a different sgRNA targeting four, closely linked, sequences in *kmo* (Fig 1a). *Ae. aegypti* endogenous pol III promoters were selected based on expression in *Ae. aegypti* Aag2 cells and *Ae. aegypti* transgenics (37, 38). Three previously verified (39) and one new sgRNA target were selected within a region of approximately 135bp, each are expressed with one of four unique backbone variants (40) to minimize repetitive sequences within the construct. This plasmid (AGG1095) uses a 1.2kb 5’ homology arm and a 1.9 kb 3’ homology arm to integrate the multiplexed sgRNA array and an AmCyan fluorescent marker into the genome. The homology arms exclude this 135 bp of *kmo* exon five, which contains all four sgRNA target sites (Fig 1a, S1), such that these are absent from the drive allele. It should be noted that this 135bp region includes sequence beyond the cut sites of even the outermost sgRNAs (Fig 1a, top line), such that even the end sgRNAs do not have precise homology at even one end. This avoids the outermost sgRNAs having a privileged position relative to the internal ones (without the requirement for resection) and addresses the critical question of whether this multiplexing design leads to a dramatic reduction in drive efficiency as has been demonstrated in *D. melanogaster* (41).

**Figure 1.**
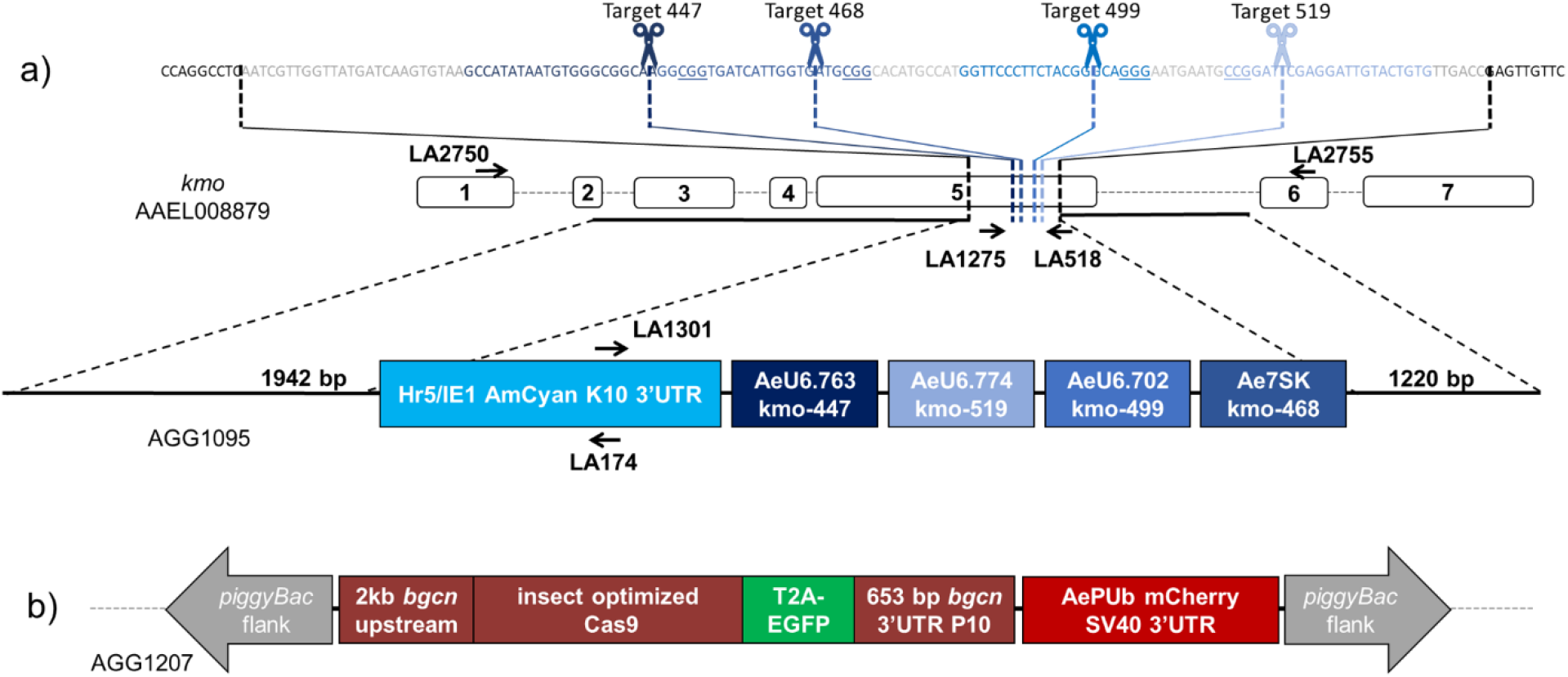
Split-drive designs. a) Four sgRNA targets were selected within exon 5 of the *kmo* (AAEL008879) gene (dashed blue lines) of *Ae. aegypti*, disruption of which results in loss of pigmentation in the eye. The 135bp region containing all four targets which is excluded from the plasmid, is depicted above with the cut sites of each sgRNA indicated by scissors and blue text, PAM sites are underlined. It should be noted that cuts at any of the sgRNA target sites would require resection to reach perfect homology (nucleotides between black dashed lines indicated in grey). Upstream and downstream regions were sequence confirmed in the Liverpool strain (WT) and the region of homology included in the plasmid is indicated by black bars. AGG1095 plasmid consists of an Hr5/IE1 expressing AmCyan fluorescent marker, and four endogenous *Ae. aegypti* RNA pol III promoters each expressing the specified sgRNA. b) The *bgcn*-Cas9 expression plasmid uses ~2 kb upstream sequence of the *bgcn* gene to express an insect codon optimized Cas9 followed by T2A-EGFP and the *bgcn* 3’UTR and an additional P10 3’UTR. A PUb-mCherry marker also contained within the *piggyBac* transposable element flanks allows for identification of transgenic mosquitoes. Individual elements are not to scale.

Embryonic microinjection with *in vitro* transcribed sgRNAs, plasmid AGG1095, and Cas9 protein generated several transgenic lines positive for the fluorescence marker (Table S1). Integration into the *kmo* gene was confirmed by PCR (Fig S4). All further investigations were carried out using a line derived from a single PCR confirmed G_1_ male (*kmo*^sgRNAs^).

Embryonic microinjections into Liverpool strain (‘WT’) with the *bgcn*-Cas9 construct and *piggyBac* transposase (Fig 1b, S1) yielded at least five insertion events (Table S1). These five transgenic lines were assessed for the absence of sex-linkage and multiple insertions. Based on these criteria, three *bgcn*-Cas9 lines (B2, C, D) were selected and further assessed for homozygous viability (Table S2) and one was selected to determine its ability to bias the inheritance of the *kmo*^sgRNAs^ element in a standardized series of crosses.

### Determination of inheritance bias by bcgn-Cas9

For the selected *bgcn*-Cas9 insertion line (D), we first crossed *bgcn*-Cas9 females to *kmo*^sgRNAs^ males and termed this the F_0_ cross (Table S3). Trans-heterozygous (*kmo*^sgRNAs^; *bgcn*-Cas9) F_1_ progeny were then crossed to Liverpool WT of the opposite sex, and their progeny scored for inheritance of the *kmo*^sgRNAs^ element (Fig 2a, Table S4). These progeny (F_2_) were collected as pools, separately for each lineage of crosses. We observed super-Mendelian inheritance of the *kmo*^sgRNAs^ (G-test: *G*_1_ = 90.875, p< 0.001, Fig 2a, Table S4 and S5). For this line, which showed evidence of inheritance bias for both sexes (68% in males *G*_1_ = 15.221, p< 0.001; 77% in females *G*_1_ = 98.201, p< 0.001, Table S6), we next set out to more accurately quantify the rate of inheritance bias, the overall germline cutting-rate and relative contribution of the individual sgRNAs.

**Figure 2.**
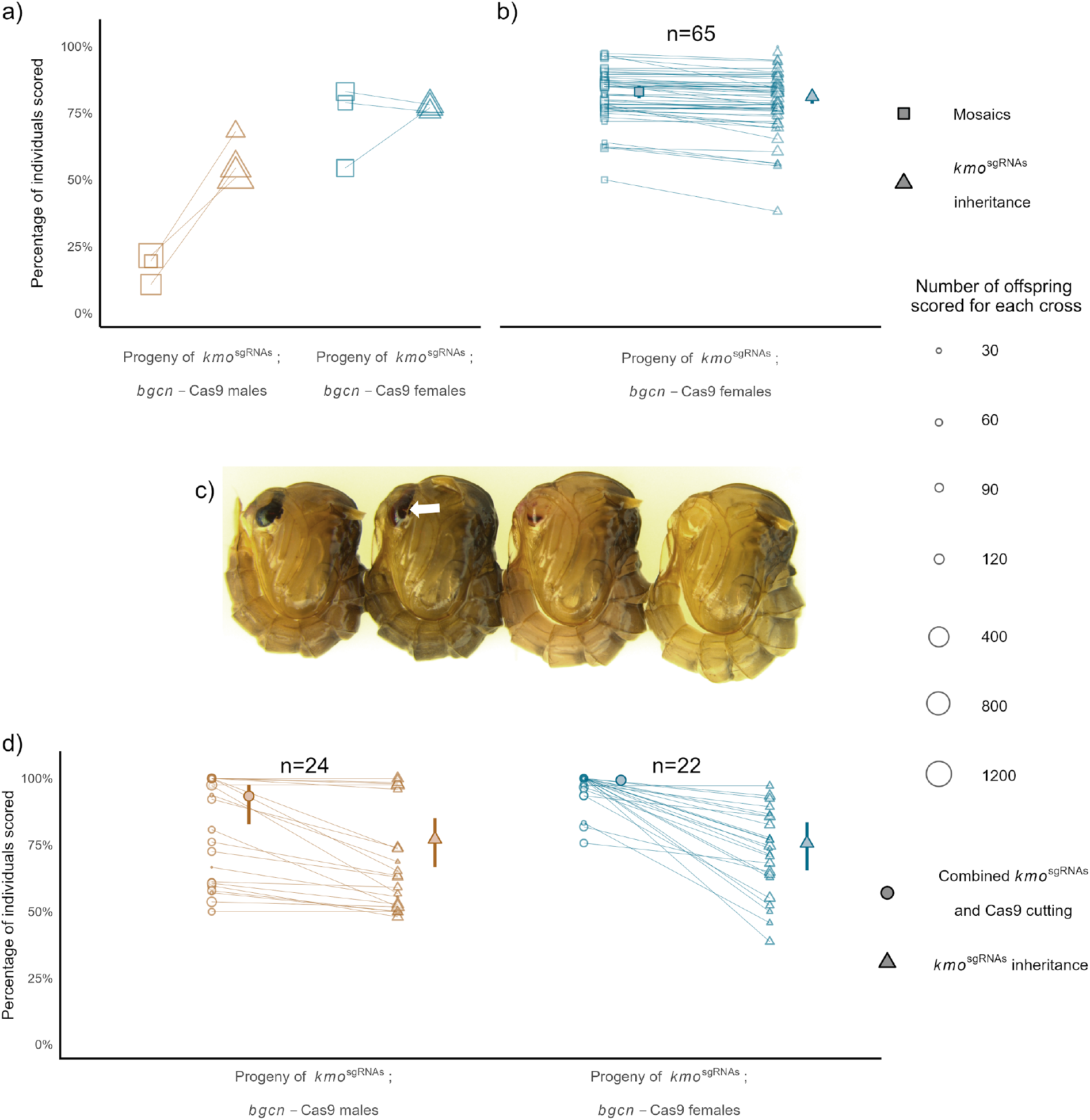
*bgcn-Cas9* line D causes biased inheritance of *kmo*^sgRNAs^. a) Female *bgcn*-Cas9 were crossed to *kmo*^sgRNAs^ males (F_0_) for an initial pooled homing assessment. Trans-heterozygous F_1_ males and females were outcrossed to WT in three replicate crosses and the F_2_ progeny scored for inheritance of the *kmo*^sgRNAs^ transgene (triangles) and eye phenotype (squares). b) A replicate cross was completed and the proportion of the F_2_ progeny inheriting the *kmo*^sgRNAs^ transgene (triangles) and eye phenotype (squares) were scored from individual F_1_ females. c) Images of pupae displaying the different eye phenotypes from left to right: wild type (dark), weak mosaic (white arrow indicating mosaicism), strong mosaic, white eye. d) Combined kmo^*sgRWAs*^ inheritance and Cas9 cutting are represented by circle points and are measured as the percentage of offspring with white eyes from crosses between males (n=24) or females (n=22) trans-heterozygous for *kmo*^sgRNAs^; *bgcn*-Cas9 and *kmo^-/-^* mosquitoes. *kmo*^sgRNAs^ inheritance alone is measured as the percentage of offspring with AmCyan fluorescence (triangles). Individual faded points represent offspring from one drive parent, and the size of the point is proportional to the number of offspring from the parent. Horizontal lines connect pools of offspring to show the relationship between cutting and inheritance rates in each replicate. Large symbols and error bars (vertical lines) represent estimated mean and 95% confidence intervals calculated by a generalized linear mixed model, with a binomial (‘logit’ link) error distribution. Data points may be overlapped and some cannot be discerned.

### Determination of inheritance bias and cutting rates using the split-drive system

To assess the overall cutting and homing efficiency of the drive, trans-heterozygous *kmo*^sgRNAs^; *bgcn*-Cas9 (F_0_ *bgcn*-Cas9 females crossed to *kmo*^sgRNAs^ males) themselves having a mosaic phenotype were crossed to a gene-edited *kmo* knock-out line (*kmo^-/-^*) line as single pair crosses and the progeny of each individual cross was scored separately. The offspring of this cross were screened for AmCyan fluorescence, indicating the inheritance of the *kmo*^gRNAs^ allele, and for eyes phenotype. The drive was inherited by 77.2% ([approx. 95% CI] = [66.8-85.1%]) of the progeny of the male trans-heterozygotes and 75.7% [65.5-83.6%] of the progeny of the female trans-heterozygotes (Fig 2d, Table S7, Model 1), substantially higher than predicted odds from Mendelian inheritance rates of 50% (*β* = 1.18 [0.77-1.60], p< 0.001, Table S8, Model 2), and with no significant effect of parental origin for the Cas9/drive allele (Odds ratio = 0.92 [0.45-0.1.88]), *p* = 0.816, Table S8, Model 1, Fig 2d). These estimates, which incorporated batch effects, were slightly elevated from the pooled data for homing, especially for males (males 71.4% [69-73.8%], females 72.9% [70.2-75.4%] (Fig 2a, Table S7, Model 4), indicative of a significant level of individual variation in homing efficiency. As a comparison, the inheritance of the Cas9 allele (which should conform to standard Mendelian inheritance), was 48.8% for males, and 50.1% for females (*β* = −0.05 [-0.2-0.1], p = 0.474; *β* = 0.05 [-0.16-0.26], p = 0.588), and indicates no major effect on viability of the *bgcn*-Cas9 allele under these conditions.

In this experimental design the only functional *kmo* allele is in the chromosome homologous to the *kmo*^sgRNAs^ allele in the trans-heterozygous parent. Progeny of this cross which lack the cyan fluorescence marker indicative of the *kmo*^sgRNAs^ element must have inherited this other, nominally wild type *kmo* allele. In such individuals, completely white eyes indicate that this allele has been mutated through cutting and error-prone repair, likely either in the germline of the trans-heterozygous parent or very early in the developing zygote, or in principle, one or more later events that still affect all relevant cells providing pigmented ommatidia. Mosaic eyes indicate non-functional mutations which were generated later in the developing zygote, such that some cells forming ommatidia have *kmo* function, but others do not (Fig 2d, Table S8). We observed white eyes in F_2_ progeny at a rate of 93.4% [82.8-97.6] from F_1_ males and 99.3% [97.5-99.8%] from F_1_ females indicating that while germline/early zygotic cutting efficiencies are very high in both sexes, offspring of trans-heterozygous females show significantly higher rates of cutting compared to the offspring of males (Odds ratio = 10.47 [2.16-50.78], p = 0.004, Table S8, Model 4), possibly due to additional cutting from maternally deposited nuclease activity, however this may also be reflective of different germline activity between the sexes.

White eyed F_2_ larvae which did not inherit the *kmo*^sgRNAs^ were collected for deep sequencing to determine the relative cutting efficiency of each sgRNA (Table S12). It is important to note that the mutations observed in these individuals include mutations that were originally present in and contributed by the *kmo^-/-^* line (Table S9). Results from this deep sequencing can therefore only indicate the relative frequency of mutation caused by respective sgRNAs but do not indicate the timing at which nuclease activity occurred or if cuts by a certain sgRNA bias HDR over NHEJ upon cleavage. We determined the prevalence of mutated nucleotides in the sequence reads relative to the wild type *kmo* sequence. We found a wide range of efficiency for each unique pol III expressed sgRNA. sgRNAs 447 (U6.763) and 499 (U6.702) resulted in approximately 60-66% of the alleles cut. sgRNA 468 (Ae7SK) had ~40% of alleles cut and for 519 (U6.774) a mere 1.6-5.2% of alleles were mutated (Fig 3). Simultaneous cuts between the two outermost sgRNAs (447 and 519) would generate deletions of 72 nt that eliminate all four sgRNA targets and create a fully cut-resistant allele. We did not observe any such deletions among the 97 non-*kmo*^sgRNAs^ inheriting larvae collected (Table S10). Deletions which span between the cut sites of sgRNA 447 and 499 or sgRNA 468 and 519 would be 51bp and would eliminate 3 target sites. Only in the samples of non-transgenic white eyed F_2_ larvae from trans-heterozygous females did we identify any reads with deletions larger than 50 nt (maximum of 4.63%, Table S10). Much larger deletions that could remove one or both primer binding sites would not be detected by this assay but the distribution of deletions that we did observe suggests such larger deletions are very rare, unless generated by a different mechanism. Having characterized the isolated metrics of the drive, we next set out to test its performance at the population level.

**Figure 3.**
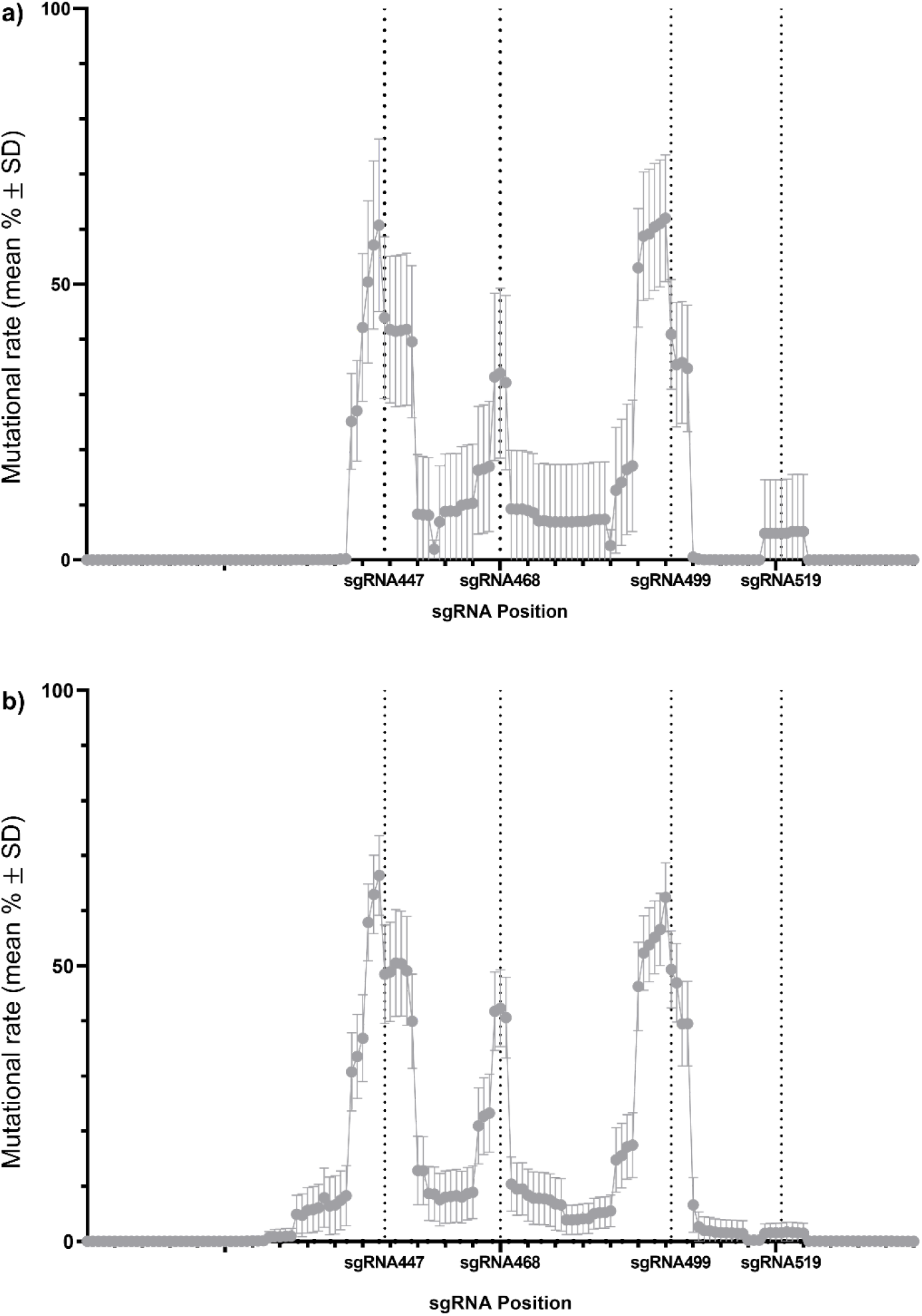
Mutational rates vary for *kmo* targets sgRNA447, sgRNA468, sgRNA499, and sgRNA519. Dotted lines represent the expected cut sites of corresponding sgRNAs. a) White-eyed F_2_ larvae which did not inherit the *kmo*^sgRNAs^ transgene (non-transgenic or inheriting only the *bgcn*-Cas9 element) from the ♂*kmo*^sgRNAs^; *bgcn*-Cas9 x ♀*kmo^-/-^* cross. b) reciprocal cross to (a): white-eyed F_2_ larvae which did not inherit the *kmo*^sgRNAs^ transgene from the ♂*kmo^-^*x ♀*kmo*^sgRNAs^; *bgcn*-Cas9 cross.

### Cage trials

To evaluate the ability of the split-drive design to spread through a WT population, we initiated two replicate experimental cages (A1 and A2) by mating 100 mosaic eyed female *kmo*^sgRNAs^;*bgcn*-Cas9 trans-heterozygotes to 100 wild type males (F_0_) and monitored both transgenes for six generations (Fig 4b, Fig S6, Dataset S1). Although such a population set up may not be realistic of the potential use of such a system in the field, it was chosen to allow robust data to be collected on the dynamics of spread of this proof-of-principle system in a reasonable time-frame. In the F_1_ generation we observed an increase in the proportion of the population carrying the *kmo*^sgRNAs^ element followed by a small decrease in the F_2_ generation (79% to 76.5% in A1 and 77.4% to 75% in A2). In the F_3_ generation the frequency of the *kmo*^sgRNAs^ substantially diverges between the replicate cages (74% in A1 and 88% in A2), presumably due to stochastic effects, but still remains within the model-predicted range. We observed a maximum *kmo*^sgRNAs^ frequency of 89% in these small cage populations, consistent with the upper end of the stochastic model prediction (Fig 4b, Dataset S1).

**Figure 4.**
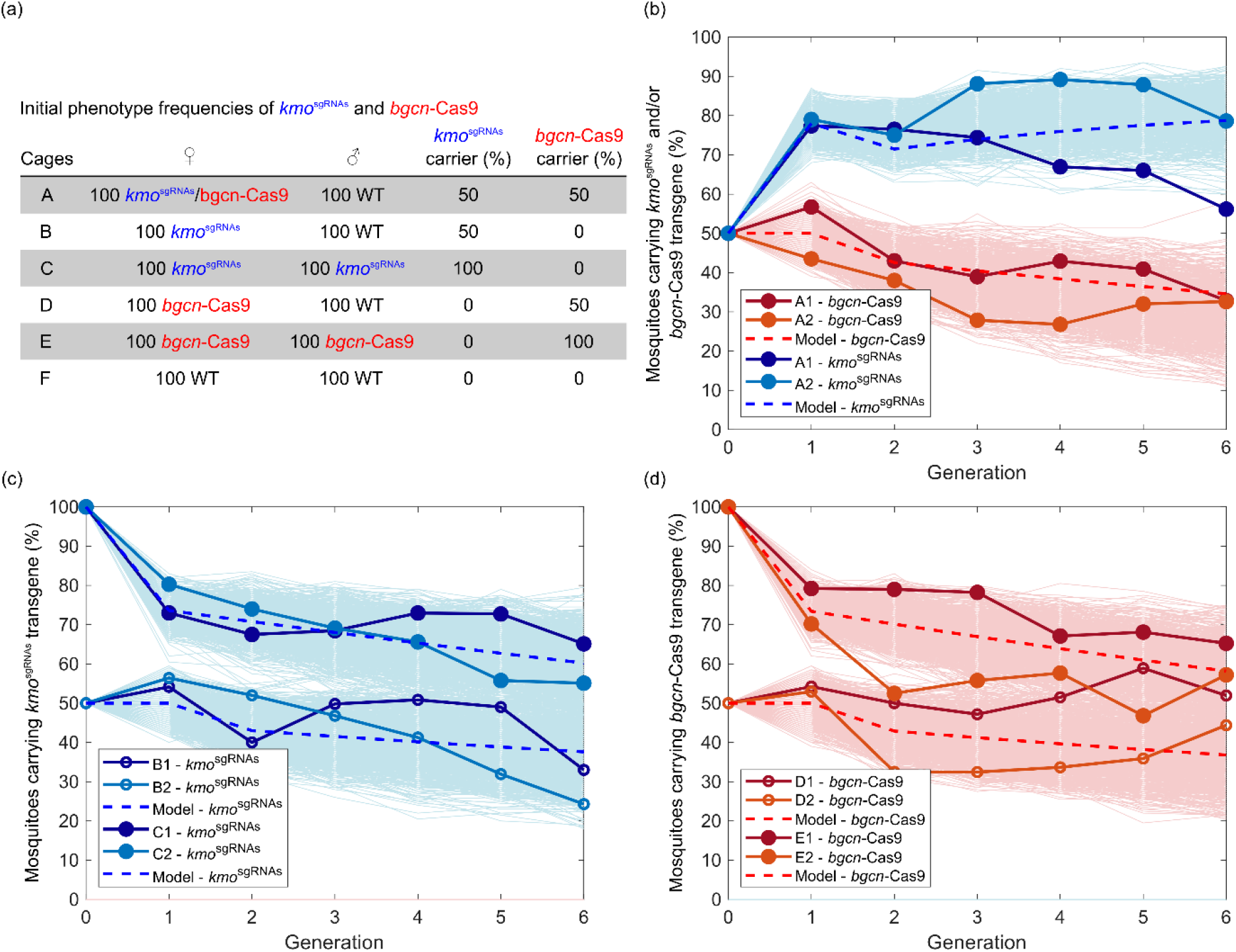
*bgcn*-Cas9; *kmo*^sgRNAs^ split-drive can increase in frequency through a caged population. Presence of *kmo*^sgRNAs^ and *bgcn*-Cas9 elements observed for 6 generations of small, caged populations and the percentages predicted by the deterministic (dashed lines) and stochastic (pale lines) models. Cages each generation beyond the initial setup were begun with 250 L1 larvae. a) Initial frequencies for each cage at the outset of the trial. b) Percentage of individuals carrying the *kmo*^sgRNAs^ transgene (blue solid lines) and/or the *bgcn*-Cas9 transgene (red solid lines). c) Percentage of *kmo*^sgRNAs^ transgene in the absence of *bgcn*-Cas9 when female heterozygotes are crossed to either male heterozygotes (filled circles) or to wild type (WT) males (open circles). d) Percentage of *bgcn*-Cas9 transgene in the absence of the *kmo*^sgRNAs^, again with female heterozygotes crossed with male heterozygotes (filled circles) or to WT males (circles). The stochastic results show the behavior produced by each of 1,000 independent numerical simulations. Details for cage F are included in Supplemental Dataset S1.

By also noting the eye color phenotype through the generations of the cage trial we can gain insight into NHEJ rates in individuals which do not carry either element. A mosaic eyed phenotype we take to be an indication of embryonic deposition of drive components. For individuals carrying the *kmo*^sgRNAs^ transgene the mosaic eye phenotype frequency decreased from the 100% in the initial trans-heterozygotes (F_0_) to about 70% in the F_1_ generation and stayed below 10% thereafter, as was similarly observed in our initial test crosses (Table S3 and S5). In those which did not carry the *kmo*^sgRNAs^ transgene, mosaicism was about 25% in the second generation. This is similar to rates we observed in the test crosses (Table S7). However, from then on, no mosaicism was observed, except for a small number of individuals in the fourth generation in cage A1 (Dataset S1).

In the experimental cages, a complete white eyed phenotype indicates that both *kmo* alleles are disrupted. In those mosquitoes which carry the *kmo*^sgRNAs^ element, we could observe white eyes in individuals with the *kmo*^sgRNAs^ element either homozygous or heterozygous with the other *kmo* allele disrupted by a non-functional mutation. We observed an increase in the frequency of white eyes in *kmo*^sgRNAs^ mosquitoes reaching a maximum of 89.15% (A1) and 82.44% (A2) in the third generation (Dataset S1). In mosquitoes which do not carry the *kmo*^sgRNAs^ element the presence of white eyes corresponds to disruption of both *kmo* alleles. In those mosquitoes which did not carry the *kmo*^sgRNAs^ element, a maximum of 60% were observed with white eyes.

### Multiple sgRNA recognition sites remained intact after the fifth generation of the trial

Having found that large deletions which could remove all sgRNA recognition sites rarely occurred (Table S10), we investigated the existence of alleles which could be cut-resistant at each recognition site without a deletion of the entire region, i.e. multiple SNPs or indels. We collected mosaic and white eyed individuals from the experimental cages (A1 and A2) at generations F_2_, F_4_, and F_5_ for deep sequencing (Table S13). We predict that alleles which are cut-resistant at multiple target sites are likely to present with a null phenotype and dark eyed individuals were deliberately excluded to prevent over-amplification of the WT allele. The proportion of reads still cleavable [retaining the original target and PAM recognition sequence] by one or more sgRNAs was calculated and plotted (Fig 5). We also included in this analysis the ‘at least one sgRNA’ category which indicates the proportion of reads cleavable by one or more of the four possible sgRNAs. In this diagram a higher percentage (y-axis) indicates a greater abundance of cleavable reads from the samples. Two replicates of five wild type adults (20 *kmo* alleles) each were also analyzed to assess the prevalence of naturally occurring SNPs, relative to the reference sequence, that are present in our wild type population. Interestingly, an average of 94.9%, 92.6%, 89.7%, 86.7% and 96.3% of reads from the background strain were shown to be cleavable by sgRNAs 447, 468, 499, 519, and at least one of the sgRNAs, respectively. This indicates an existing prevalence of alleles resistant to cutting by at least one of the four sgRNAs in our cage population before the cage trial was carried out, with resistance being present at the highest level for sgRNA 519.

**Figure 5.**
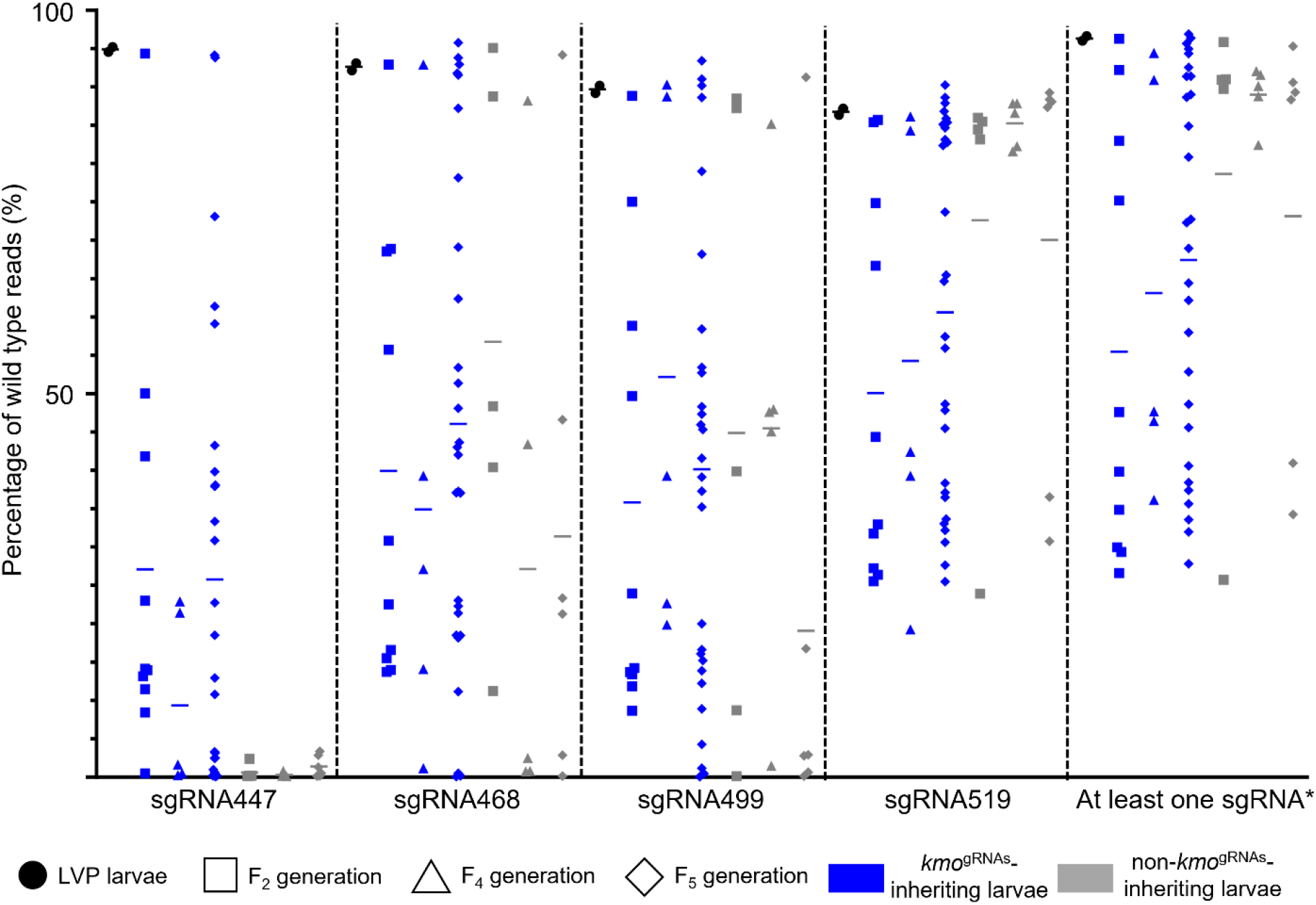
Individuals collected from the cage trial have wild-type *kmo* alleles. Liverpool (WT) adults are represented by black circles while adults from F_2_, F_4_, and F_5_ generations of the cage trial are represented by square, triangle, and rhombus shapes, respectively. Samples are separated by their inheritance of the *kmo*^sgRNAs^ element (blue for inheriting mosquitoes, grey for non-inheriting mosquitoes). Mean percentages are shown as solid lines in the plot. *Percentage of reads having at least one sgRNA recognition sequence unedited.

The proportions of cleavable reads obtained from the individuals collected during F_2_, F_4_, and F_5_ generations of the cage trial have shown <28% of reads from *kmo*^sgRNAs^-inheriting individuals and <2% of reads from non-*kmo*^sgRNAs^ inheriting individuals, on average, still cleavable by sgRNA 447 (Fig 5). The availability of cleavable reads for this sgRNA was consistently lower than the other sgRNAs in each generation assessed, in line with previous observation of the high cutting efficiency of sgRNA 447 (Fig 3). In comparison, the average percentage of cleavable reads fluctuated between 27%-57% and 19%-53% for sgRNAs 468 and 499, respectively between the three generations. Approximately 50%-89% of reads were still cleavable by sgRNA 519 or at least one of the multiplexed sgRNAs (Fig 5). This suggests that while alleles cut-resistant to sgRNA 447 had formed and accumulated to a high level in the population, a substantial subset of these alleles should still be able to be cut by another sgRNA even at generation F_5_.

### Modeling of drive behavior

Fitness effects of the two transgenic constructs used in this study were explored using deterministic, discrete generation, population genetics mathematical model. A stochastic modeling framework (23) was also used to provide a prediction as to the potential range within which we would expect experimental results to vary. We begin with a simple model describing the behavior of a single transgenic construct (i.e., in absence of homing) and use a simple least-squares regression approach to obtain fitness parameters for heterozygous and homozygous individuals. Full details of the deterministic and stochastic mathematical models and parameter fitting procedure are given in Supplemental information S4. Briefly, for *bgcn*-Cas9 the best fit of the deterministic model to the experimental data is obtained where heterozygous and wild type individuals are equally fit while homozygotes have a fitness cost of 21% (S4 Fig S1, with model output in Fig 4d). Non-exclusive potential explanations of such a fitness cost could be, for example, a deleterious threshold of Cas9 expression, insertional mutagenesis at the target site, or insertion linked to deleterious recessive alleles (42). Using the same approach, we obtain a best fit for *kmo*^sgRNAs^ where heterozygotes have wild type fitness and homozygous individuals have a fitness cost of 19% (S4 Fig S2, with model output in Fig 4c). There is contradictory evidence relating to fitness costs of *kmo* in the literature, most notably a high load observed in *An. stephensi*, although knock-outs and knock-ins were previously described in *Ae. aegypti* fitness effects were not measured (18, 39). In our own recent experience with *Culex quinquefasciatus kmo^-/-^* could be generated and maintained as homozygotes, but an insertional mutant expressing a fluorescent protein was homozygous lethal (43, 44). Using these best fit parameter values within the stochastic model shows that experimental results fall within the expected range. While these parameters produced the best fit of the deterministic model to experimental data, Figures S1(a) and S2(a) from S4 demonstrate a range of relative fitness parameters that can produce a similarly good fit.

We then utilize a deterministic population genetics mathematical model including both transgenic constructs and the effect of homing to predict the behavior observed within the experimental treatment cages (full details are available in S4). This model was parameterized using directly measured homing rates (Fig 2) and the fitness parameters obtained above. For the remaining genotypes (i.e., those carrying both constructs) additive and multiplicative combinations as well as independent least-squares regression for the fitness of each genotype were compared. Each approach yielded only a marginal difference in the goodness of fit. We therefore considered additive parameter combinations since they provide a simple and intuitive explanation of interactions between multiple fitness parameters. These were used to predict the behavior of the split-drive system using both deterministic and stochastic mathematical models, giving a fit to experimental data that is broadly within the expected range (Fig 4b). This suggests that our assumed model of the drive behavior and all parameters derived here provide a good understanding of the system (at least in our cage trial setting). We found some minor differences between the model predictions and the experimental data that are likely caused by factors not considered within these mathematical models (e.g., multiplexed sgRNAs, end-joining mediated resistance and maternal deposition of transgenic constructs). We also note that fluctuations due to stochastic effects appear larger in the experimental results than in the results of our models. This is potentially due to the effective population size being lower than the census size of the caged populations, with the latter being used within our models.

## Discussion

Multi-generational lab trials are a critical step towards assessing the utility of novel gene drive systems in the field by considering complex fitness components such as fecundity, longevity, and mating competition. Here we evaluated the spread of a CRISPR/Cas9 multiplexing split-drive in multi-generational, caged, lab populations of *Ae. aegypti*. Using regulatory sequences from the *Ae. aegypti bgcn* homologue to express Cas9 in the germline and a separate sgRNAs expressing homing cassette integrated into the *Ae. aegypti kmo* gene; we demonstrated highly effective germline cutting rates and bias in the inheritance of our genetic element such that the frequency of individuals carrying at least one allele increased from an initial 50% up to a maximum of 89% in 5 generations, in line with the upper bound predicted by stochastic modeling. These results showed an improvement in the inheritance-bias in this mosquito species compared to previous studies (38). Over the course of our cage trial, the drive produced substantial increases in cut-resistant alleles across all four target sites, but no deletions which removed all four target sites. A complementary strategy to managing target-site sequence variation has targeted highly conserved and, ideally, functionally constrained sequences with a single sgRNA (10, 45). This strategy has proved highly effective and combining these approaches would likely improve gene drive conversion efficiencies through further reduction of resistance allele formation, although new designs may be required as the highly conserved RNA sequences used to date are likely not large enough to identify multiple sgRNA targets. More complex strategies such as targeting and recoding essential genes could also be used which should provide selection against r2 alleles and allow the target sgRNA expressing allele to approach fixation (21, 32, 36, 46).

Model fitting to the cage trial data showed that both elements of the split-drive carried moderate fitness costs expressed in homozygotes which may have impeded the rate of spread in the drive, and prevented the drive from reaching fixation. We found a significant difference in observed mean homing rates in the offspring of males between pooled and individual mating crosses. This was likely due to mating competitiveness, as in individual crosses all males have an equal chance to mate, but in pooled crosses some genotypes may contribute fewer offspring to the next generation. We also found high variability in the apparent homing and cutting rates between individuals of both sexes in our individual mating crosses. This likely represents several different fitness costs at work. Taking into consideration batch effects with individual crosses we found homing to be the same between male and female F_1_. The insertion sites of the transgenes in this study played an essential role in Cas9 expression and efficacy. Our selected *bgcn*-Cas9 transgenic was found to have significant recessive fitness costs. Analysis of additional insertion sites or use of insertions into the endogenous locus could yield lines which maintain the ability to bias inheritance or even improve upon it, with lower associated fitness costs. Similarly, *kmo*^sgRNAs^ also showed significant recessive fitness costs, which was not anticipated when this program was initiated. In order to maximize the efficiency of a “population modification” drive, it will be important to identify “cargo” elements and drive insertion sites that minimize fitness costs.

The control of the expression of Cas9 in gene drive systems is critical, as expression either too late in the germline or in somatic cells is likely to result in repair by NHEJ and the formation of cut-resistant alleles (36, 47). *bgcn* has been identified and characterized as a regulator of cystoblast formation in *D. melanogaster*. Transcripts are restricted to a few cells, including germline stem cells. This pattern should be ideal for confining Cas9 expression to the germline and minimizing mosaicism. Our results, however, indicate some somatic expression which means that either our transgene could not recapitulate the endogenous expression pattern of this gene and/or there are significant differences in the expression pattern between *D. melanogaster* and *Ae. aegypti*. In publicly available *Ae. aegypti* RNA-Seq datasets *bgcn* was found to be expressed in females in ovaries both pre- and post-bloodmeal as well as male and female brains but more precise localization data is not available (48, 49). In *D. melanogaster bgcn* was identified by *in situ* hybridization to be restricted to germline stem cells and cystoblasts in the germarium of ovaries, and highly expressed by Northern blot in males testes (50). There is clear scope for the identification of further germline specific genes which can be used either in the endogenous context or whose regulatory elements could be used to express nucleases from a transgene construct. The observed differences in homing between males from pooled and individual crosses may have captured the effect of small fitness loads in the heterozygotes (or those which are strong mosaics and thus somatically homozygous) on male sexual competitiveness. In pooled crosses, males with minimal transgene expression may gain disproportionate shares of reproduction, though the underlying mechanism for fitness costs is unknown. Previous work in this system has also noted the high levels of individual variation we observed in our study of homing rates across both sexes (38). One of the strengths of split gene-drive systems is that it allows future work to test new constructs in different combinations, which would allow these issues to be addressed in the future.

We speculate that maternal deposition of Cas9/sgRNAs in our drive may have acted to increase the inheritance rates of the *kmo*^sgRNAs^ rather than resulting in NHEJ, perhaps due to the multiplex design. In split-drives using a single sgRNA target, maternal deposition often resulted in early embryonic cutting favoring NHEJ rather than HDR, generating resistance alleles at the expense of homing (29, 32, 41, 51). With additional target sites still available in our design there may be additional opportunities for deposited Cas9 to cleave within a later HDR-conducive window resulting in some level of transgenerational effect (“shadow drive”), (51, 52). Further studies into additional germline specific promoters could improve inheritance rates and decrease NHEJ resulting from somatic expression and/or deposition of Cas9.

## Materials and methods

### Plasmids and cloning

#### Design and cloning of *kmo^sgRNAs^* multiplexed sgRNA expression construct

To generate the *kmo*^sgRNAs^ knock-in plasmid we first sought to sequence confirm the *kmo* locus of our Liverpool strain. Those regions upstream and downstream of exon 5 which we were able to confirm were used as homology arms (1942 bp upstream and 1241 bp downstream of our target sites) in the final construct. An Hr5/IE1 AmCyan K10 3’UTR cassette (AGG1036) was used to enable detection of the transgene by fluorescent microscopy. The multiplexed sgRNA expression cassette was synthesized (Genewiz) to contain an array of four cassettes each consisting of 600 bp upstream region of an endogenous *Ae. aegypti* pol III RNA (Ae U6.763 (AAEL017763), Ae U6.774 (AAEL017774), Ae U6.702 (AAEL017702), Ae 7SK (AAEL018514)) (37), an sgRNA targeting exon 5 of the *kmo* gene (cutting at 447, 468, 499, and 519 bp into exon 5 of *kmo*) (39), with one of four sgRNA backbone variants (23 with a 5 bp extended stem loop, 29, 9, 25) (40), and a poly-t (7 nt) terminator for the pol III promoter (Fig 1, S1). Three of these targets are previously validated (39) and the fourth was designed using CHOPCHOP and selected by location, closest off-targets for all sgRNAs as determined by CHOPCHOP are listed in Table S14. Complete primer sequences are listed in Table S11.

#### Identification of germline promoter, design and construction of *bgcn-Cas9* expression plasmid

Blastp using the *D. melanogaster* amino acid sequence was performed. The *Ae. aegypti* ortholog was identified (with 28% aa sequence identity) as AAEL004117, annotated as an ATP-dependent RNA helicase, consistent with the *D. melanogaster* gene annotation. The *bgcn*-Cas9 expression construct was built based on plasmids kindly provided by Omar Akbari, with several modifications (38) (Fig1b, S1). The fluorescent marker OpIE2-DsRED cassette was replaced with more easily visualized *AePUb-mCherry* (53). The human codon optimized *Streptococcus pyogenes-Cas9* was replaced with an insect codon optimized version (VectorNTI) synthesized by Genewiz. 5’ and 3’ RACE ready cDNA was generated from RNA extracted from ~30 pairs of ovaries or testes dissected from 5-7 days-post-emergence (dpe) Liverpool strain adults using Trizol (Invitrogen). Primers LA1076 then nested with LA1352 (5’) and LA1074 then nested LA1075 (3’) were used to amplify the 5’ and 3’ ends of the cDNA transcript and these amplicons were sequenced to verify the annotated UTRs (Fig S5, Table S11).

*Aebgcn* promoter and 3’UTR fragments were amplified using primers LA1725 and LA1726 (2213 bp upstream of ATG) and LA1737 and LA1738 (629 bp downstream of stop codon) (Table S11) from genomic DNA prepared from our Liverpool WT colony using the NucleoSpin Tissue kit (Macherey-Nagel) and ligated into the plasmid sequentially by standard restriction enzyme-based cloning to generate *bgcn*-Cas9 (AGG1207).

#### An improved *piggyBac* helper plasmid

Hyperactive piggyBac (53) has been used to increase the insertion efficiency in insects and so we synthesized (Genewiz) an *Ae. aegypti* codon optimized (ATGme) (54) version. This along with pGL3-PUb (gift from Zach Adelman, Addgene plasmid # 52891; http://n2t.net/addgene:52891;RRID:Addgene_52891) were digested with *Nco* I and *Fse* I and ligated using T4 DNA ligase (NEB) to generate AGG1245.

### Mosquitoes, transgenics and cage trial

#### Mosquito rearing

*Ae. aegypti* Liverpool strain (WT) was used for all experiments. All mosquitoes were reared under constant conditions: 28°C, 65-75% relative humidity and 14:10 light/dark cycle with 1 h of dawn and 1 h of dusk. Larvae were fed with ground TetraMin flake fish food (TetraMin) and adults were provided with 10% sucrose solution *ad libitum*. Females were blood fed with defibrinated horse blood (TCS Bioscience) using a Hemotek feeder (Hemotek Ltd) covered with Parafilm (Bemis).

#### Microinjections, crosses, screening

Transgenic *Ae. aegypti* mosquitoes were generated by microinjection of embryos less than 2 h post oviposition as described previously (55). G_0_ embryos were hatched one week after injection and larvae reared as described above. For the generation of the Cas9 line, embryos were injected with 500 ng/μl AGG1207 and 300 ng/μl AGG1245 (PUb hyperactive *piggyBac* transposase) in 1X injection buffer. For generation of *kmo*^sgRNAs^ transgenics, embryos were injected with 300 ng/μl Cas9 protein (PNABio), *in vitro* transcribed sgRNAs at 40 ng/μl sgRNA447, 40 ng/μl sgRNA519, and 300 ng/μl AGG1095 in 1x injection buffer.

Templates for *in vitro* transcription were designed as described previously (56) with LA925 (sgRNA447), LA926 (sgRNA519), LA927 (sgRNA499), LA928 (sgRNA468) and LA924 (common R) (Table S11). sgRNAs were *in vitro* transcribed using the MEGAscript T7 in vitro transcription kit (ThermoFisher) according to the manufacturers’ instructions. RNA was purified using the MEGAclear *in vitro* transcription reaction clean-up kit (ThermoFisher) aliquoted and stored at −80°C until use.

All G_0_ adults were crossed to WT mosquitoes. G_0_ males were crossed individually to 5 WT females for 2-3 days and then pooled to approximately 20 G_0_ individuals in a cage, while G_0_ females were crossed to WT males as a pool of approximately 20 G_0_ females to 20 WT males. G_1_ progeny were screened for presence of the fluorescent marker using a Leica MZ165FC microscope (Leica Biosystems).

#### Generation of white eyed mutant (*kmo^-/-^*) strain

To determine the rate of CRISPR/Cas9 induced cutting and germline homing, a *kmo^-/-^* knockout line was generated by crossing two white eyed non-drive inheriting individuals (one male and one female) generated from an inheritance assessment cross. The region encompassing the sgRNA recognition sites was amplified with primers LA1275 + LA518 and mutations identified by Sanger sequencing (Eurofins) and listed in Table S9. Deep sequencing of four replicates of *kmo^-/-^* adults (n = 24) indicates there are at least eight distinct *kmo* knockout alleles in the *kmo^-/-^* line even though the line was generated by crossing only two non-drive-inheriting founders (Table S9, Table S10). It is likely that the different mutant alleles in the germline of the founders were generated by nuclease activity originating from (i.e., deposited by) their trans-heterozygous parent.

#### Confirmation of insertion

Flanking PCRs (S2) were performed on *bgcn*-Cas9 transgenic lines according to previously reported methods (57, 58). gDNA was extracted from 10 individuals from pools *bgcn*-Cas9D using the NucleoSpin Tissue kit (Macherey-Nagel). DNA was digested with the restriction enzymes BamHI, MspI and NcoI (New England Biolabs) and PCRs were performed with DreamTaq (Thermo Fisher Scientific) and primers LA182, LA184, LA186 and LA187.

Five transgenic *kmo*^sgRNAs^ G_1_ males identified from the same pool (N) were individually outcrossed to five females each to obtain isolines. Genomic DNA was extracted from each founder male using the NucleoSpin Tissue kit (Macherey-Nagel) and subjected to two separate PCR reactions with primers LA2750, LA174 and LA1301, LA2755 to confirm correct homology-directed repair of the construct. PCR amplicons were further sequence confirmed by Sanger sequencing.

#### Phenotype data analysis

*kmo*^sgRNAs^ adult females and males (at least 20) were crossed to the opposite sex *bgcn*-Cas9 adults to generate trans-heterozygous adults (F_1_). For initial assessments of *kmo*^sgRNAs^ transgene inheritance, F1 adults were pooled into groups of at least 5 trans-heterozygous females or males and crossed to WT. For Line D, the majority of F_1_ trans-heterozygotes displayed a mosaic eyed phenotype, and these were crossed separately from their F_1_ siblings with a WT phenotype. Progeny (F_2_) were screened for presence of each transgene and eye color phenotype. We used a likelihood ratio test to compare rates of transgene inheritance compared to an expected distribution under standard Mendelian inheritance. We were able to interrogate additivity and total fit to compare the effects of insertion site, maternal/paternal inheritance of Cas9, mosaicism and replication on assessments of inheritance bias. Exponentiated log odds and standard errors were used to generate approximate 95% confidence intervals. This pooling approach allowed us to identify insertion lines demonstrating >50% inheritance of the *kmo*^sgRNAs^ transgene. However, this pooling approach does not take into account potential individual differences in fitness, mating rates or homing.

To account for these batch effects, and to accurately quantify rates of Cas9 cleavage in relation to homing, F_1_ trans-heterozygous females and males generated by crossing *kmo*^sgRNAs^ males to *bgcn*-Cas9 females were also crossed to a *kmo^-/-^* line. Crosses were performed as single pair crosses, and females were allowed to lay eggs individually. Progeny (F_2_) were screened as before for the presence of each transgene and eye color phenotype. Analyses for the proportion of F_2_ progeny with white eyes and *kmo*^sgRNAs^ inheritance were made by fitting a generalized linear mixed model, with a binomial (‘logit’ link) error distribution. This accounts for replication, and results in slightly different estimates from pooled data, with increased estimate intervals.

White-eyed progeny without the transgene were snap-frozen in liquid nitrogen and stored at −80°C for further analysis. Genomic DNA was extracted using the NucleoSpin Tissue kit (Macherey-Nagel). Further sequencing was carried out by Illumina MiSeq using primers LA4507, LA4508 flanking a 500 bp fragment including the sgRNA target sites following a previously published procedure and detailed below (59).

#### Cage trial

A cage trial was undertaken to study the performance of the split *bgcn*-Cas9/*kmo*^sgRNAs^ drive in a small laboratory population. A total of 12 cage populations were established (6 ratios in duplicate, designated 1 and 2 for each condition) with the following adults: experimental cages 100 *kmo*^sgRNAs^;*bgcn*-Cas9 trans-heterozygous females and 100 WT males (A1 and A2); control cages of 100 *kmo*^sgRNAs^ heterozygous females and 100 WT males (B1 and B2), 100 *kmo*^sgRNAs^ heterozygous females and 100 *kmo*^sgRNAs^ heterozygous males (C1 and C3), 100 heterozygous *bgcn*-Cas9 females and 100 WT males (D1 and D2), 100 heterozygous *bgcn*-Cas9 females and 100 heterozygous *bgcn*-Cas9 males (E1 and E2), and 100 WT females and 100 WT males (F1 and F2) (Fig 4a). Individuals destined for cages A, B, and C were derived from an initial cross of trans-heterozygous (*kmo*^sgRNAs^; *bgcn*-Cas9) males to WT females. Adults for *bgcn*-Cas9 only cages (D and E) were selected from a maintenance *bgcn*-Cas9 line (generation 9). WT adults for cages F were selected from the Liverpool mosquito line. The trans-heterozygous females used to establish cages A1-2 presented mosaic eyes, so the initial frequency for this eye phenotype in the experimental cages was 50%. To establish each generation of the cages, eggs were hatched in degassed reverse osmosis water and 250 L1 larvae/condition were randomly separated using a Biosorter (Union Biometrica). To keep all conditions as homogeneous as possible, larvae were reared in standardized trays with a set volume of water (2L) and following a feeding scheme described previously in (60). Pupae were separated by sex and females and males allowed to eclose separately in cages and provided with 10% sucrose *ad libitem*. Five days post eclosion all adults were anesthetized with CO_2_ and simultaneously transferred to the final cage (W24.5 x D24.5 x H24.5 cm) (BugDorm) so all the adults would have the same chances to mate. The trial was continued for six generations (Fig S6). At each generation, two ovipositions were collected from each cage and after the second oviposition the adults were snap-frozen and stored at −80°C for further molecular analysis. Screening for fluorescence and eye phenotype were performed at pupae stage.

#### Amplicon sequencing

Amplicon sequencing was carried out as previously published (59). gDNAs were extracted using the NucleoSpin Tissue kit (Macherey-Nagel). Approximately 500 bp surrounding the sgRNA target sites was amplified using primers listed in Table S11. A second round of PCR was performed using the Nextera XT index kit, and Nextera XT index kit D (Illumina). Amplicon sizes were verified on a Tapestation using the High Sensitivity D1000 Screentape (Agilent). The NEBNext Library Quant kit (NEB) was used to quantify the amplicons prior to pooling. Sequencing was carried out by the Bioinformatics, Sequencing and Proteomics facility at The Pirbright Institute.

### Data analysis, software, and packages

#### Amplicon-Seq analysis

The Illumina Miseq reads were first checked for sequence quality using FastQC (61). The low-quality regions and sequencing adapters were trimmed using the Trimmomatic tool (62). The reads were again subjected to quality control using FastQC (61) and further trimmed to a 200bp window (i.e., 100bp flanking either side of the sgRNA468 cut-site) using fastx_trimmer (63). Seqkit tool (64) was used to select the reads which are exactly 200bp in length and any reads less than that were removed. The selected reads were then aligned to the *kmo* reference sequence using BWA mem with default parameters (65). Any unmapped reads were removed. The obtained sam files were converted to bam file format, indexed, and sorted using the SAMtools (66). IGV viewer (67) was used for visualizing the sequence alignment file.

For the cutting assay samples, the variant information was extracted from the alignment file (sorted .bam file) using the tool pysamstats. The resulted stats file were read into a pandas (Python Library) data array and indel rate at each base location was calculated using the formula (59), variants [‘indel_rate’] = (variants [‘insertions’] + variants [‘deletions’]) /variants [‘reads_all’]). The identified indel rates were exported as CSV (comma-separated file) and plotted using Graphpad Prism 10.

For the analysis of cage trial samples, the libraries of *kmo*^sgRNAs^ containing amplicons and non *kmo*^sgRNAs^ containing amplicons were grouped separately and then analyzed. The sam files were obtained as above. The query sequence field in the sam file was used to identify the sequences which contain all four sgRNA sites (sgRNAs 447, 468, 499, and 519) including an intact PAM sequence. The number of wild-type reads in each individual dataset was calculated using SAMtools (66), exported as CSV and plotted using GraphPad Prism 10.

#### Phenotype data analysis

We carried out all phenotype analyses using R version 3.6.2 (R Development Core Team) (68). Data sets were summarized using ‘tidyverse’ (69) and figures generated using ‘ggplot2’ (70). Likelihood ratio tests carried out with ‘DescTools’ (71). Generalized linear mixed model analyses were carried out using ‘lme4’ (72), and summarized with ‘emmeans’ (73) and ‘sjPlot’ (74), model residuals were checked for violations of assumptions using the ‘DHARMa’ package (https://github.com/Philip-Leftwich/Population-level-demonstration-of-multiplex-drive-Aedes-aegypti) (75).

#### Mathematical modeling

Complete details of the model available in Supplementary information S4. All Matlab scripts used in the mathematical modeling are freely available via the Open Science Framework (osf.io/bp4yh).

## Data availability

All raw reads from amplicon-Seq were submitted to NCBI SRA with the accession number PRJNA741076. All scripts used are provided in Supplemental S3. Any intermediate analysis files will be provided on request.

## Author Contributions

MAEA, THS, PTL, KE and LA designed the research. MAEA, EG, DKP, JXDA, LS and KN performed the research. MAEA, THS, PTL, SANV contributed reagents. DKP, JA, PTL, MPE contributed analytic tools and analyzed the data. All authors contributed to writing and editing the paper.

## Acknowledgments

MAEA, EG, JXDA, LS, KN, SANV, and LA were funded through a Defense Advanced Research Projects Agency (DARPA) award [N66001-17-2-4054] to Kevin Esvelt at MIT. MPE and PTL were supported by the Wellcome Trust [110117/Z/15/Z]. For the purpose of Open Access, the author has applied a CC BY public copyright license to any Author Accepted Manuscript (AAM) version arising from this submission. LA and THS were funded by the UK Biotechnology and Biological Sciences Research Council [BBS/E/I/00007033, BBS/E/I/00007038, and BBS/E/I/00007039 to The Pirbright Institute]. DKP’s PhD studentship was funded by The Pirbright Institute. The views, opinions and/or findings expressed are those of the authors and should not be interpreted as representing the official views or policies of the U.S. Government. The funders had no role in study design, data collection and analysis, decision to publish, or preparation of the manuscript.

We would like to thank Graham Freimanis at the Bioinformatics, Sequencing and Proteomics core facility for running the Illumina MiSeq, and advice and consultations with regards to the data. We would also like to thank Rebecka Ireland, Jessica Mavica, and Sophia Fochler for their assistance in the Insectary in the early stages of this project.

