## Supplemental information for "A multiplexed, confinable CRISPR/Cas9 gene drive propagates in caged *Aedes aegypti* populations"

**This PDF file includes:**

Supplementary text S1 to S4
Figures S1 to S7
Tables S1 to S14
Legends for Dataset S1

**Other supplementary materials for this manuscript include the following:**

Dataset S1 Cage trial

**S1. Complete plasmid sequences**

>AGG1095

```
FEATURES             Location/Qualifiers
     misc_RNA        7336..7417
                     /label="sgRNA_backbone_29"
     intron          1966..2069
                     /label="Intron 2-3"
     Terminator      5289..6070
                     /label="K10\3'UTR"
     misc_feature    5232..5270
                     /label="NLS"
     misc_feature    10013..10015
                     /label="PAM"
     Promoter        complement(545..649)
                     /label="AmpR promoter-006_"
     misc_feature    6772..7314
                     /label="U6-2"
     misc_RNA        9993..10012
                     /label="kmo447"
     misc_RNA        1160..1179
                     /label="kmo447"
     misc_feature    7978..8057
                     /label="sgRNA_backbone-09"
     misc_RNA        7316..7335
                     /label="kmo-519"
     CDS             complement(983..1051)
                     /label="LacZ alpha"
     misc_feature    10364..11039
                     /label="pUC Origin"
     misc_feature    complement(join(11187..11458,1..544))
                     /label="KanR"
     terminator      8761..8767
                     /label="pol_III\terminator"
     protein_bind_site 10040..10062
                     /label="lac operator"
     misc_RNA        6659..6678
                     /label="kmo447"
     misc_feature    10016..10088
                     /label="LacZ Alpha"
     terminator      8058..8090
                     /label="pol_III\terminator"
     CDS             4520..5220
                     /label="AmCyan"
     exon            2578..2665
                     /label="Exon 4"
     misc_feature    6071..6658
                     /label="U6-1"
     Promoter        3133..4334
                     /label="Hr5-IE1\promoter"
     terminator      7418..7424
```

/label="pol_III\terminator"
misc_feature 6679..6764
/label="sgRNA_backbone-23"
misc_feature 1189..3130
/label="HA 5'"
promoter complement(10066..10096)
/label="lac promoter"
intron 2666..2724
/label="Intron 4-5"
misc_feature 8773..9992
/label="HA 3'"
Promoter complement(545..643)
/label="AmpR promoter-008_"
misc_feature 8681..8760
/label="sgRNA_backbone"
misc_feature 930..1159
/label="LacZ Alpha"
misc_RNA 7960..7977
/label="kmo-449"
misc_RNA 8663..8680
/label="kmo-468"
misc_feature 7425..7959
/label="U6-3"
terminator 6765..6771
/label="pol_III\terminator"
misc_signal 4469..4478
/label="Kozak sequence-002_"
exon 2070..2312
/label="Exon 3"
exon 1896..1965
/label="Ex2"
misc_feature 8091..8660
/label="7SK promoter"
CDS 4524..5224
/label="AmCyan"
misc_feature 4488..4514
/label="NLS"
intron 4383..4452
/label="adh"
intron 2313..2577
/label="Intron 3-4"

ORIGIN
1 tttctttcca gacttgttca acaggccagc cattacgctc gtcatcaaaa tcactcgcat
61 caaccaaacc gttattcatt cgtgattgcg cctgagcgag acgaaatacg cgatcgctgt
121 taaaaggaca attacaaaca ggaatcgaat gcaaccggcg caggaacact gccagcgcat
181 caacaatatt ttcacctgaa tcaggatatt cttctaatac ctggaatgct gttttcccag
241 ggatcgcagt ggtgagtaac catgcatcat caggagtacg gataaaatgc ttgatggtcg
301 gaagaggcat aaattccgtc agccagttta gtctgaccat ctcatctgta acatcattgg
361 caacgctacc tttgccatgt ttcagaaaca actctggcgc atcgggcttc ccatacaatc
421 gatagattgt cgcacctgat tgcccgacat tatcgcgagc ccatttatac ccatataaat
481 cagcatccat gttggaattt aatcgcggcc tagagcaaga cgtttcccgt tgaatatggc

541 tcatactctt cctttttcaa tattattgaa gcatttatca gggttattgt ctcatgagcg
601 gatacatatt tgaatgtatt tagaaaaata aacaaatagg ggttccgcgc acatttcccc
661 gaaaagtgcc acctgacgtc taagaaacca ttattatcat gacattaacc tataaaaata
721 ggcgtatcac gaggcccttt cgtctcgcgc gtttcggtga tgacggtgaa aacctctgac
781 acatgcagct cccggagacg gtcacagctt gtctgtaagc ggatgccggg agcagacaag
841 cccgtcaggg cgcgtcagcg ggtgttggcg ggtgtcgggg ctggcttaac tatgcggcat
901 cagagcagat tgtactgaga gtgcaccata tgcggtgtga aataccgcac agatgcgtaa
961 ggagaaaata ccgcatcagg cgccattcgc cattcaggct gcgcaactgt tgggaagggc
1021 gatcggtgcg ggcctcttcg ctattacgcc agctggcgaa agggggatgt gctgcaaggc
1081 gattaagttg ggtaacgcca gggttttccc agtcacgacg ttgtaaaacg acggccagtg
1141 aattgacgcg tattgggatg ccatataatg tgggcggcac gggggcccgg cgtagtagat
1201 gttacctgtg gcggtagggt ttttttcca tgtttagtga aaacggaaga agaagagacg
1261 ttctaagagt ttttgttttc caagactgtg ttcaagtagg attaccaccg ctagctttcg
1321 tcaacgggtc caacgaaaaa tctaaggcac gggccttaac aaggatcgta aagggttcaa
1381 tagtggaaaa atagtactga gaaatactat ttgttaagca ctgaaaagta ctgtttttgt
1441 agcattgaga agtactgtag ctttaggtag tttttaaaaa ctttcaaaag caaagagaat
1501 tcgtgtacac acagcacgaa ggtacgatgc gcactgacat tcgttgtagg actcgatttt
1561 tacagcactt gtcgtaatta tccaacgcag tgatcctcgt tgcaacttat ttgtaaacta
1621 gtattacaaa ataattttat ttgacgaaca aatttaaaat ttgagtagta cgtggaatct
1681 ctaaaaaata ctctagtata cacagattga gttatttcat aagtcaatac atatttgtcc
1741 aaattttata tcaaatcgta taaaatcatg ggtaggacga atgtcttact cagcattttt
1801 tgtgcgtagg acatatcctt tttgtcctac ccacttctcg cgccactggt aacgacatag
1861 aaaatctaat attgttctaa cattctgctt tcaaggttgg ctctctattt gcactccacc
1921 tcggcaaaaa aggacacact gtggacctgt atgagtacag agaaggtact tgaacattct
1981 ttccgaaaaa cacttccgcc aatttgtcgc gttcgtcatt caaccattca ttgcgttcac
2041 taatccctca tgcttcctac atttcgcaga cattcgcacg gccgaactgg tcattggtcg
2101 tagtatcaac ttggcgctgt ctgcccgagg ccgcaaagca ctggccgagg tgggcctgga
2161 ggacgccctc ctccagcatg gcattccaat gaagggccgc atgttacacg atctgaaggg
2221 aaaccgtaag attgtgccgt acgatgccaa caccaaccaa tgcatctact cggtggggcg
2281 aaaacatctg aacgaggtgc tgctcgacgg tgagtaaggc gtgtggtttc tggagaaatt
2341 cgatcggtat cggttacaaa gtagcgctgt tctggtgaac ggattatgct actgaagcct
2401 acacgtgtga tgtgtatttg tgcgttttgt tctgaatggg catgacggac atttggcggg
2461 cttcactcac tacatttcat tgcgtagcaa gtttaccttg ggcgaacggg aaacatgaga
2521 aatttttatt ctatgttatg taaaaggcta actgataatg ttgtttattt tagcggccga
2581 gaagtacccg aacattcatc tttatttcaa caaaaaactc caatcagcta atctggatga
2641 gggagagatg agtttcattg agtgagtaca cttaattggt ttccttgtga caagcgttat
2701 caccttcccc cgaatttcag tccaacgacg aaggaatcta ctcacaccaa ggccgatctg
2761 atcgtgggct gcgatggagc ttacagtgct gttcgcaaag aaatcgtcaa acggccgggt
2821 tatgactaca gtcagacgta catcgaacat ggctatctgg agctgtgtat tcctccgacc
2881 aaggatgggg atttcgcgat gcctcacaac tatttgcaca tttggccccg gggaaagttt
2941 atgatgatcg ccctgcccaa tcaggatcgc acttggacgg tgacgctgtt catgccgttc
3001 accaacttca acagtattaa gtgcgatggc gatttgttga agttcttccg gacatacttc
3061 cccgatgcga ttgatctgat tggtcgtgag cggttggtta aggatttctt taagaccagg
3121 cctcaaccta gggctttacg agtagaattc tacgcgtaaa acacaatcaa gtatgagtca
3181 taatctgatg tcatgttttg tacacggctc ataaccgaac tggctttacg agtagaattc
3241 tacttgtaat gcacgatcag tggatgatgt catttgtttt tcaaatcgag atgatgtcat
3301 gttttgcaca cggctcataa actcgcttta cgagtagaat tctacgtgta acgcacgatc
3361 gattgatgag tcatttgttt tgcaatatga tatcatacaa tatgactcat ttgtttttca
3421 aaaccgaact tgatttacgg gtagaattct acttgtaaag cacaatcaaa aagatgatgt
3481 catttgtttt tcaaaactga actcgcttta cgagtagaat tctacgtgta aaacacaatc
3541 aagaaatgat gtcatttgtt ataaaaataa aagctgatgt catgttttgc acatggctca
3601 taactaaact cgctttacgg gtagaattct acgcgtaaaa catgattgat aattaaataa

3661 ttcatttgca agctatacgt taaatcaaac ggacgctcga ggttgcacaa cactattatc
3721 gatttgcagt tcgggacata aatgtttaaa tatatcgatg tctttgtgat gcgcgcgaca
3781 tttttgtagg ttattgataa aatgaacgga tacgttgccc gacattatca ttaaatcctt
3841 ggcgtagaat ttgtcgggtc cattgtccgt gtgcgctagt agcatgcccg taacggacct
3901 cgtacttttg gcttcaaagg ttttgcgcac agacaaaatg tgccacactt gcagctctgc
3961 atgtgtgcgc gttaccacaa atcccaacgg cgcagtgtac ttgttgtatg caaataaatc
4021 tcgataaagg cgcggcgcgc gaatgcagct gatcacgtac gctcctcgtg ttccgttcaa
4081 ggacggtgtt atcgacctca gattaatgtt tatcggccga ctgttttcgt atccgctcac
4141 caaacgcgtt tttgcattaa cattgtatgt cggcggatgt tctatatcta atttgaataa
4201 ataaacgata accgcgttgg ttttagaggg cataataaaa gaaatattgt tatcgtgttc
4261 gccattaggg cagtataaat tgacgttcat gttggatatt gtttcagttg caagttgaca
4321 ctggcggcga caagcaattg gtacccgggt aggatcctag tgaattccta atctggcggt
4381 aagttgatca aaggaaacgc aaagttttca agaaaaaaca aaactaattt gatttataac
4441 acctttagaa agcgaagttg agattcaggc caccatggga gatcccaccc cacccaagaa
4501 gaagcgcaaa gctagcgtta tggccctgtc caacaagttc atcggcgacg acatgaagat
4561 gacctaccac atggacggct gcgtgaacgg ccactacttc accgtgaagg gcgagggcag
4621 cggcaagccc tacgagggca cccagacctc caccttcaaa gtcacaatgg ccaacggcgg
4681 ccccctggcc ttctccttcg acatcctgtc caccgtgttc atgtacggca accgctgctt
4741 caccgcctac cccaccagca tgcccgacta cttcaagcag gccttccccg acggcatgtc
4801 ctacgagaga accttcacct acgaggacgg cggcgtggcc accgccagct gggagatcag
4861 cctgaagggc aactgcttcg agcacaagtc caccttccac ggcgtgaact tccccgccga
4921 cggccccgtg atggccaaga agaccaccgg ctgggacccc tccttcgaga agatgaccgt
4981 gtgcgacggc atcttgaagg gcgacgtgac cgccttcctg atgctgcaag gcggcggcaa
5041 ctacagatgc cagttccaca cctcctacaa gaccaagaag cccgtgacca tgccccccaa
5101 ccacgtggtg gagcaccgca tcgccagaac cgacctggac aagggcggca acagcgtgca
5161 gctgaccgag cacgccgtgg cccacatcac ctccgtggtg cccttctccg gactccgctc
5221 ccagatctcc cgacccaaga aaaagcggaa ggtggaggac ccgtaagatc caccggatct
5281 agataactgg agcttgataa cattatacct aaacccatgg tcaagagtaa acatttctgc
5341 ctttgaagtt gagaacacaa ttaagcatcc cctggttaaa cctgacattc atacttgtta
5401 atagcgccat aaacatagca ccaatttcga agaaatcagt taaaagcaat tagcaattag
5461 caattagcaa taactctgct gacttcaaaa cgagaagagt tgcaagtatt tgtaaggcac
5521 agtttataga ccaccgacgg ctcattaggg ctcgtcatgt aactaagcgc ggtgaaaccc
5581 aattgaacat atagtggaat tattattatc aatggggaag atttaaccct caggtagcaa
5641 agtaatttaa ttgcaaatag agagtcctaa gactaaataa tatatttaaa aatctggccc
5701 tttgaccttg cttgtcaggt gcatttgggt tcaatcgtaa gttgcttcta tataaacact
5761 ttccccatcc ccgcaataat gaagaatacc gcagaataaa gagagatttg caacaaaaaa
5821 taaaggcatt gcgaaaactt tttatggggg atcattacac tcgggcctac ggttacaatt
5881 cccagccact taagcgacaa gtttggccaa caatccatct aatagctaat agcgcaatca
5941 ctggtaatcg caagagtata taggcaatag aacccatgga tttgaccaaa ggtaaccgag
6001 acaatggaga agcaagagga tttcaaactg aacacccaca gtactgtgta ctaccactgg
6061 cgcgtttggg ggccggccgt ttccagactt tccctcccgg taaacggaga caaaacgaca
6121 gacgtaagta ggtacatatg cataccgcac ggacaaatca aatttgtctg gcagctccaa
6181 ttagagtcgt taaaaattta acgatgcgct aaataacttc aagctatttg tctcgctgga
6241 ttggcttcga gtggtaagat cctatcaaat gccgaaaaca aaaaactttc ttcttaattg
6301 ttcgttcttc aacacctctc catggtgata acggatacgg tttcattgtc agcatccatc
6361 ctccgaaaaa tacattacgc cttgaaatat gcaatcgcaa acacggatct gtttggaaca
6421 tttattttac tatgaagaga tgcgataggt aatatttatt tgagcgttta agatactcat
6481 tgttctctca aagaatgtca ttgaaagcca acgaggtcaa atcaaatatt ataataaaaa
6541 ggtcaaagag gactaactta aagctctctt tatggatagg aaaaaatatt ttcgcccatc
6601 gctagaactt ttaccgtttc cattgagtat ataactaaga tgaatgaggc taattgatgc
6661 catataatgt gggcggcagt tccagagtcg tgctgggaac agcacgacaa gttggaataa
6721 ggcaagtccg ttatcatgcc ggaaggcagg caccgattcg gtgctttttt tggaacccga

6781 atttagtgct atataattta attccactag agtttgtatc ctttgataga tacgcgtatt
6841 tcgacctcaa ctgcaaggcc gtcgtgtact agacttgact aatccagact ggtcttttag
6901 ttatgacttc tgtccacatc tccatacatt caacgcactg tgcggctgtg ctgtgcgact
6961 ccgtcgagtc gaccaacata gttgaaacaa attgaatatt taattgatcg ttataggaat
7021 ggtgttagat gagtcatcct ttacagtaag cacatacagt attataattg aagatcgtcg
7081 gcagataggt gtgtagggta gagtatcagc aataagttgg gacgtttgac tttttgtagg
7141 tagacaaaaa ctaaactttt tttcgcttct ctatgtgtgc ccccccgggt agcgtatcgt
7201 tccgattgtg gtgcgaacga atgaaatcgc ccatcgagtt gatacgtcca tccatcgcta
7261 gaaccgcgtt cgctgtagaa gactatataa gagcagaggc aagagtagtg aaatGcacag
7321 tacaatcctc gaatcgtcgc agagcatctg aaaagatgca agttgcgata aggcaagtcc
7381 gttatcaagc tcgggagagc tggcaccgag tcggtgcttt ttttgagtct ggtaacccta
7441 gtgcacgcaa atatctcgcg ggcatatttg gttgctgagg tatatttata tttgaacgcc
7501 atgagaaaaa gcggaagaaa ttggctcatg gccgatttta aggatattta aaaattgtac
7561 aatgtacata taataggcca ggagaagtgg atgaacttgt cattcatttt tctgtcaatt
7621 ctcatacaaa atctactttt tctttgacat aaattcactc taggtgaacc acttcccctg
7681 gcctattaaa catccgttcc ttcaatgtgt tcttttttttt aagcgtgtgt taaaagtttg
7741 ctctgctggt gaattcacgc tctacccgtt caggcagcat tcatcgaaaa gccctatctg
7801 ctcgcacaca tttacaaaat gctgattgcg ttgtgtgctg aatgggtcac tcgtccgtca
7861 ctgcttgctg tgtacactgt acagttacgc agtctgtgca tcgctagaat catatttacg
7921 gaagagtatt atatataccc gatgcgttgc tcttcgattg gttcccttct acgggcagtc
7981 ctagagccat gaaaatggca agttaggata aggctagtcc gtattcaacg ctgaaaagcg
8041 tggcaccgag tcggtgcttt ttttgggaaa cccgaaacag attttatttt atgctccatt
8101 ctccgccact tgttgatgcg gaccctaacc acgtggtcgc tcctctgctc accggagcac
8161 gtttcataca gcctgacgac gacgagcaat cagaggtatg gtgagcatgc gcatggagag
8221 tggacagcag tgcaccctaa aatcaattca cacatcatgt gtcaatagct gtgtcaatgt
8281 tgcacagcct tttcttatta aatttactcc ttttgtgacc atttctcttt catccaccgt
8341 tattttaatg agttttgtgt tccggtggac gaacgttcac acaaaaaatg tgtaaatctt
8401 aatcaaccag aacacaaagt atagtgaaaa aattaaagtg tgtggctttt atacatccta
8461 actgtaaatt atttttagag tgcgtgcgat cgttctctcg aaccacgctc tccgctacac
8521 attcgcagcg aatggcgtga atggatgaaa gaacaaacta aagtttattt ttagattcgt
8581 ctcaaaacaa ctgctgtgca tcgctagaac caagaaatac gccactcagt atatatagca
8641 cttccaaccc cgctttcctc ggcggtgatc attggtgatg gttgcagaga cacgggagtg
8701 tcaagttgca ataaggccag tccgttatca gacgtgggaa cgtcggcacc gattcggtgc
8761 ttttttgcc gcggagttgt tcaatcaaca tggcagtgac gttgatagga tactggctga
8821 gtttagtgat acgcgttggg aggatgcaca ctctatctgc gatctggcca tgtataatta
8881 tgttgaggtt agtatatggt cttttattta tatcgtacgt tttgtatgcg gtcgttttgt
8941 aggtaccgta aattcgggtg aaattgatca gtagggtgaa attgatcact gtgtcacacg
9001 attttatttc cctctaatag agcacagaaa ccaatgcaac ttatgtaaat gaacgttgtt
9061 tgtcttaact attgttaaat tgcatgttgt gaagcttttt gtgttatgga atatttattt
9121 caattaaaat aataggaaat tccatcatcg tgttctgtgt ggtactggca gacatcaata
9181 aacttatagt ttcaaacaag gatttggaca tggtgtcaac ctataaattt gcaaggatgc
9241 ttgaaatata tccccaaaac gaattttatc atcaaaatac gtaccaattt gttcattttg
9301 ttgtaataat tgaatttgaa ttgtatagat atcatcaaat tccttcggga attgcctaca
9361 ttcaggcgtt ttctgcggta ttgctgaaaa ttaattgtta attatttcca taaattttgc
9421 attgttaaga atttggcaag catatttgga ttcaggaggc tcaaattaaa taagtaaagt
9481 tgtttttaaa gctaacaata attgttttaa caggtgatca atttcacctc gaaatggaat
9541 tccttgattt tttattttag agacgatttt cagcactaaa atcacaactg tatgaaaatt
9601 tgagtatata aaccaatgaa gctcaccgtc gtacttgttt tcctgcatta agttgtttgg
9661 aattggcaac ataatagaaa acagacatgg gaaacagtga aaagtgatca atttcacccg
9721 aaattacggt atacgcagaa ttaaaatgac aagaagttcc accgcttctt tgatatccta
9781 tgaacattat aacattgact gcaatttgag tgacttcagt aatttcactg gtaacgtaaa
9841 gtgggggcaa attgatcact ggggtgaatt taatcaggtc ggtaccatat agcatctctt

9901 ccaagaatgc ttaatgtttc attggattca cagacattgc atgttttcta gtttatagat
9961 gtccaatgat gattttccaa taggggcgcg ccgccatata atgtgggcgg cacggtggtc
10021 atagctgttt cctgtgtgaa attgttatcc gctcacaatt ccacacaaca tacgagccgg
10081 aagcataaag tgtaaagcct ggggtgccta atgagtgagc taactcacat taattgcgtt
10141 gcgctcactg cccgctttcc agtcgggaaa cctgtcgtgc cagctgcatt aatgaatcgg
10201 ccaacgcgcg gggagaggcg gtttgcgtat tgggcgctct tccgcttcct cgctcactga
10261 ctcgctgcgc tcggtcgttc ggctgcggcg agcggtatca gctcactcaa aggcggtaat
10321 acggttatcc acagaatcag gggataacgc aggaaagaac atgtgagcaa aaggccagca
10381 aaaggccagg aaccgtaaaa aggccgcgtt gctggcgttt ttccataggc tccgcccccc
10441 tgacgagcat cacaaaatca caaaaatcga cgctcaagtc agaggtggcg aaacccgaca
10501 ggactataaa gataccaggc gtttccccct ggaagctccc tcgtgcgctc tcctgttccg
10561 accctgccgc ttaccggata cctgtccgcc tttctccctt cgggaagcgt ggcgctttct
10621 catagctcac gctgtaggta tctcagttcg gtgtaggtcg ttcgctccaa gctgggctgt
10681 gtgcacgaac cccccgttca gcccgaccgc tgcgccttat ccggtaacta tcgtcttgag
10741 tccaacccgg taagacacga cttatcgcca ctggcagcag ccactggtaa caggattagc
10801 agagcgaggt atgtaggcgg tgctacagag ttcttgaagt ggtggcctaa ctacggctac
10861 actagaagaa cagtatttgg tatctgcgct ctgctgaagc cagttacctt cggaaaaaga
10921 gttggtagct cttgatccgg caaacaaacc accgctggta gcggtggttt ttttgtttgc
10981 aagcagcaga ttacgcgcag aaaaaaagga tctcaagaag atcctttgat cttttctacg
11041 gggtctgacg ctcagtggaa cgaaaactca cgttaaggga ttttggtcat gagattatca
11101 aaaaggatct tcacctagat ccttttaaat taaaaatgaa gttttaaatc aatctaaagt
11161 atatatgagt aaacttggtc tgacagttag aaaaactcat cgagcatcaa atgaaactgc
11221 aatttattca tatcaggatt atcaatacca tatttttgaa aaagccgttt ctgtaatgaa
11281 ggagaaaact caccgaggca gttccatagg atggcaagat cctggtatcg gtctgcgatt
11341 ccgactcgtc caacatcaat acaacctatt aatttcccct cgtcaaaaat aaggttatca
11401 agtgagaaat caccatgagt gacgactgaa tccggtgaga atggcaaaag tttatgca
//

>AGG1207

FEATURES Location/Qualifiers
rep_origin complement(14461..15143)
/label="ColE1 origin"
Promoter complement(16099..16203)
/label="AmpR promoter-006_"
misc_recomb 13211..13911
/label="PiggyBacL"
CDS complement(15238..16098)
/label="AmpR"
protein_bind_site 14099..14121
/label="lac operator"
misc_feature 8751..8804
/label="T2A"
CDS complement(59..127)
/label="LacZ alpha"
misc_feature 9525..10177
/label="AeBCGN 3'UTR"
protein_bind_site 14170..14191
/label="CAP binding site"
misc_feature 4602..8702
/label="aCas9"

```
     old_sequence      13912..14040
                       /label="flanking seq"
     Promoter          10863..12253
                       /label="Ae PUb Promoter"
     misc_binding      1979..2263
                       /label="attB site"
     misc_binding      4569..4594
                       /label="NLS"
     CDS               12257..12961
                       /label="a_mCherry"
     terminator        complement(12965..13193)
                       /label="SV40"
     old_sequence      225..932
                       /label="flanking sequence"
     misc_feature      8732..8750
                       /label="NLS"
     promoter          complement(14125..14155)
                       /label="lac promoter"
     misc_recomb       933..1930
                       /label="PiggyBacR"
     ORF               8805..9524
                       /label="GFP"
     misc_feature      10178..10854
                       /label="p10 3'UTR"
     misc_feature      2266..4485
                       /label="AeBCGN pro"

ORIGIN
        1 accatatgcg gtgtgaaata ccgcacagat gcgtaaggag aaaataccgc atcaggcgcc
       61 attcgccatt caggctgcgc aactgttggg aagggcgatc ggtgcgggcc tcttcgctat
      121 tacgccagct ggcgaaaggg ggatgtgctg caaggcgatt aagttgggta acgccagggt
      181 tttcccagtc acgacgttgt aaaacgacgg ccagtgccaa gctttgttta aaatataaca
      241 aaattgtgat cccacaaaat gaagtggggc aaaatcaaat aattaactag tgtccgtaaa
      301 cttgttggtc ttcaactttt tgaggaacac gttggacggc aaatcgtgac tataacacaa
      361 gttgatttaa taattttagc caacacgtcg ggctgcgtgt tttttgcgct ctgtgtacac
      421 gttgattaac tggtcgatta aataatttaa tttttggttc ttctttaaat ctgtgatgaa
      481 attttttaaa ataactttaa attcttcatt ggtaaaaaat gccacgtttt gcaacttgtg
      541 agggtctaat atgaggtcaa actcagtagg agttttatcc aaaaaagaaa acatgattac
      601 gtctgtacac gaacgcgtat taacgcagag tgcaaagtat aagagggtta aaaaatatat
      661 tttacgcacc atatacgcat cgggttgata tcgttaatat ggatcaattt gaacagttga
      721 ttaacgtgtc tctgctcaag tctttgatca aaacgcaaat cgacgaaaat gtgtcggaca
      781 atatcaagtc gatgagcgaa aaactaaaaa ggctagaata cgacaatctc acagacagcg
      841 ttgagatata cggtattcac gacagcaggc tgaataataa aaaaattaga aactattatt
      901 taaccctaga aagataatca tattgtgacg tacgttaaag ataatcatgc gtaaaattga
      961 cgcatgtgtt ttatcggtct gtatatcgag gtttatttat taatttgaat agatattaag
     1021 ttttattata tttacactta catactaata ataaattcaa caaacaattt atttatgttt
     1081 atttatttat taaaaaaaaa caaaaactca aaatttcttc tataaagtaa caaaactttt
     1141 aaacattctc tcttttacaa aaataaactt attttgtact ttaaaaacag tcatgttgta
     1201 ttataaaata agtaattagc ttaacttata cataatagaa acaaattata cttattagtc
     1261 agtcagaaac aactttggca catatcaata ttatgctctc gacaaataac ttttttgcat
     1321 tttttgcacg atgcatttgc ctttcgcctt attttagagg ggcagtaagt acagtaagta
     1381 cgtttttca ttactggctc ttcagtactg tcatctgatg taccaggcac ttcatttggc
```

1441 aaaatattag agatattatc gcgcaaatat ctcttcaaag taggagcttc taaacgctta
1501 cgcataaacg atgacgtcag gctcatgtaa aggtttctca taaatttttt gcgactttga
1561 accttttctc ccttgctact gacattatgg ctgtatataa taaaagaatt tatgcaggca
1621 atgtttatca ttccgtacaa taatgccata ggccacctat tcgtcttcct actgcaggtc
1681 atcacagaac acatttggtc tagcgtgtcc actccgcctt tagtttgatt ataatacata
1741 accatttgcg gtttaccggt actttcgttg atagaagcat cctcatcaca agatgataat
1801 aagtatacca tcttagctgg cttcggttta tatgagacga gagtaagggg tccgtcaaaa
1861 caaaacatcg atgttcccac tggcctggag cgactgtttt tcagtacttc cggtatctcg
1921 cgtttgtttg atcgcacggt tcccacaatg gttaattcga gctcgcccgg ggtcctaggt
1981 cgacgatgta ggtcacggtc tcgaagccgc ggtgcgggtg ccagggcgtg cccttgggct
2041 ccccgggcgc gtactccacc tcacccatct ggtccatcat gatgaacggg tcgaggtggc
2101 ggtagttgat cccggcgaac gcgcggcgca ccgggaagcc ctcgccctcg aaaccgctgg
2161 gcgcggtggt cacggtgagc acgggacgtg cgacggcgtc ggcgggtgcg gatacgcggg
2221 gcagcgtcag cgggttctcg acggtcacgg cgggcatgtc gacgcggccg cCAACGTTGG
2281 ggcgtcataa gccttaaaaa aatcagaggt ttcatacaaa aaacaagggg ttgaaaaact
2341 tgaaatttag cgtcagtgtg taccatccca tagaggagga tcttctcttt atagtagcat
2401 tgctgaataa cttgtaaatc cattcgtttg gtcagtaaca gcgtactaca cacttgctta
2461 ccaatggctt ttagggcgag attcataagt taagcgacat gttatacaat atcttgcata
2521 acttataatt ccccctaaaa gccattgact aaaacaaaat cttaaattta aagattagag
2581 cactgcttag tcttcgaaat atgtgttctt caaaatttga ggtgtggttt caatccacac
2641 accaacaatt ttgagtgtgg ctcgttgatt tcaagcaaat gaatggagca aataaaaact
2701 atcatcactg ggggatagag caggatattc tcttcatcgt tgcaatatta aataacatgt
2761 aaatccatta gtttgctcag taacaacaag ctacacagtt agggggagat cataaattgg
2821 cttgtttagg ggtatcctgt ttttcacata ttctgattct acgttaaaaa ctaaaccaga
2881 atccaaagtt tcataattat ccatgctctg gaaccaaaga caaattaaag acctccatgt
2941 cccttgcagt tgctcaaaat tttatggctc aaatggtaat ctagaatgcc gtgaaaagtg
3001 tactgcgatc tgtcaaactt gggtaccttt tgtactaccg agggacgaaa tgcagtacta
3061 gaagtggtac ccactctgag cccgagtgtg acggacctgc tggaactatg tttaaaacac
3121 gtttgaaatt agtatggaaa tcgcgtttga atcacccctc ggctaatttt acttcggcac
3181 gaactgtcat tgtgcaaatt gaaatgtcat taagtcgacg gattgaaatt ttttttattc
3241 atttattata tccaaataaa gatattgaag cgcaacaaaa tcgtgttaca cctagaaaaa
3301 tgtgctcttt cgatcaagct acaaaaaact gtgaatttca tttcatcact aaacaaccca
3361 attgggtcct aaaattttta atgaaatctt gtttttattt aataacacga aaaagcatgt
3421 tactgtaact actttagcaa tttttcccgc tcaaataatg gctatatcat gtttaaactt
3481 taatttaaaa atttggtcca taaatgaacc acagacaaac agacgtcaca ctctcatcat
3541 tgtccatcga ccaccttttt aacggtcgat tcaaaaacat ggtaggtggc caatccgcca
3601 cccgcagcgc tcgcatcgtt tttgttcgca tttgacgttt acacactacc gccatctgtt
3661 gacctgtcgg ccagacacgc ttattttagc attgggcgta catgttctca tgactataat
3721 tttgatcgag atttgttcta agtgttacgt ctgtttgtct gtgaatgaac cttgacactt
3781 tggatcatgt ttgacgttcg cttagtcgac aaaaacacca caggggtttt agttcgacca
3841 ctggggttgt tcctatctga catttcgtaa gggacacgga aaacaaaata cacctaaaat
3901 ttgagtttag gccaaaggat gtgacaaaat ctaaataaat gtttttttagg cttaaaccaa
3961 cgaaaaacat tagaaaattg agtaaacatg tgtttttggc ctaaacttaa gcgtttggca
4021 cttaaattga aacatggctt taggacccta ttaagcgaga tataaacaga ttatccgcga
4081 aacaatttat tttgaaaaat gctccgcgag tcaaaaagtc tggaaacccc ttgcatccgt
4141 gagtgcctct tatgacgttg cgtccgtcgt gcagtgagtg gttgttaggt tttcagcttc
4201 ggttcgtttc gtcgtccgtc caacacactt cttgacagtc gagcgaaaaa tttggtgggc
4261 tctccttcgc aaaggtgcaa aaatcgtcat ctgcgtttcg aatggtcaat tttgctaccg
4321 tacgcatcct ggaacaggac aatttctagc acattatctg ctgcaccgat tccgccagca
4381 gccgaggtaa gtttcagtat ggaaaaccag caggacaggg cggaaaattg ataaaatatt
4441 tttgcatttt cgcgaaccga aaaaaaaacg aaaacagctc caacctcgag atggactata
4501 aggaccacga cggagactac aaggatcatg atattgatta caaagacgat gacgataaga

4561 tgggagatcc caccccaccc aagaagaagc gcaaagctag cgataagaag tactcgatcg
4621 gactggatat cggaaccaac tccgtgggat gggccgtgat caccgatgaa tacaaggtgc
4681 catcgaagaa gttcaaggtg ctgggaaaca ccgatcgtca ctcgatcaag aagaacctga
4741 tcggagccct gctgttcgat tcgggagaaa ccgccgaagc cacccgtctg aagcgtaccg
4801 cccgtcgtcg ttacacccgt cggaagaacc gtatctgcta cctgcaagaa atcttctcga
4861 acgaaatggc caaggtggat gattcgttct tccaccgtct ggaagaatcg ttcctggtgg
4921 aagaagataa gaagcacgaa cgtcacccaa tcttcggaaa catcgtggat gaagtggcct
4981 accacgaaaa gtacccaacc atctaccacc tgcggaagaa gctggtggat tcgaccgata
5041 aggccgatct gcgtctgatc tacctggccc tggcccacat gatcaagttc cgtggacact
5101 tcctgatcga aggagatctg aacccagata actcggatgt ggataagctg ttcatccagc
5161 tggtgcagac ctacaaccag ctgttcgaag aaaacccaat caacgcctcg ggagtggatg
5221 ccaaggccat cctgtcggcc cgtctgtcga agtcgcgtcg tctggaaaac ctgatcgccc
5281 agctgccagg agaaaagaag aacggactgt tcggaaacct gatcgccctg tcgctgggac
5341 tgaccccaaa cttcaagtcg aacttcgatc tggccgaaga tgccaagctg cagctgtcga
5401 aggataccta cgatgatgat ctggataacc tgctggccca gatcggagat cagtacgccg
5461 atctgttcct ggccgccaag aacctgtcgg atgccatcct gctgtcggat atcctgcgtg
5521 tgaacaccga aatcaccaag gccccactgt cggcctcgat gatcaagcgt tacgatgaac
5581 accaccagga tctgaccctg ctgaaggccc tcgtgcgtca gcagctgcca gaaaagtaca
5641 aggaaatctt cttcgatcaa tcgaagaacg gatacgccgg atacatcgat ggaggagcct
5701 cgcaggaaga attctacaag ttcatcaagc caatcctgga aaagatggat ggaaccgaag
5761 aactgctggt gaagctgaac cgtgaagatc tgctgcgtaa gcagcgtacc ttcgataacg
5821 gatcgatccc acaccaaatc cacctgggag aactgcacgc catcctgcgt cgtcaggaag
5881 atttctaccc attcctgaaa gataaccgtg aaaagatcga aaagatcctg accttccgta
5941 tcccatacta cgtgggacca ctggcccgtg gaaactcgcg tttcgcctgg atgacccgga
6001 agtcggaaga aaccatcacc ccgtggaact tcgaagaagt ggtggataag ggagcctcgg
6061 cccagtcgtt catcgaacgt atgaccaact tcgataagaa cctgccaaac gaaaaggtgc
6121 tgccaaagca ctcgctgctg tacgaatact tcaccgtgta caacgaactg accaaagtga
6181 agtacgtgac cgaaggaatg cggaagccag ccttcctgtc gggagaacag aagaaggcca
6241 tcgtggatct gctgttcaag accaaccgga aggtgaccgt gaagcagctg aaggaagatt
6301 acttcaagaa gatcgaatgc ttcgattcgg tggaaatctc gggagtggaa gatcgtttca
6361 acgcctcgct gggaacctac cacgatctgc tgaagatcat caaggataag gatttcctgg
6421 ataacgaaga aaacgaagat atcctggaag atatcgtgct gaccctgacc ctgttcgaag
6481 atcgtgaaat gatcgaagaa cgtctgaaaa cctacgccca cctgttcgat gataaagtga
6541 tgaagcagct gaagcgtcgt cgttacaccg gatggggacg tctgtcgcgg aagctgatca
6601 acggaatccg tgataagcag tcgggaaaga ccatcctgga tttcctgaag tcggatggat
6661 tcgccaaccg taacttcatg cagctgatcc acgatgattc gctgaccttc aaggaagata
6721 tccagaaggc ccaggtgtcg ggacagggag attcgctgca cgaacacatc gccaacctgg
6781 ccggatcgcc agccatcaag aagggaatcc tgcagaccgt gaaggtggtg gatgaactgg
6841 tgaaggtgat gggacgtcac aagccagaaa acatcgtgat cgaaatggcc cgtgaaaacc
6901 agaccaccca gaagggacag aagaactcgc gtgaacgtat gaagcgtatc gaagaaggaa
6961 tcaaggaact gggatcgcag atcctgaagg aacacccagt ggaaaacacc cagctgcaga
7021 acgaaaagct gtacctgtac tacctgcaaa acggacgtga tatgtacgtg gatcaggaac
7081 tggatatcaa ccgtctgtcg gattacgatg tggatcacat cgtgccacag tcgttcctga
7141 aggatgattc gatcgataac aaggtgctga cccgttcgga taagaaccgt ggaaagtcgg
7201 ataacgtgcc atcggaagaa gtggtgaaga agatgaagaa ctactggcgt cagctgctga
7261 acgccaagct gatcacccag cggaagttcg ataacctgac caaggccgaa cgtggaggac
7321 tgtcggaact ggataaggcc ggattcatca agcgtcagct ggtggaaacc cgtcagatca
7381 ccaagcacgt ggcccagatc ctggattcgc gtatgaacac caaatacgat gaaaacgata
7441 agctgatccg tgaagtgaag gtgatcaccc tgaagtcgaa gctggtgtcg gatttccgga
7501 aggatttcca gttctacaaa gtccgtgaaa tcaacaacta ccaccacgcc cacgatgcct
7561 acctgaacgc cgtcgtggga accgccctga tcaagaagta cccaaagctg gaatcggaat
7621 tcgtgtacgg agattacaag gtgtacgacg tccgtaagat gatcgccaag tcggaacagg

7681 aaatcggaaa ggccaccgcc aagtacttct tctactcgaa catcatgaac ttcttcaaaa
7741 ccgaaatcac cctggccaac ggagaaatcc ggaagcgtcc actgatcgaa accaacggag
7801 aaaccggaga aatcgtgtgg gataagggac gtgatttcgc caccgtgcgt aaggtgctgt
7861 cgatgccaca agtgaacatc gtgaagaaaa ccgaagtgca gaccggagga ttctcgaagg
7921 aatcgatcct gccaaagcgt aactcggata agctgatcgc ccggaagaag gattgggacc
7981 caaagaagta cggaggattc gattcgccaa ccgtggccta ctcggtgctg gtggtggcca
8041 aggtggaaaa gggaaagtcg aagaagctga agtcggtgaa ggaactgctg ggaatcacca
8101 tcatggaacg ttcgtcgttc gaaaagaacc caatcgattt cctggaagcc aagggataca
8161 aggaagtgaa gaaggatctg atcatcaagc tgccaaagta ctcgctgttc gaactggaaa
8221 acggacggaa gcgtatgctg gcctcggccg gagaactgca gaagggaaac gaactggccc
8281 tgccatcgaa atacgtgaac ttcctgtacc tggcctcgca ctacgaaaag ctgaagggat
8341 cgccagaaga taacgaacag aagcagctgt tcgtggaaca gcacaagcac tacctggatg
8401 aaatcatcga acagatctcg gaattctcga agcgtgtgat cctggccgat gccaacctgg
8461 ataaggtgct gtcggcctac aacaagcacc gtgataagcc aatccgtgaa caggccgaaa
8521 acatcatcca cctgttcacc ctgaccaacc tgggagcccc agccgccttc aagtacttcg
8581 ataccaccat cgatcggaag cgttacacct cgaccaagga agtgctggat gccaccctga
8641 tccaccagtc gatcaccgga ctgtacgaaa cccgtatcga tctgtcgcaa ctgggaggag
8701 ataaaaggcc ggcggccacg aaaaaggccg gccaggcaaa aaagaaaaag gagggcagag
8761 gaagtcttct aacatgcggt gacgtggagg agaatcccgg ccctatggtg agcaagggcg
8821 aggagctgtt caccggggtg gtgcccatcc tggtcgagct ggacggcgac gtaaacggcc
8881 acaagttcag cgtgtccggc gagggcgagg gcgatgccac ctacggcaag ctgaccctga
8941 agttcatctg caccaccagc aagctgcccg tgccctggcc caccctcgtg accaccctga
9001 cctacggcgt gcagtgcttc agccgctacc ccgaccacat gaagcagcac gacttcttca
9061 agtccgccat gcccgaaggc tacgtccagg agcgcaccat cttcttcaag gacgacggca
9121 actacaagac ccgcgccgag gtgaagttcg agggcgacac cctggtgaac cgcatcgagc
9181 tgaagggcat cgacttcaag gaggacggca acatcctggg gcacaagctg gagtacaact
9241 acaacagcca caacgtctat atcatggccg acaagcagaa gaacggcatc aaggtgaact
9301 tcaagatccg ccacaacatc gaggacggca gcgtgcagct cgccgaccac taccagcaga
9361 acacccccat cggcgacggc cccgtgctgc tgcccgacaa ccactacctg agcacccagt
9421 ccgccctgag caaagacccc aacgagaagc gcgatcacat ggtcctgctg gagttcgtga
9481 ccgccgccgg gatcactctc ggcatggacg agctgtacaa gtaattaatt aaatcttgat
9541 acgtctcttc atcaagctaa aaccgtcatc cccgtatagc taatacacaa agaaagagaa
9601 aaaaaataca aaggttgcaa tgatattcaa atttcaacac tgtaacataa cgtaatttta
9661 tcactaaaag tatggtaaaa caaggtaaaa cattttctca ttcgcaacag aatacaaagt
9721 tataaagtta taacacaaat agagacatga gtaataatgt aaaaccaata gcattcagcg
9781 attgttagaa catgaattca tacgtgacgc ttcgtgtcac gtgcataaaa ttctctctct
9841 atgactatct tgtttggatt attgtatctc aaagacaact cgtgtgaaga aacgaactaa
9901 ctgaataaca ttaggtattc gatttgagcc caagtaacaa gatacaaacc gtaataaaga
9961 aaaaaacaaa ctacaaagca cggattaatt gcaattcttg aatgtggtgg caattacgga
10021 caattgcaat cctcggaaat cagaatacaa agaataagct gcaaatgtac tgtttaattt
10081 aaattaaatt aaattgtaaa agtggaataa aactgaaaac ttttaaagtt atcaatgtga
10141 aagaatgaca aattaaacat agctcgaggg cgattaacta gaatgaatcg tttttaaaat
10201 aacaaatcaa ttgttttata atattcgtac gattctttga ttatgtaata aaatgtgatc
10261 attaggaaga ttacgaaaaa tataaaaaat atgagttctg tgtgtataac aaatgctgta
10321 aacgccacaa ttgtgtttgt tgcaaataaa cccatgatta tttgattaaa attgttgttt
10381 tctttgttca tagacaatag tgtgttttgc ctaaacgtgt actgcataaa ctccatgcga
10441 gtgtatagcg agctagtggc taacgcttgc ccccaccaaag tagattcgtc aaaatcctca
10501 atttcatcac cctcctccaa gtttaacatt tggccgtcgg aattaacttc taaagatgcc
10561 acataatcta ataaatgaaa tagagattca aacgtggcgt catcgtccgt ttcgaccatt
10621 tccgaaaaga actcgggcat aaactctatg atttctctgg acgtggtgtt gtcgaaactc
10681 tcaaagtacg cagtcaggaa cgtgcgcgac atgtcgtcgg gaaactcgcg cggaaacatg
10741 ttgttgtaac cgaacgggtc ccatagcgcc aaaaccaaat ctgccagcgt caatagaatg

10801 agcacgatgc cgacaatgga gctggcttgg atagcgattc gagttaacgg ccggggcgcg
10861 cctatcttta catgtagctt gtgcattgaa tccaattata atttgccttg gcaccagctg
10921 agccagacaa gaaagaaagc ttcccagaag tatatcgatt tagaagggtt gacgtcactt
10981 gctgactgca ctaatacagc aaatgatgca attagaatga ttcaagtgaa attcccaaat
11041 tactgatttt tctctggatt tggttatcag attacattcg aagctaagat tagctaccga
11101 aattgtcgat caaatcagga aatcctttct ctatcgaaaa aggcattcgc acatcttcct
11161 ctctatgcca tatacacgaa gggtaggtac attgacgtct ttgccagaag ttgaactgca
11221 tcgttcaagg tacagaatga acgactaaca gacacaagca cgttttgctg tccattcaga
11281 cacagggatg gtacccatat tcgatcgata tagagccatc caaccgaaca gaggtatatg
11341 tatgaatgta ttgctgaaat tttctagaag tacaaccacc actacgacag tgtctataaa
11401 acgcccctgc aaaggcgaaa ccagctcaat cgaatacgtt tcctagtgga gtgaacatta
11461 cgcggcccaa gtaagcagtg ccagtgcaag tgaagtgaag tctctagtga aaaagagtga
11521 tccaattagc cagaggagaa aatttcagag tgaacaaagc tttattcaaa ggacaattac
11581 tattaaattg gtgaaagtgc atttcggtga agggaatctt ctagtgaagg taggtaaatt
11641 aattgatgaa attatagcta tgagcgaaaa ctagtttggt gaatgattcc tttgtctttg
11701 aatgagcaaa ctattttcca agatggcgac tattgagctt tgagtgatta gtgaaaattt
11761 gcaacgcagt ttcatcatca ttgataaaac ccaattgtga ttcacagcga taatcatatt
11821 tcgttgaatc atcgctacta attgaattaa atttctagaa taataagaat aacgtatttg
11881 ctccgtcaca tatctaaaat aaatattttg atggtaatta cccattaagg taatattaac
11941 acatatcgag aaaaaccttg aggaaatcgt gaaaacttga agatacgcaa tttcaaaact
12001 acgtagttca aagtcgaaaa caagttaatt tttcacttaa aagtagggcg ttgttgtgac
12061 gtcatcacct tcaagtgtat atttttcact tggcctgcga ctgcaaacgc agacaaagca
12121 aaacaagttt aaaacctgtc gtgtcgtgct cgaagccaaa ggcaatgaat caatatcaaa
12181 tgagagtttg catttcacaa ccaattactc aagcgtttcc tcgtttcttt ttctgctcaa
12241 cagagatttc aacatggtgt cgaagggaga agaagataac atggccatca tcaaggagtt
12301 catgcgtttc aaagtccaca tggaaggatc ggtgaacgga cacgaatttg aaatcgaagg
12361 agaaggagaa ggacgcccgt acgaaggaac ccagaccgcc aagctgaaag tgacgaaggg
12421 aggaccgctg ccgttcgcct gggatatcct gtcgccgcag ttcatgtacg gatcgaaggc
12481 ctacgtgaag cacccggccg atatcccgga ttacctgaag ctgtcgttcc cggaaggatt
12541 caagtgggaa cgtgtgatga actttgaaga tggaggagtg gtgactgtga cccaggattc
12601 gtcgttgcag gatggagagt tcatctacaa ggtgaagctg cgtggaacca acttcccgtc
12661 ggatggacca gtgatgcaga agaagactat gggctgggaa gcctcgtcgg aacgcatgta
12721 cccggaagat ggagccctga agggagaaat caagcagcgt ctgaagctga aggatggagg
12781 acactacgat gccgaagtga agaccaccta caaggccaag aagccggtgc agctgccggg
12841 agcctacaac gtgaacatca agctggatat cacctcgcac aacgaagatt acaccatcgt
12901 ggaacagtac gaacgtgccg aaggacgcca ctcgaccgga ggaatggatg aactgtacaa
12961 gtaatgatca taatcagcca taccacattt gtagaggttt tacttgcttt aaaaaacctc
13021 ccacacctcc ccctgaacct gaaacataaa atgaatgcaa ttgttgttgt taacttgttt
13081 attgcagctt ataatggtta caaataaagc aatagcatca caaatttcac aaataaagca
13141 tttttttcac tgcattctag ttgtggtttg tccaaactca tcaatgtatc ttagggcccA
13201 GGTgccggcc gatctcggat ctgacaatgt tcagtgcaga gactcggcta cgcctcgtgg
13261 actttgaagt tgaccaacaa tgtttattct tacctctaat agtcctctgt ggcaaggtca
13321 agattctgtt agaagccaat gaagaacctg gttgttcaat aacattttgt tcgtctaata
13381 tttcactacc gcttgacgtt ggctgcactt catgtacctc atctataaac gcttcttctg
13441 tatcgctctg gacgtcatct tcacttacgt gatctgatat ttcactgtca gaatcctcac
13501 caacaagctc gtcatcgctt tgcagaagag cagagaggat atgctcatcg tctaaagaac
13561 tacccatttt attatatatt agtcacgata tctataacaa gaaaatatat atataataag
13621 ttatcacgta agtagaacat gaaataacaa tataattatc gtatgagtta aatcttaaaa
13681 gtcacgtaaa agataatcat gcgtcatttt gactcacgcg gtcgttatag ttcaaaatca
13741 gtgacactta ccgcattgac aagcacgcct cacgggagct ccaagcggcg actgagatgt
13801 cctaaatgca cagcgacgga ttcgcgctat ttagaaagag agagcaatat ttcaagaatg
13861 catgcgtcaa ttttacgcag actatctttc tagggttaaa aaagatttgc gctttactcg

13921 acctaaactt taaacacgtc atagaatctt cgtttgacaa aaaccacatt gtggccaagc
13981 tgtgtgacgc gacgcgcgct aaagaatggc aaaccaagtc gcgcgagcgt cgactctaga
14041 ggatccccgg gtaccgagct cgaattcgta atcatggtca tagctgtttc ctgtgtgaaa
14101 ttgttatccg ctcacaattc cacacaacat acgagccgga agcataaagt gtaaagcctg
14161 gggtgcctaa tgagtgagct aactcacatt aattgcgttg cgctcactgc ccgctttcca
14221 gtcgggaaac ctgtcgtgcc agctgcatta atgaatcggc caacgcgcgg ggagaggcgg
14281 tttgcgtatt gggcgctctt ccgcttcctc gctcactgac tcgctgcgct cggtcgttcg
14341 gctgcggcga gcggtatcag ctcactcaaa ggcggtaata cggttatcca cagaatcagg
14401 ggataacgca ggaaagaaca tgtgagcaaa aggccagcaa aaggccagga accgtaaaaa
14461 ggccgcgttg ctggcgtttt tccataggct ccgcccccct gacgagcatc acaaaaatcg
14521 acgctcaagt cagaggtggc gaaacccgac aggactataa agataccagg cgtttccccc
14581 tggaagctcc ctcgtgcgct ctcctgttcc gaccctgccg cttaccggat acctgtccgc
14641 ctttctccct tcgggaagcg tggcgctttc tcaatgctca cgctgtaggt atctcagttc
14701 ggtgtaggtc gttcgctcca agctgggctg tgtgcacgaa ccccccgttc agcccgaccg
14761 ctgcgcctta tccggtaact atcgtcttga gtccaacccg gtaagacacg acttatcgcc
14821 actggcagca gccactggta acaggattag cagagcgagg tatgtaggcg gtgctacaga
14881 gttcttgaag tggtggccta actacggcta cactagaagg acagtatttg gtatctgcgc
14941 tctgctgaag ccagttacct tcggaaaaag agttggtagc tcttgatccg gcaaacaaac
15001 caccgctggt agcggtggtt tttttgtttg caagcagcag attacgcgca gaaaaaaagg
15061 atctcaagaa gatcctttga tcttttctac ggggtctgac gctcagtgga acgaaaactc
15121 acgttaaggg attttggtca tgagattatc aaaaaggatc ttcacctaga tccttttaaa
15181 ttaaaaatga agttttaaat caatctaaag tatatatgag taaacttggt ctgacagtta
15241 ccaatgctta atcagtgagg cacctatctc agcgatctgt ctatttcgtt catccatagt
15301 tgcctgactc cccgtcgtgt agataactac gatacgggag ggcttaccat ctggccccag
15361 tgctgcaatg ataccgcgag acccacgctc accggctcca gatttatcag caataaacca
15421 gccagccgga agggccgagc gcagaagtgg tcctgcaact ttatccgcct ccatccagtc
15481 tattaattgt tgccgggaag ctagagtaag tagttcgcca gttaatagtt tgcgcaacgt
15541 tgttgccatt gctacaggca tcgtggtgtc acgctcgtcg tttggtatgg cttcattcag
15601 ctccggttcc caacgatcaa ggcgagttac atgatccccc atgttgtgca aaaaagcggt
15661 tagctccttc ggtcctccga tcgttgtcag aagtaagttg gccgcagtgt tatcactcat
15721 ggttatggca gcactgcata attctcttac tgtcatgcca tccgtaagat gcttttctgt
15781 gactggtgag tactcaacca agtcattctg agaatagtgt atgcggcgac cgagttgctc
15841 ttgcccggcg tcaatacggg ataataccgc gccacatagc agaactttaa aagtgctcat
15901 cattggaaaa cgttcttcgg ggcgaaaact ctcaaggatc ttaccgctgt tgagatccag
15961 ttcgatgtaa cccactcgtg cacccaactg atcttcagca tcttttactt tcaccagcgt
16021 ttctgggtga gcaaaaacag gaaggcaaaa tgccgcaaaa aagggaataa gggcgacacg
16081 gaaatgttga atactcatac tcttcctttt tcaatattat tgaagcattt atcagggtta
16141 ttgtctcatg agcggataca tatttgaatg tatttagaaa aataaacaaa taggggttcc
16201 gcgcacattt ccccgaaaag tgccacctga cgtctaagaa accattatta tcatgacatt
16261 aacctataaa aataggcgta tcacgaggcc ctttcgtctc gcgcgtttcg gtgatgacgg
16321 tgaaaacctc tgacacatgc agctcccgga gacggtcaca gcttgtctgt aagcggatgc
16381 cgggagcaga caagcccgtc agggcgcgtc agcgggtgtt ggcgggtgtc ggggctggct
16441 taactatgcg gcatcagagc agattgtact gagagtgc
//

>AGG1245

FEATURES Location/Qualifiers
gene complement(join(6115..6554,1..421))
/label="AmpR gene"
Promoter 1619..3000

/label="Aedes aegypti polyubiquitin Promoter"
source join(4793..6554,1..1299)
/label="luciferase reporter vector"
CDS 3006..4790
/label="AaHyPiggyBac"
misc_binding 1326..1610
/label="attB site"
regulatory 1139..1292
/label="upstream poly(A) signal"
rep_origin 553..1008
/label="f1 origin"
regulatory 4807..5028
/label="SV40 late poly(A) signal"
rep_origin 5353..5353
/label="ColE1 origin"
misc_feature 1300..1306
/label="multiple cloning site"

ORIGIN

```
        1 cgatcgttgt cagaagtaag ttggccgcag tgttatcact catggttatg gcagcactgc
       61 ataattctct tactgtcatg ccatccgtaa gatgcttttc tgtgactggt gagtactcaa
      121 ccaagtcatt ctgagaatag tgtatgcggc gaccgagttg ctcttgcccg gcgtcaatac
      181 gggataatac cgcgccacat agcagaactt taaaagtgct catcattgga aaacgttctt
      241 cggggcgaaa actctcaagg atcttaccgc tgttgagatc cagttcgatg taacccactc
      301 gtgcacccaa ctgatcttca gcatctttta ctttcaccag cgtttctggg tgagcaaaaa
      361 caggaaggca aaatgccgca aaaaagggaa taagggcgac acggaaatgt tgaatactca
      421 tactcttcct ttttcaatat tattgaagca tttatcaggg ttattgtctc atgagcggat
      481 acatatttga atgtatttag aaaaataaac aaataggggt tccgcgcaca tttccccgaa
      541 aagtgccacc tgacgcgccc tgtagcggcg cattaagcgc ggcgggtgtg gtggttacgc
      601 gcagcgtgac cgctacactt gccagcgccc tagcgcccgc tcctttcgct ttcttccctt
      661 cctttctcgc cacgttcgcc ggctttcccc gtcaagctct aaatcggggg ctccctttag
      721 ggttccgatt tagtgcttta cggcacctcg accccaaaaa acttgattag ggtgatggtt
      781 cacgtagtgg gccatcgccc tgatagacgg tttttcgccc tttgacgttg gagtccacgt
      841 tctttaatag tggactcttg ttccaaactg gaacaacact caaccctatc tcggtctatt
      901 cttttgattt ataagggatt ttgccgattt cggcctattg gttaaaaaat gagctgattt
      961 aacaaaaatt taacgcgaat tttaacaaaa tattaacgct tacaatttgc cattcgccat
     1021 tcaggctgcg caactgttgg gaagggcgat cggtgcgggc ctcttcgcta ttacgccagc
     1081 ccaagctacc atgataagta agtaatatta aggtacggga ggtacttgga gcggccgcaa
     1141 taaaatatct ttattttcat tacatctgtg tgttggtttt ttgtgtgaat cgatagtact
     1201 aacatacgct ctccatcaaa acaaaacgaa acaaaacaaa ctagcaaaat aggctgtccc
     1261 cagtgcaagt gcaggtgcca gaacatttct ctatcgatag gtaccgagct cgcccggggt
     1321 cctaggtcga cgatgtaggt cacggtctcg aagccgcggt gcgggtgcca gggcgtgccc
     1381 ttgggctccc cgggcgcgta ctccacctca cccatctggt ccatcatgat gaacgggtcg
     1441 aggtggcggt agttgatccc ggcgaacgcg cggcgcaccg ggaagccctc gccctcgaaa
     1501 ccgctgggcg cggtggtcac ggtgagcacg ggacgtgcga cggcgtcggc gggtgcggat
     1561 acgcggggca gcgtcagcgg gttctcgacg gtcacggcgg gcatgtcgac gcggccgcta
     1621 tctttacatg tagcttgtgc attgaatcca attataattt gccttggcac cagctgagcc
     1681 agacaagaaa gaaagcttcc cagaagtata tcgatttaga agggttgacg tcactttgct
     1741 gactgcacta atacagcaaa tgatacaatt agaatgattc aagtgaaatt cccaaattac
     1801 tgctttgtct ctggatttgg ttatcagatt acattcgaag ctaagaatag ctaccgaaat
     1861 tgtcgatcaa atcaggaaat cctttctcta tcgaaaaagg cattcgcaca tctttctctg
     1921 tatgccatat acacgaatgg taggtacatt gacgtctttg ccagaagttg aactgcatcg
     1981 ttcaaggtac agaatgaacg actaacagac acaagcacgt tttgctgtcc attcagacac
```

2041 agggatggta cccatagtcg atcgatttag agccatccaa ccgaacagag gtatatgtat
2101 gaatggattg cagaaatttt ctagaagtac aaccaccact acggcagtgt ctataaaacg
2161 cccctgcaaa ggcaaaacca gctcaatcga atacgtttcc tagtggagtg aacattacgc
2221 ggtccaagta agcagtgcca gtgcaagtga agtgaagtct ctagtgaaaa agagtgatcc
2281 aattagccag aggagaaaat ttcagagtga acaaagcttt gttcaaagga caattactat
2341 taaatttgtg aaagtgcatt tcggtgaagg gaatcttcta gtgaaggtag gtaaattaaa
2401 tgatgaaatt atagctatga gcgaaaacta gtttggtgaa tgattccttt gtctttgaat
2461 gagcaaacta ttttccaaga tggcgactat tgagctttga gtgattagtg aaaatttgca
2521 acgcagtttc atcatcattg ataaaaccca attgtgattc acggcgataa tcatatttcg
2581 ttgaatcatc gctgctaatt gaattaaatt tctagagcaa gcgcgaattc gccatatttc
2641 taaaattaaa tattgtggtg ataattaccc attaaggtaa tattaacaca tatcgagaaa
2701 aaccttgagg aaatcgtgaa aacttgaaga tacgcaattt ccaaactacg tagttcaaag
2761 tcgaaaacaa gttaattttt cactaaaaag tagggcgttg ttgtgacgtc atcaccttca
2821 agtgtatatt tttcacttgg cctgcgactg caaacgcaga caaagcaaaa caagtttaaa
2881 acctgtcgtg tcgtgctcga agccaaaggc aatgaatcaa tatcaaatga gagtttgcat
2941 ttcacaacca attactgaag cgtttcctcg tttctttttc tgctcaacag agatttcaac
3001 ccaccatggg atcgtcgctg gatgatgaac acatcctgtc ggccctgctc cagtcggatg
3061 atgaactcgt tggagaagat tcggattcgg aagtgtcgga tcacgtgtcg gaagatgatg
3121 tgcagtcgga taccgaagaa gccttcatcg atgaagtgca cgaagtgcag ccgacctcgt
3181 cgggatcgga aatcctggat gaacagaacg tgatcgaaca gccgggatcg tcgctggcct
3241 cgaaccgtat cctgaccctg ccgcagcgta ccatccgtgg aaagaacaag cactgctggt
3301 cgacctcgaa gccgacccgt cgttcgcgtg tgtcggccct gaacatcgtg cgttcccaac
3361 gtggaccgac gcgcatgtgc cgtaacatct acgatccgct gctgtgcttc aagctgttct
3421 tcaccgatga aatcatctcg gaaatcgtga agtggaccaa cgccgaaatc tcgctgaagc
3481 gtcgtgaatc gatgacctcg gccaccttcc gtgataccaa cgaagatgaa atctacgcct
3541 tcttcggaat cctggtgatg accgccgtgc gtaaggataa ccacatgtcg accgatgatc
3601 tgttcgatcg ttcgctgtcg atggtctacg tgtcggtgat gtcgcgtgat cgtttcgatt
3661 tcctgatccg ttgcctgaag atggatgata agtcgatccg tccgaccctg cgtgaaaacg
3721 atgtgttcac ccccgtccgc aagatctggg atctgttcat ccaccagtgc atccagaact
3781 acaccccccgg agcccacctg accatcgatg aacagctgct gggattccgt ggacgttgcc
3841 cgttccgtgt gtacatcccg aacaagccgt cgaagtacgg aatcaagatc ctgatgatgt
3901 gcgattcggg aaccaagtac atgatcaacg gaatgccgta cctgggacgt ggaacccaga
3961 ccaacggagt gccgctggga gaatactacg tgaaggaact gtcgaagccg gtgcacggat
4021 cgtgccgtaa catcacctgc gataactggt tcacctcgat cccgctggcc aagaacctgc
4081 tccaggaacc gtacaagctg accatcgtgg gaaccgtgcg ttcgaacaag cgtgaaatcc
4141 cggaagtgct gaagaactcg cgttcgcgtc cggtgggaac ctcgatgttc tgcttcgatg
4201 gaccgctgac cctggtgtcg tacaagccga agccggccaa gatggtgtac ctgctgtcgt
4261 cgtgcgatga agatgcctcg atcaacgaat cgaccggaaa gccgcagatg gtgatgtact
4321 acaaccagac caagggagga gtggataccc tggatcagat gtgctcggtg atgacctgct
4381 cccgcaagac caaccgttgg ccgatggccc tgctgtacgg aatgatcaac atcgcctgca
4441 tcaactcgtt catcatctac tcgcacaacg tgtcgtcgaa gggagaaaag gtgcagtcgc
4501 gcaaaaagtt catgcgtaac ctgtacatgg gactgacctc gtcgttcatg cgcaaacgtc
4561 tggaagcccc gaccctgaag cgttacctgc gtgataacat ctcgaacatc ctgccgaagg
4621 aagtgccggg aacctcggat gattcgaccg aagaaccggt gatgaagaag cgtacctact
4681 gcacctactg cccgtcgaag atccgtcgca agtcgaacgc ctcgtgcaag aagtgcaaga
4741 aggtgatctg ccgtgaacac aacatcgata tgtgccagtc gtgcttctaa ggccggccgc
4801 ttcgagcaga catgataaga tacattgatg agtttggaca aaccacaact agaatgcagt
4861 gaaaaaaatg ctttatttgt gaaatttgtg atgctattgc tttatttgta accattataa
4921 gctgcaataa acaagttaac aacaacaatt gcattcattt tatgtttcag gttcaggggg
4981 aggtgtggga ggttttttaa agcaagtaaa acctctacaa atgtggtaaa atcgataagg
5041 atccgtcgac cgatgccctt gagagccttc aacccagtca gctccttccg gtgggcgcgg
5101 ggcatgacta tcgtcgccgc acttatgact gtcttcttta tcatgcaact cgtaggacag

5161 gtgccggcag cgctcttccg cttcctcgct cactgactcg ctgcgctcgg tcgttcggct
5221 gcggcgagcg gtatcagctc actcaaaggc ggtaatacgg ttatccacag aatcagggga
5281 taacgcagga aagaacatgt gagcaaaagg ccagcaaaag gccaggaacc gtaaaaaggc
5341 cgcgttgctg gcgtttttcc ataggctccg cccccctgac gagcatcaca aaaatcgacg
5401 ctcaagtcag aggtggcgaa acccgacagg actataaaga taccaggcgt ttccccctgg
5461 aagctccctc gtgcgctctc ctgttccgac cctgccgctt accggatacc tgtccgcctt
5521 tctcccttcg ggaagcgtgg cgctttctca tagctcacgc tgtaggtatc tcagttcggt
5581 gtaggtcgtt cgctccaagc tgggctgtgt gcacgaaccc cccgttcagc ccgaccgctg
5641 cgccttatcc ggtaactatc gtcttgagtc caacccggta agacacgact tatcgccact
5701 ggcagcagcc actggtaaca ggattagcag agcgaggtat gtaggcggtg ctacagagtt
5761 cttgaagtgg tggcctaact acggctacac tagaagaaca gtatttggta tctgcgctct
5821 gctgaagcca gttaccttcg gaaaaagagt tggtagctct tgatccggca aacaaaccac
5881 cgctggtagc ggtggttttt ttgtttgcaa gcagcagatt acgcgcagaa aaaaaggatc
5941 tcaagaagat cctttgatct tttctacggg gtctgacgct cagtggaacg aaaactcacg
6001 ttaagggatt ttggtcatga gattatcaaa aaggatcttc acctagatcc ttttaaatta
6061 aaaatgaagt tttaaatcaa tctaaagtat atatgagtaa acttggtctg acagttacca
6121 atgcttaatc agtgaggcac ctatctcagc gatctgtcta tttcgttcat ccatagttgc
6181 ctgactcccc gtcgtgtaga taactacgat acgggagggc ttaccatctg gccccagtgc
6241 tgcaatgata ccgcgagacc cacgctcacc ggctccagat ttatcagcaa taaaccagcc
6301 agccggaagg gccgagcgca gaagtggtcc tgcaacttta tccgcctcca tccagtctat
6361 taattgttgc cgggaagcta gagtaagtag ttcgccagtt aatagtttgc gcaacgttgt
6421 tgccattgct acaggcatcg tggtgtcacg ctcgtcgttt ggtatggctt cattcagctc
6481 cggttcccaa cgatcaaggc gagttacatg atcccccatg ttgtgcaaaa aagcggttag
6541 ctccttcggt cctc
//

**S2. Flanking sequences for *bgcn*-Cas9 lines.** Flanking PCRs were performed on *bgcn*-Cas9 transgenics by digesting the gDNA extracted from 10 individuals with the restriction enzymes BamHI, MspI and NcoI (New England Biolabs). Sequencing of the fragments flanking the piggyBac sites was performed from purified the PCRs products. Primers used: LA182, LA184, LA186 and LA187 (see Table S11). BLAST search was performed to find potential insertion site in the genome.

***bgcn*-Cas9 line D**

BamHI
GAGACGACAGAAAGGGCGTGGTGCGGAAGGCGGTGGATCCCGGAAATCAAATAAATGTGCATAC
TGATATTCTGGTATATGGTGAGGATTCTGGCATTCAGTGGGAATGGTTAAATGGAAATCCTAGGAA
TCCTGTGAGAATCTTGGGCTTTCAGTGGCAATACGTGGGAATTCTGTGTGTTATATTGAAATGCGT
GCAAATGTTGTGGAGTAGGGTGATGTGCTAGTATCCATCGTATTAAGCATTGATCAGTGAAAACCT
GCAATTTTCCCATTACTGCAGTGAATGATGCTGATACAAACCGATGCCAAAAAATTCTTTCCCAAA
ACAGCTTTCGCATTGTACAATAATACTTTGATAAATCTTGATCGTCTTTCTGGAAATAATGCAATGAA
ATTGCATCTTGCCGTATTCTTAATTATTCGTATCGTTCTCATTTCTGTCGCAATCAGTATTCAGATTC
GTCTCACAAATTTCGGCTCATTGTGATAAGTTTTAGGTCAAAATACCACACTTCTTCTTTCTCTTCT
TTCTGGCGTTACGTCCCCACTGGGATAGAGCTTGCTTCTCAACTCAAGTGTTCTTATGAGCACTT
CCATAGTTATTAACTGAGAGCTTACTATGCCAGCAATGACCATTTATATCGAGTGGCAGGTACGATG
ATACTTTATGCCCTGGGAAGTCGAGAAAATTTCCAATCCGAAAATATCCTCGGCCGGTGGGATTC
GAACCCACGACCCTCAGCTTTTTCTTGCTGAATAGCTGCGCGTTTACCGCTACGGCTATCTTGGC
CCCAAAAATACCACACATCAACGCTATTATAAAATATTTTACTTGAATACATAGATCTACAAGCATCT
CTCTCCATTCCTTCAGCATAATGAAATGTATAAATTTCCTTTAACCCTAGAAAGATAGTCTGCGTAAA
ATTGACGCATGCATTCTTAAAATATTGCTCTCTCTTTCTAAATAGCGCGAATTCCGTCGCTGGA

Length=1032

Score E

AaegL5_1 | organism=Aedes_aegypti_LVP_AGWG | version=AaegL5 | l... 785 0.0
AaegL5_2 | organism=Aedes_aegypti_LVP_AGWG | version=AaegL5 | l... 751 0.0
AaegL5_3 | organism=Aedes_aegypti_LVP_AGWG | version=AaegL5 | l... 708 0.0

Sequence found repetitive along *Ae. aegypti* genome

MspI
AGCGTGAAGACGACAGAAAGGGCGTGGTGCGGAGGGCGGGGTGTAGCGTGAAGACGACAGAA
AGGGCGTGGTGCGGAGGGCGGTGCGGTGGGATTCGAACCCACGACCCTCAGCTTTTTCTTGCT
GAATAGCTGCGCGTTTACCGCTACGGCTATCTTGGCCCCAAAAATACCACACATCAACGCTATTAT
AAAATATTTTACTTGAATACATAGATCTACAAGCATCTCTCTCCATTCCTTCAGCATAATGAAATGTAT
AAATTTCCTTTAACCCTAGAAAGATAGTCTGCGTAAAATTGACGCATGCATCCTTGAAATATTGCTC
TCTCTTTCTAAATAGCGCGAATCCGTCGCAA

Length=358

Score E
AaegL5_1 | organism=Aedes_aegypti_LVP_AGWG | version=AaegL5 | l... 239 6e-60
AaegL5_2 | organism=Aedes_aegypti_LVP_AGWG | version=AaegL5 | l... 214 2e-52
AaegL5_3 | organism=Aedes_aegypti_LVP_AGWG | version=AaegL5 | l... 210 3e-51

Sequence found repetitive along *Ae. aegypti* genome

NcoI

GGTTACTTCTACACCAAAAATGCAATAAAACTACTGTAAATTCAATAAATAACTCTAGTTAAATTATCC
AATTGCACTTTTCATCGAAATCGCATGGACTAACACGATACGAAAAAACTGCTTAGCGATTGACAA
CAGTCACACCAGAGTGGATCTAAATAAGAGTGGCGTCAATGTCAAAATTGGAACAGCGGCGAACT
GTGAATTCCATACAAATCCGCGTTCCAATGAAGCAGGAGAAAAGGGAAAGGGCGGAGGGGGGTT
CACTCTTGTTTACAGGCACTCTAAGTCACACCACTTTGTGATACAAGTCTCTTCCCGACTACCCCC
ATACTCTACCCCTCTCGGCACGCCATTTTCACTAAAACCTACAAACATAATCACATAACGACTTAAT
TCAAATAAAACCACGCAAAAAAATTACACATCACAAAAATTTATAAAATTTCACCAATGACCTCTTTT
AATTTAAAAAGCAAACAAAGTTAAATCCATTGTAATGAAAAAAAAAACAAACTTTTGCCCATCAACG
TGTGCGGGAAGCCCCGCGTATAAAAAAAGGTTCCTTTGATTGTGTTGTTTTTCCTGGCACTGGAA
CAAATATCCAGATCAATCCCTCAAAAAAAAAAGCCATAATATTTTGAGACCCAATAAGAAGGGCAAA
AATTAAATTTAACCCTAAAAAGAAAATCAATATTTGAAGTACCTTAAAAAAAAACAAGCCTTAAATTTA
CCC

Length=736

| | Score | E |
|---|---|---|
| AaegL5_3 \| organism=Aedes_aegypti_LVP_AGWG \| version=AaegL5 \| l... | 176 | 1e-40 |
| AaegL5_2 \| organism=Aedes_aegypti_LVP_AGWG \| version=AaegL5 \| l... | 174 | 4e-40 |
| AaegL5_1 \| organism=Aedes_aegypti_LVP_AGWG \| version=AaegL5 \| l... | 174 | 4e-40 |

Sequence found repetitive along *Ae. aegypti* genome

**S3. Scripts used to perform the deep sequencing analysis of the cutting assay samples and cage trial samples.**

a) Script from the cutting assessment

```
#!/bin/sh
#Prerequisite - FastQC, Trimmomatic, fastx_trimmer, Seqkit, BWA, samtools, pysamstats, python2.7
& python packages (pandas, numpy, matplotlib & seaborn)

#Select Raw fastq file for the FastQC
pathsa=$(zenity --file-selection --title="Select a Raw FastQ file for FastQC"); echo "\"$pathsa\"
selected.";

#FastQc analysis of the raw .fq file
./fastqc $pathsa

#Trimming of adapters using the Trimmomatic Tool
#Please change the directories of the Trimmomatic & Nextera adapters
java -jar /data/purusothaman/indels/Trimmomatic-0.36/trimmomatic-0.36.jar SE -phred33 $pathsa
${pathsa%%.fastq.gz}"trimmed.fq"
ILLUMINACLIP:/data/purusothaman/Trimmomatic-0.36/adapters/NexteraPE-PE.fa:2:30:10
LEADING:3 TRAILING:3

#FastQc of the trimmed fastq file
./fastqc ${pathsa%%.fastq.gz}"trimmed.fq"

#Trimming the library reads to 200bp window
fastx_trimmer -f 150 -l 350 -v -i ${pathsa%%.fastq.gz}"trimmed.fq" -o
${pathsa%%.fastq.gz}"trimmed150_350.fq"
seqkit seq -m 200 ${pathsa%%.fastq.gz}"trimmed150_350.fq" -o
${pathsa%%.fastq.gz}"trimmed150_350_200bp.fq"

#Select reference file for creating GMAP database
pathsb=$(zenity --file-selection --title="Select a reference file for creating BWA database"); echo
"\"$pathsb\" selected.";

#Building BWA database
bwa index $pathsb

#Alignment of trimmed fasta file using BWA & generation of sam file
bwa mem $pathsb ${pathsa%%.fastq.gz}"trimmed150_350_200bp.fq" >
${pathsa%%.fastq.gz}"trimmed150_350_200bp.sam"

# Conversion of sam to bam file
samtools view -bS ${pathsa%%.fastq.gz}"trimmed150_350_200bp.sam" >
${pathsa%%.fastq.gz}"trimmed150_350_200bp.bam"

#Sorting bam files
samtools sort ${pathsa%%.fastq.gz}"trimmed150_350_200bp.bam" >
${pathsa%%.fastq.gz}"trimmed150_350_200bp_sorted.bam"
```

```
#index bam files
samtools index ${pathsa%%.fastq.gz}"trimmed150_350_200bp_sorted.bam"

#Generation of variant file using pysamstats
pysamstats --type variation -D 5000000 ${pathsa%%.fastq.gz}"trimmed150_350_200bp_sorted.bam"
--fasta $pathsb > input.variant.stats

#python script for variant analysis and plotting
python indel.py
```

b) Script from cutting indel calculation

```
import sys,csv,re,glob
import numpy as np
import pandas as pd
import matplotlib as mp
import seaborn

#Read variant file using pandas
variants = pd.read_csv('input.variant.stats','\t')

#Calculate inDel Rate
variants['indel_rate'] = (variants['insertions'] + variants['deletions']) / variants['reads_all']

#Enter the sgRNA Position in the reference sequence
numpos = input("Enter a sgRNA position: ")
kmo_ref_sgRNA_pos = numpos ;

#variant analysis
sgRNA_pos = pd.DataFrame(data = {'chrom' : ['kmo_ref'], 'sgRNA_pos' : [kmo_ref_sgRNA_pos]})
variants = pd.merge(variants,sgRNA_pos)
variants['rel_pos'] = variants['pos'] - variants['sgRNA_pos']

#Select window size for plotting, +/- 100 selects the 100bp upstream and downstream of the sgRNA
cutsite
variants_trimmed = variants[(variants['rel_pos'] > -100) & (variants['rel_pos'] < 100)]
new = variants_trimmed[['chrom', 'indel_rate', 'rel_pos']].copy()

#write the plotting data to CSV, you can use this file for making plots using GraphPad Prism
new.to_csv('variant.csv', sep=',')

#Generating the mutational profile using seaborn point plot
chrom_plot =
seaborn.pointplot(data=variants_trimmed,x='rel_pos',y='indel_rate',hue='chrom',height=10,aspect=5)

#Save the generated plot to pwd
chrom_plot.figure.savefig('/user/directory/image1.png')

#Calculation of the maximum cut rate
maximum_cut_rate = variants_trimmed.groupby('chrom').apply(max)['indel_rate']
print(maximum_cut_rate)
```

c) Script from cage trial assay

```
#!/bin/sh
#Prerequisite - FastQC, Trimmomatic, fastx_trimmer, Seqkit, BWA, samtools, awk.

#Select Raw fastq file for the FastQC
pathsa=$(zenity --file-selection --title="Select a Raw FastQ file for FastQC"); echo "\"$pathsa\"
selected.";

#FastQc analysis of the raw .fq file
./fastqc $pathsa

#Trimming of adapters using the Trimmomatic Tool
#Please change the directories of the Trimmomatic & Nextera adapters
java -jar /data/purusothaman/indels/Trimmomatic-0.36/trimmomatic-0.36.jar SE -phred33 $pathsa
${pathsa%%.fastq.gz}"trimmed.fq"
ILLUMINACLIP:/data/purusothaman/Trimmomatic-0.36/adapters/NexteraPE-PE.fa:2:30:10
LEADING:3 TRAILING:3

#FastQc of the trimmed fastq file
./fastqc ${pathsa%%.fastq.gz}"trimmed.fq"

#Trimming the library reads to 200bp window
fastx_trimmer -f 150 -l 350 -v -i ${pathsa%%.fastq.gz}"trimmed.fq" -o
${pathsa%%.fastq.gz}"trimmed150_350.fq"
seqkit seq -m 200 ${pathsa%%.fastq.gz}"trimmed150_350.fq" -o
${pathsa%%.fastq.gz}"trimmed150_350_200bp.fq"

#Select reference file for creating GMAP database
pathsb=$(zenity --file-selection --title="Select a reference file for creating BWA database"); echo
"\"$pathsb\" selected.";

#Building BWA database
bwa index $pathsb

#Alignment of trimmed fasta file using BWA & generation of sam file
bwa mem $pathsb ${pathsa%%.fastq.gz}"trimmed150_350_200bp.fq" >
${pathsa%%.fastq.gz}"trimmed150_350_200bp.sam"

#Calculation of number of reads where all 4 sgRNA sites were not available
#Add sgRNA target sequences including PAM site
samtools view -h -F 4 ${pathsa%%.fastq.gz}"trimmed150_350_200bp.sam" | awk '$10 !~
/sgRNA1_targetseq+PAM/ && $10 !~ /sgRNA2_targetseq+PAM/ && $10 !~
/sgRNA3_targetseq+PAM/ && $10 !~ /sgRNA4_targetseq+PAM/' | samtools view -bS - >
${pathsa%%.fastq.gz}"trimmed150_350_200bp_no4sgRNAs.bam"
samtools view -c ${pathsa%%.fastq.gz}"trimmed150_350_200bp_no4sgRNAs.bam"

#-----Please Note----------
#Calculation of number of reads where atleast 1 sgRNA sites are available for cutting
#Total Number of mapped reads in the library - No of reads where all 4 sgRNA sites are not available.
```

### S4 Mathematical Modelling

#### S4.1 Fitting Relative Fitness for Individual Gene Drive Constructs

To obtain relative fitness parameter values for individuals heterozygous or homozygous for a single transgenic construct, we first formulate a deterministic population genetics mathematical model that considers an infinite, closed (no migration), panmictic (random mating) population. This model is simulated using the following set of difference equations

$$m_{ww}(t) = \frac{1}{2}\Big[m_{ww}(t-1)f_{ww}(t-1) + \frac{1}{2}m_{ww}(t-1)f_{Xw}(t-1) + \frac{1}{2}m_{Xw}(t-1)f_{ww}(t-1) + \frac{1}{4}m_{Xw}(t-1)f_{Xw}(t-1)\Big]/\tilde{\Omega}, \tag{1}$$

$$m_{Xw}(t) = \frac{\Omega_{Xw}}{2}\Big[\frac{1}{2}m_{ww}(t-1)f_{Xw}(t-1) + m_{ww}(t-1)f_{XX}(t-1) + \frac{1}{2}m_{Xw}(t-1)f_{ww}(t-1) + \frac{1}{2}m_{Xw}(t-1)f_{Xw}(t-1) + \frac{1}{2}m_{Xw}(t-1)f_{XX}(t-1) + m_{XX}(t-1)f_{Xw}(t-1) + \frac{1}{2}m_{XX}(t-1)f_{Xw}(t-1)\Big]/\tilde{\Omega}, \tag{2}$$

$$m_{XX}(t) = \frac{\Omega_{XX}}{2}\Big[\frac{1}{4}m_{Xw}(t-1)f_{Xw}(t-1) + \frac{1}{2}m_{Xw}(t-1)f_{XX}(t-1) + \frac{1}{2}m_{XX}f_{Xw}(t-1) + m_{XX}(t-1)f_{XX}(t-1)\Big]/\tilde{\Omega}, \tag{3}$$

$$f_{ww}(t) = \frac{1}{2}\Big[m_{ww}(t-1)f_{ww}(t-1) + \frac{1}{2}m_{ww}(t-1)f_{Xw}(t-1) + \frac{1}{2}m_{Xw}(t-1)f_{ww}(t-1) + \frac{1}{4}m_{Xw}(t-1)f_{Xw}(t-1)\Big]/\tilde{\Omega}, \tag{4}$$

$$f_{Xw}(t) = \frac{\Omega_{Xw}}{2}\Big[\frac{1}{2}m_{ww}(t-1)f_{Xw}(t-1) + m_{ww}(t-1)f_{XX}(t-1) + \frac{1}{2}m_{Xw}(t-1)f_{ww}(t-1) + \frac{1}{2}m_{Xw}(t-1)f_{Xw}(t-1) + \frac{1}{2}m_{Xw}(t-1)f_{XX}(t-1) + m_{XX}(t-1)f_{Xw}(t-1) + \frac{1}{2}m_{XX}(t-1)f_{Xw}(t-1)\Big]/\tilde{\Omega}, \tag{5}$$

$$f_{XX}(t) = \frac{\Omega_{XX}}{2}\Big[\frac{1}{4}m_{Xw}(t-1)f_{Xw}(t-1) + \frac{1}{2}m_{Xw}(t-1)f_{XX}(t-1) + \frac{1}{2}m_{XX}f_{Xw}(t-1) + m_{XX}(t-1)f_{XX}(t-1)\Big]/\tilde{\Omega}, \tag{6}$$

in which $m$ and $f$ represent males and females of the genotype indicated in their respective subscripts ($ww$ denoting wild-type, $Xw$ transgene heterozygote and $XX$ transgene homozygote) and $\Omega$ is the fitness of a given genotype (in subscript) relative to wild-type - assumed equal for both sexes. The overall fitness of the entire population $\tilde{\Omega}$ is calculated as the sum of all numerators in equations (1)-(6) and is used as a normalising factor to ensure genotype frequencies fill the entire range from zero to one.

To fit this model to experimental data (main text) we take a simple least squares regression approach. In particular we simultaneously fit relative fitness parameters for transgene heterozygous ($\Omega_{Xw}$) and homozygous ($\Omega_{XX}$) individuals by minimising the total error between two numerical simulations of the mathematical model representing the different single transgene experimental scenarios (fitting to the mean of the two experimental replicates). We model the scenario in which the initial cage setup consists of transgene heterozygous females and wild-type males via initial conditions of the form

$$m_{ww}(0) = 0.5, \quad m_{Xw}(0) = 0, \quad m_{XX}(0) = 0, \quad f_{ww}(0) = 0, \quad f_{Xw}(0) = 0.5, \quad f_{XX}(0) = 0,$$

and the case in which the initial cage consists of transgene heterozygous males and females via initial conditions of the form

$$m_{ww}(0) = 0, \quad m_{Xw}(0) = 0.5, \quad m_{XX}(0) = 0, \quad f_{ww}(0) = 0, \quad f_{Xw}(0) = 0.5, \quad f_{XX}(0) = 0.$$

The mathematical model presented here is simulated across a discretised parameter grid representing the full range of possible parameter vales for the relative fitness of transgene heterozygote and transgene homozygote individuals, with the sum of squared errors between the numerical simulation and mean experimental results calculated at each point. The parameter combination giving the lowest squared error is the deemed to provide the best fit and is thus carried forward into other areas of investigation. Note that to provide a fair comparison to experimental data, mean squared errors are based on transgene carrier frequencies - calculated from our modelling results as the sum of $Xw$ and $XX$ genotype frequencies. All parameter fitting was performed using Matlab (version R2019a; The MathWorks Inc., Natick, MA).

#### S4.2 Element B

Applying the procedure outlined above to the experimental data for cages containing only transgenic construct B (*bgcn*-Cas9) gives the results in Figure S1. This provides a best fit to the experimental data where B transgene heterozygotes have relative fitness $\Omega_{Xw}$=1 (i.e. no fitness cost) and B transgene homozygotes have relative fitness $\Omega_{XX}$=0.79 (i.e. a fitness cost of 21%). In practice this means there is a significant fitness cost associated with being homozygous for transgenic construct B but no cost to being heterozygous (i.e. they display full wild-type fitness).

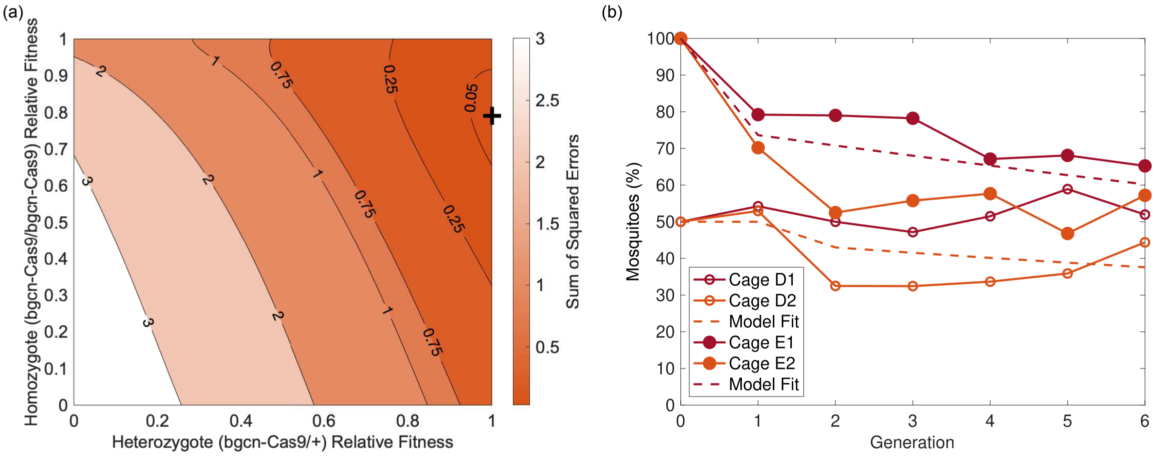


Figure S1: **Results of model fitting to experimental data from cage trials containing transgenic construct B (*bgcn*-Cas9) only.** (a) Sum of squared errors between numerical simulations and the mean of experimental data for the full range of relative fitness parameters. Here the colour map represents the sum of squared errors with contour lines for certain values included to provide visual clarity. The black cross indicates the parameter combination providing the minimal sum of squared errors. (b) Visualisations of the optimal fit obtained between the mathematical model (dashed lines) and experimental data (solid lines) for the case in which B transgene heterozygous females and wild-type males are used to setup the cage (blue) and where B transgene heterozygote males and females were used to setup the cage (red).

#### S4.3 Element A

A similar procedure was followed for cages containing only transgenic construct A ($kmo^{\text{gRNAs}}$). This gives the results in Figure S2, with the best fit being obtained where A transgene heterozygotes have relative fitness $\Omega_{Xw}$=1 (i.e. no fitness cost) and A transgene homozygotes have relative fitness $\Omega_{XX}$=0.81 (i.e. a fitness cost of 19%). As in the case of the B element, here A construct homozygotes display a fairly significant fitness cost - albeit smaller cost than the B element - while heterozygotes display equal fitness to wild-type individuals.

#### S4.4 Predicting Split-Drive Dynamics

The previous sections used population genetics mathematical models and least squares regression techniques to obtain estimated relative fitness parameters for B+/++, BB/++, ++/A+ and ++/AA genotypes; i.e. those in which only one transgenic construct is present and therefore will not undergo any Cas9-based cleavage nor the resulting homology directed or end-joining repair mechanisms. Since

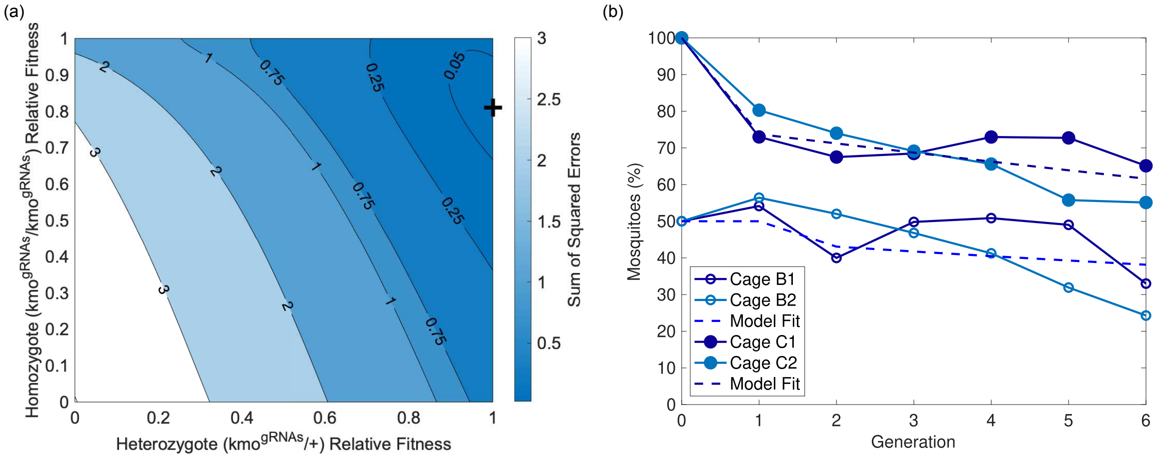


Figure S2: **Results of model fitting to experimental data from cage trials containing transgenic construct A (*kmo*$^{\text{gRNAs}}$) only.** (a) Sum of squared errors between numerical simulations and experimental data for all relative fitness parameters considered here. Here the colour map represents the sum of squared errors with contour lines for certain values included to provide visual clarity. The black cross indicates the parameter combination providing the minimal sum of squared errors. (b) Visualisations of the optimal fit obtained between the mathematical model (dashed lines) and experimental data (solid lines) for cases in which B transgene heterozygous females and wild-type males are used to setup the cage (blue) or where B transgene heterozygote males and females were used to setup the cage (red).

experimental work (in the main text) has directly measured homing rate parameters in both males and females, we simply require estimated relative fitness parameters for genotypes in which both the B and A transgenic constructs are present (i.e. B+/A+, B+/AA, BB/A+ and BB/AA). Comparing additive and multiplicative combinations of those relative fitness parameters obtained for B+/++, BB/++, ++/A+ and ++/AA genotypes within the population genetics model outlined in Section 2.1 reveals a slightly smaller sum of squared errors between the mean of experimental data for carrier frequencies of only B or A and those carrying both B and A for a model considering additive fitness costs. This results in relative fitness parameters $\Omega_{B+/A+} = 1$ (no fitness cost), $\Omega_{B+/AA} = 0.81$ (19% fitness cost), $\Omega_{BB/A+} = 0.79$ (21% fitness cost) and $\Omega_{BB/AA} = 0.60$ (40% fitness cost). In practise the restriction to additive or multiplicative fitness costs means only the relative fitness parameter for the BB/AA genotype displays any difference between the two cases due to the earlier parameter fittings that found zero fitness costs for individuals heterozygous for either one of the transgenic constructs. Allowing all four of the relative fitness parameters for B+/A+, B+/AA, BB/A+ and BB/AA genotypes to take any value in the range zero to one produces a slightly improved fit to experimental data, however the resulting parameter values did not produce results with any intuitive explanation. Thus we favour the use of an additive combination of relative fitness parameters obtained earlier, producing the results in Figure S3.

##### S4.4.1 Split-Drive Mathematical Model

As in Section 1, we formulate here a deterministic population genetics mathematical model that considers an infinite, closed (no migration), panmictic (random mating) population with a total of nine different genotypes, namely ++/++ (wild-type), ++/A+, ++/AA, B+/++, B+/A+, B+/AA, BB/++,

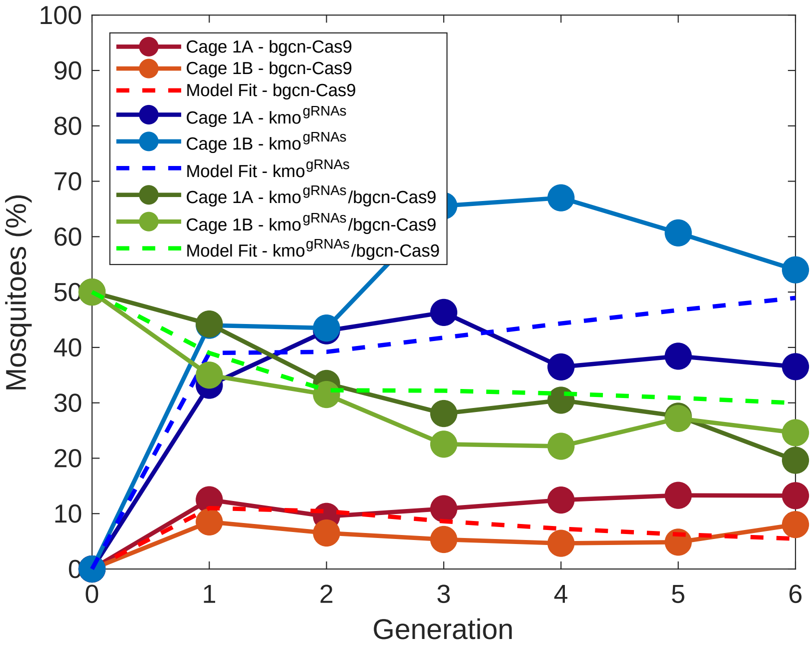


Figure S3: **Results of model fitting to experimental data from cage trials containing both split drive transgenic components (*kmo*$^{\text{gRNAs}}$ (A) and *bgcn*-Cas9(B)).** Here solid lines show experimental data for the percentage of mosquitoes carrying only the B (red) or A (blue) construct or both the B and A (green) constructs. Dashed lines show results from a population genetics mathematical model considering an additive combination of relative fitness parameters obtained using the single construct mathematical models in Section 1.

BB/A+ and BB/AA. This model is simulated using the following set of difference equations

$$
\begin{aligned}
m_{++/++} = \frac{\Omega_{++/++}}{2} \Big[ & m_{++/++}(t-1)f_{++/++}(t-1) + \frac{1}{2}m_{++/++}(t-1)f_{++/A+}(t-1) \\
& + \frac{1}{2}m_{++/++}(t-1)f_{B+/++}(t-1) + \frac{1}{4}m_{++/++}(t-1)f_{B+/A+}(t-1)(1-\Phi_F) \\
& + \frac{1}{2}m_{++/A+}(t-1)f_{++/++}(t-1) + \frac{1}{4}m_{++/A+}(t-1)f_{++/A+}(t-1) \\
& + \frac{1}{4}m_{++/A+}(t-1)f_{B+/++}(t-1) + \frac{1}{8}m_{++/A+}(t-1)f_{B+/A+}(t-1)(1-\Phi_F) \\
& + \frac{1}{2}m_{B+/++}(t-1)f_{++/++}(t-1) + \frac{1}{4}m_{B+/++}(t-1)f_{++/A+}(t-1) \\
& + \frac{1}{4}m_{B+/++}(t-1)f_{B+/++}(t-1) + \frac{1}{8}m_{B+/++}(t-1)f_{B+/A+}(t-1)(1-\Phi_F) \\
& + \frac{1}{4}m_{B+/A+}(t-1)(1-\Phi_M)f_{++/++}(t-1) + \frac{1}{8}m_{B+/A+}(t-1)(1-\Phi_M)f_{++/A+}(t-1) \\
& + \frac{1}{8}m_{B+/A+}(t-1)(1-\Phi_M)f_{B+/++}(t-1) + \frac{1}{16}m_{B+/A+}(t-1)(1-\Phi_M)f_{B+/A+}(t-1)(1-\Phi_F) \Big] / \tilde{\Omega},
\end{aligned} \tag{7}
$$

$$
\begin{aligned}
m_{++/A+} = \frac{\Omega_{++/A+}}{2} \Big[ & \frac{1}{2}m_{++/++}(t-1)f_{++/A+}(t-1) + m_{++/++}(t-1)f_{++/AA}(t-1) \\
& + \frac{1}{4}m_{++/++}(t-1)f_{B+/A+}(t-1)(1-\Phi_F) + \frac{1}{2}m_{++/++}(t-1)f_{B+/AA}(t-1)
\end{aligned} \tag{8}
$$

$$
\begin{aligned}
&+\frac{1}{2}m_{++/A+}(t-1)f_{++/++}(t-1)+\frac{1}{2}m_{++/A+}(t-1)f_{++/A+}(t-1)\\
&+\frac{1}{2}m_{++/A+}(t-1)f_{++/AA}(t-1)+\frac{1}{4}m_{++/A+}(t-1)f_{B+/++}(t-1)\\
&+\frac{1}{4}m_{++/A+}(t-1)f_{B+/A+}(t-1)(1-\Phi_F)+\frac{1}{4}m_{++/A+}(t-1)f_{B+/AA}(t-1)\\
&+m_{++/AA}(t-1)f_{++/++}(t-1)+\frac{1}{2}m_{++/AA}(t-1)f_{++/A+}(t-1)\\
&+\frac{1}{2}m_{++/AA}(t-1)f_{B+/++}(t-1)+\frac{1}{4}m_{++/AA}(t-1)f_{B+/A+}(t-1)(1-\Phi_F)\\
&+\frac{1}{4}m_{B+/++}(t-1)f_{++/A+}(t-1)+\frac{1}{2}m_{B+/++}(t-1)f_{++/AA}(t-1)\\
&+\frac{1}{8}m_{B+/++}(t-1)f_{B+/A+}(t-1)(1-\Phi_F)+\frac{1}{4}m_{B+/++}(t-1)f_{B+/AA}(t-1)\\
&+\frac{1}{4}m_{B+/A+}(t-1)(1-\Phi_M)f_{++/++}(t-1)+\frac{1}{4}m_{B+/A+}(t-1)(1-\Phi_M)f_{++/A+}(t-1)\\
&+\frac{1}{4}m_{B+/A+}(t-1)(1-\Phi_M)f_{++/AA}(t-1)+\frac{1}{8}m_{B+/A+}(t-1)(1-\Phi_M)f_{B+/++}(t-1)\\
&+\frac{1}{8}m_{B+/A+}(t-1)(1-\Phi_M)f_{B+/A+}(t-1)(1-\Phi_F)+\frac{1}{8}m_{B+/A+}(t-1)(1-\Phi_M)f_{B+/AA}(t-1)\\
&+\frac{1}{2}m_{B+/AA}(t-1)f_{++/++}(t-1)+\frac{1}{4}m_{B+/AA}(t-1)f_{++/A+}(t-1)\\
&+\frac{1}{4}m_{B+/AA}(t-1)f_{B+/++}(t-1)+\frac{1}{8}m_{B+/AA}(t-1)f_{B+/A+}(t-1)(1-\Phi_F)\\
&+\frac{1}{2}m_{++/++}(t-1)f_{B+/A+}(t-1)\Phi_F+\frac{1}{4}m_{++/A+}(t-1)f_{B+/A+}(t-1)\Phi_F\\
&+\frac{1}{4}m_{B+/++}(t-1)f_{B+/A+}(t-1)\Phi_F+\frac{1}{8}m_{B+/A+}(t-1)f_{B+/A+}(t-1)\Phi_F(1-\Phi_M)\\
&+\frac{1}{2}m_{B+/A+}(t-1)f_{++/++}(t-1)\Phi_M+\frac{1}{4}m_{B+/A+}(t-1)f_{++/A+}(t-1)\Phi_M\\
&+\frac{1}{4}m_{B+/A+}(t-1)f_{B+/++}(t-1)\Phi_M+\frac{1}{8}m_{B+/A+}(t-1)f_{B+/A+}(t-1)\Phi_M(1-\Phi_F)\\
&\left.+\ \frac{1}{8}(1-\Phi_M)m_{B+/A+}(t-1)\Phi_F f_{BB/A+}(t-1)\right]/\tilde{\Omega},
\end{aligned}
$$

$$
\begin{aligned}
m_{++/AA}=\frac{\Omega_{++/AA}}{2}&\left[\frac{1}{4}m_{++/A+}(t-1)f_{++/A+}(t-1)+\frac{1}{2}m_{++/A+}(t-1)f_{++/AA}(t-1)\right. \qquad (9)\\
&+\frac{1}{8}m_{++/A+}(t-1)f_{B+/A+}(t-1)(1-\Phi_F)+\frac{1}{4}m_{++/A+}(t-1)f_{B+/AA}(t-1)\\
&+\frac{1}{2}m_{++/AA}(t-1)f_{++/A+}(t-1)+m_{++/AA}(t-1)f_{++/AA}(t-1)\\
&+\frac{1}{4}m_{++/AA}(t-1)f_{B+/A+}(t-1)(1-\Phi_F)+\frac{1}{2}m_{++/AA}(t-1)f_{B+/AA}(t-1)\\
&+\frac{1}{8}m_{B+/A+}(t-1)(1-\Phi_M)f_{++/A+}(t-1)+\frac{1}{4}m_{B+/A+}(t-1)(1-\Phi_M)f_{++/AA}(t-1)\\
&+\frac{1}{16}m_{B+/A+}(t-1)(1-\Phi_M)f_{B+/A+}(t-1)(1-\Phi_F)+\frac{1}{8}m_{B+/A+}(t-1)(1-\Phi_M)f_{B+/AA}(t-1)\\
&+\frac{1}{4}m_{B+/AA}(t-1)f_{++/A+}(t-1)+\frac{1}{2}m_{B+/AA}(t-1)f_{++/AA}(t-1)\\
&+\frac{1}{8}m_{B+/AA}(t-1)f_{B+/A+}(t-1)(1-\Phi_F)+\frac{1}{4}m_{B+/AA}(t-1)f_{B+/AA}(t-1)\\
&+\frac{1}{4}m_{++/A+}(t-1)f_{B+/A+}(t-1)\Phi_F+\frac{1}{2}m_{++/AA}(t-1)f_{B+/A+}(t-1)\Phi_F\\
&+\frac{1}{8}m_{B+/A+}(t-1)f_{B+/A+}(t-1)\Phi_F(1-\Phi_M)+\frac{1}{4}m_{B+/AA}(t-1)f_{B+/A+}(t-1)\Phi_F\\
&+\frac{1}{4}m_{B+/A+}(t-1)f_{++/A+}(t-1)\Phi_M+\frac{1}{2}m_{B+/A+}(t-1)f_{++/AA}(t-1)\Phi_M\\
&+\frac{1}{8}m_{B+/A+}(t-1)f_{B+/A+}(t-1)\Phi_M(1-\Phi_F)+\frac{1}{4}m_{B+/A+}(t-1)f_{B+/AA}(t-1)\Phi_M
\end{aligned}
$$

$$+\frac{1}{4}m_{B+/A+}(t-1)f_{B+/A+}(t-1)\Phi_M\Phi_F+\frac{1}{8}(1-\Phi_M)m_{B+/A+}(t-1)\Phi_F f_{BB/A+}(t-1)\Big]/\tilde{\Omega},$$

$$m_{B+/++}=\frac{\Omega_{B+/++}}{2}\Big[\frac{1}{2}m_{++/++}(t-1)f_{B+/++}(t-1)+\frac{1}{4}m_{++/++}(t-1)f_{B+/A+}(t-1)(1-\Phi_F) \quad (10)$$

$$+m_{++/++}(t-1)f_{BB/++}(t-1)+\frac{1}{2}m_{++/++}(t-1)f_{BB/A+}(t-1)(1-\Phi_F)$$

$$+\frac{1}{4}m_{++/A+}(t-1)f_{B+/++}(t-1)+\frac{1}{8}m_{++/A+}(t-1)f_{B+/A+}(t-1)(1-\Phi_F)$$

$$+\frac{1}{2}m_{++/A+}(t-1)f_{BB/++}(t-1)+\frac{1}{4}m_{++/A+}(t-1)f_{BB/A+}(t-1)(1-\Phi_F)$$

$$+\frac{1}{2}m_{B+/++}(t-1)f_{++/++}(t-1)+\frac{1}{4}m_{B+/++}(t-1)f_{++/A+}(t-1)$$

$$+\frac{1}{2}m_{B+/++}(t-1)f_{B+/++}(t-1)+\frac{1}{4}m_{B+/++}(t-1)f_{B+/A+}(t-1)(1-\Phi_F)$$

$$+\frac{1}{2}m_{B+/++}(t-1)f_{BB/++}(t-1)+\frac{1}{4}m_{B+/++}(t-1)f_{BB/A+}(t-1)(1-\Phi_F)$$

$$+\frac{1}{4}m_{B+/A+}(t-1)(1-\Phi_M)f_{++/++}(t-1)+\frac{1}{8}m_{B+/A+}(t-1)(1-\Phi_M)f_{++/A+}(t-1)$$

$$+\frac{1}{4}m_{B+/A+}(t-1)(1-\Phi_M)f_{B+/++}(t-1)+\frac{1}{8}m_{B+/A+}(t-1)(1-\Phi_M)f_{B+/A+}(t-1)(1-\Phi_F)$$

$$+\frac{1}{4}m_{B+/A+}(t-1)(1-\Phi_M)f_{BB/++}(t-1)+\frac{1}{8}m_{B+/A+}(t-1)(1-\Phi_M)f_{BB/A+}(t-1)(1-\Phi_F)$$

$$+m_{BB/++}(t-1)f_{++/++}(t-1)+\frac{1}{2}m_{BB/++}(t-1)f_{++/A+}(t-1)$$

$$+\frac{1}{2}m_{BB/++}(t-1)f_{B+/++}(t-1)+\frac{1}{4}m_{BB/++}(t-1)f_{B+/A+}(t-1)(1-\Phi_F)$$

$$+\frac{1}{2}m_{BB/A+}(t-1)(1-\Phi_M)f_{++/++}(t-1)+\frac{1}{4}m_{BB/A+}(t-1)(1-\Phi_M)f_{++/A+}(t-1)$$

$$+\frac{1}{4}m_{BB/A+}(t-1)(1-\Phi_M)f_{B+/++}(t-1)+\frac{1}{8}m_{BB/A+}(t-1)(1-\Phi_M)f_{B+/A+}(t-1)(1-\Phi_F)\Big]/\tilde{\Omega},$$

$$m_{B+/A+}=\frac{\Omega_{B+/A+}}{2}\Big[\frac{1}{4}m_{++/++}(t-1)f_{B+/A+}(t-1)(1-\Phi_F)+\frac{1}{2}m_{++/++}(t-1)f_{B+/AA}(t-1) \quad (11)$$

$$+\frac{1}{2}m_{++/++}(t-1)f_{BB/A+}(t-1)(1-\Phi_F)+m_{++/++}(t-1)f_{BB/AA}(t-1)$$

$$+\frac{1}{4}m_{++/A+}(t-1)f_{B+/++}(t-1)+\frac{1}{4}m_{++/A+}(t-1)f_{B+/A+}(t-1)(1-\Phi_F)$$

$$+\frac{1}{4}m_{++/A+}(t-1)f_{B+/AA}(t-1)+\frac{1}{2}m_{++/A+}(t-1)f_{BB/++}(t-1)$$

$$+\frac{1}{2}m_{++/A+}(t-1)f_{BB/A+}(t-1)(1-\Phi_F)+\frac{1}{2}m_{++/A+}(t-1)f_{BB/AA}(t-1)$$

$$+\frac{1}{2}m_{++/AA}(t-1)f_{B+/++}(t-1)+\frac{1}{4}m_{++/AA}(t-1)f_{B+/A+}(t-1)(1-\Phi_F)$$

$$+m_{++/AA}(t-1)f_{BB/++}(t-1)+\frac{1}{2}m_{++/AA}(t-1)f_{BB/A+}(t-1)(1-\Phi_F)$$

$$+\frac{1}{4}m_{B+/++}(t-1)f_{++/A+}(t-1)+\frac{1}{2}m_{B+/++}(t-1)f_{++/AA}(t-1)$$

$$+\frac{1}{4}m_{B+/++}(t-1)f_{B+/A+}(t-1)(1-\Phi_F)+\frac{1}{2}m_{B+/++}(t-1)f_{B+/AA}(t-1)$$

$$+\frac{1}{4}m_{B+/++}(t-1)f_{BB/A+}(t-1)(1-\Phi_F)+\frac{1}{2}m_{B+/++}(t-1)f_{BB/AA}(t-1)$$

$$+\frac{1}{4}m_{B+/A+}(t-1)(1-\Phi_M)f_{++/++}(t-1)+\frac{1}{4}m_{B+/A+}(t-1)(1-\Phi_M)f_{++/A+}(t-1)$$

$$+\frac{1}{4}m_{B+/A+}(t-1)(1-\Phi_M)f_{++/AA}(t-1)+\frac{1}{4}m_{B+/A+}(t-1)(1-\Phi_M)f_{B+/++}(t-1)$$

$$+\frac{1}{4}m_{B+/A+}(t-1)(1-\Phi_M)f_{B+/A+}(t-1)(1-\Phi_F)+\frac{1}{4}m_{B+/A+}(t-1)(1-\Phi_M)f_{B+/AA}(t-1)$$

$$
\begin{aligned}
&+\frac{1}{4}m_{B+/A+}(t-1)(1-\Phi_M)f_{BB/++}(t-1)+\frac{1}{4}m_{B+/A+}(t-1)(1-\Phi_M)f_{BB/A+}(t-1)(1-\Phi_F)\\
&+\frac{1}{4}m_{B+/A+}(t-1)(1-\Phi_M)f_{BB/AA}(t-1)+\frac{1}{2}m_{B+/AA}(t-1)f_{++/++}(t-1)\\
&+\frac{1}{4}m_{B+/AA}(t-1)f_{++/A+}(t-1)+\frac{1}{2}m_{B+/AA}(t-1)f_{B+/++}(t-1)\\
&+\frac{1}{4}m_{B+/AA}(t-1)f_{B+/A+}(t-1)(1-\Phi_F)+\frac{1}{2}m_{B+/AA}(t-1)f_{BB/++}(t-1)\\
&+\frac{1}{4}m_{B+/AA}(t-1)f_{BB/A+}(t-1)(1-\Phi_F)+\frac{1}{2}m_{BB/++}(t-1)f_{++/A+}(t-1)\\
&+m_{BB/++}(t-1)f_{++/AA}(t-1)+\frac{1}{4}m_{BB/++}(t-1)f_{B+/A+}(t-1)(1-\Phi_F)\\
&+\frac{1}{2}m_{BB/++}(t-1)f_{B+/AA}(t-1)+\frac{1}{2}m_{BB/A+}(t-1)(1-\Phi_M)f_{++/++}(t-1)\\
&+\frac{1}{2}m_{BB/A+}(t-1)(1-\Phi_M)f_{++/A+}(t-1)+\frac{1}{2}m_{BB/A+}(t-1)(1-\Phi_M)f_{++/AA}(t-1)\\
&+\frac{1}{4}m_{BB/A+}(t-1)(1-\Phi_M)f_{B+/++}(t-1)+\frac{1}{4}m_{BB/A+}(t-1)(1-\Phi_M)f_{B+/A+}(t-1)(1-\Phi_F)\\
&+\frac{1}{4}m_{BB/A+}(t-1)(1-\Phi_M)f_{B+/AA}(t-1)+m_{BB/AA}(t-1)f_{++/++}(t-1)\\
&+\frac{1}{2}m_{BB/AA}(t-1)f_{++/A+}(t-1)+\frac{1}{2}m_{BB/AA}(t-1)f_{B+/++}(t-1)\\
&+\frac{1}{4}m_{BB/AA}(t-1)f_{B+/A+}(t-1)(1-\Phi_F)+\frac{1}{2}m_{++/++}(t-1)f_{B+/A+}(t-1)\Phi_F\\
&+m_{++/++}(t-1)f_{BB/A+}(t-1)\Phi_F+\frac{1}{4}m_{++/A+}(t-1)f_{B+/A+}(t-1)\Phi_F\\
&+\frac{1}{2}m_{++/A+}(t-1)f_{BB/A+}(t-1)\Phi_F+\frac{1}{2}m_{B+/++}(t-1)f_{B+/A+}(t-1)\Phi_F\\
&+\frac{1}{2}m_{B+/++}(t-1)f_{BB/A+}(t-1)\Phi_F+\frac{1}{4}m_{B+/A+}(t-1)f_{B+/A+}(t-1)\Phi_F(1-\Phi_M)\\
&+\frac{1}{4}m_{B+/A+}(t-1)f_{BB/A+}(t-1)\Phi_F(1-\Phi_M)+\frac{1}{2}m_{BB/++}(t-1)f_{B+/A+}(t-1)\Phi_F\\
&+\frac{1}{4}m_{BB/A+}(t-1)f_{B+/A+}(t-1)\Phi_F(1-\Phi_M)+\frac{1}{2}m_{B+/A+}(t-1)f_{++/++}(t-1)\Phi_M\\
&+\frac{1}{4}m_{B+/A+}(t-1)f_{++/A+}(t-1)\Phi_M+\frac{1}{2}m_{B+/A+}(t-1)f_{B+/++}(t-1)\Phi_M\\
&+\frac{1}{4}m_{B+/A+}(t-1)f_{B+/A+}(t-1)\Phi_M(1-\Phi_F)+\frac{1}{2}m_{B+/A+}(t-1)f_{BB/++}(t-1)\Phi_M\\
&+\frac{1}{4}m_{B+/A+}(t-1)f_{BB/A+}(t-1)\Phi_M(1-\Phi_F)+m_{BB/A+}(t-1)f_{++/++}(t-1)\Phi_M\\
&+\frac{1}{2}m_{BB/A+}(t-1)f_{++/A+}(t-1)\Phi_M+\frac{1}{2}m_{BB/A+}(t-1)f_{B+/++}(t-1)\Phi_M\\
&\left.+\frac{1}{4}m_{BB/A+}(t-1)f_{B+/A+}(t-1)\Phi_M(1-\Phi_F)\right]/\tilde{\Omega},
\end{aligned}
$$

$$
\begin{aligned}
m_{B+/AA}=\frac{\Omega_{B+/AA}}{2}&\left[\frac{1}{8}m_{++/A+}(t-1)f_{B+/A+}(t-1)(1-\Phi_F)+\frac{1}{4}m_{++/A+}(t-1)f_{B+/AA}(t-1)\right.\\
&+\frac{1}{4}m_{++/A+}(t-1)f_{BB/A+}(t-1)(1-\Phi_F)+\frac{1}{2}m_{++/A+}(t-1)f_{BB/AA}(t-1)\\
&+\frac{1}{4}m_{++/AA}(t-1)f_{B+/A+}(t-1)(1-\Phi_F)+\frac{1}{2}m_{++/AA}(t-1)f_{B+/AA}(t-1)\\
&+\frac{1}{2}m_{++/AA}(t-1)f_{BB/A+}(t-1)(1-\Phi_F)+m_{++/AA}(t-1)f_{BB/AA}(t-1)\\
&+\frac{1}{8}m_{B+/A+}(t-1)(1-\Phi_M)f_{++/A+}(t-1)+\frac{1}{4}m_{B+/A+}(t-1)(1-\Phi_M)f_{++/AA}(t-1)\\
&+\frac{1}{8}m_{B+/A+}(t-1)(1-\Phi_M)f_{B+/A+}(t-1)(1-\Phi_F)+\frac{1}{4}m_{B+/A+}(t-1)(1-\Phi_M)f_{B+/AA}(t-1)
\end{aligned}
\qquad (12)
$$

$$
\begin{aligned}
&+\frac{1}{8}m_{B+/A+}(t-1)(1-\Phi_M)f_{BB/A+}(t-1)(1-\Phi_F)+\frac{1}{4}m_{B+/A+}(t-1)(1-\Phi_M)f_{BB/AA}(t-1)\\
&+\frac{1}{4}m_{B+/AA}(t-1)f_{++/A+}(t-1)+\frac{1}{2}m_{B+/AA}(t-1)f_{++/AA}(t-1)\\
&+\frac{1}{4}m_{B+/AA}(t-1)f_{B+/A+}(t-1)(1-\Phi_F)+\frac{1}{2}m_{B+/AA}(t-1)f_{B+/AA}(t-1)\\
&+\frac{1}{4}m_{B+/AA}(t-1)f_{BB/A+}(t-1)(1-\Phi_F)+\frac{1}{2}m_{B+/AA}(t-1)f_{BB/AA}(t-1)\\
&+\frac{1}{4}m_{BB/A+}(t-1)(1-\Phi_M)f_{++/A+}(t-1)+\frac{1}{2}m_{BB/A+}(t-1)(1-\Phi_M)f_{++/AA}(t-1)\\
&+\frac{1}{8}m_{BB/A+}(t-1)(1-\Phi_M)f_{B+/A+}(t-1)(1-\Phi_F)+\frac{1}{4}m_{BB/A+}(t-1)(1-\Phi_M)f_{B+/AA}(t-1)\\
&+\frac{1}{2}m_{BB/AA}(t-1)f_{++/A+}(t-1)+m_{BB/AA}(t-1)f_{++/AA}(t-1)\\
&+\frac{1}{4}m_{BB/AA}(t-1)f_{B+/A+}(t-1)(1-\Phi_F)+\frac{1}{2}m_{BB/AA}(t-1)f_{B+/AA}(t-1)\\
&+\frac{1}{4}m_{++/A+}(t-1)f_{B+/A+}(t-1)\Phi_F+\frac{1}{2}m_{++/A+}(t-1)f_{BB/A+}(t-1)\Phi_F\\
&+\frac{1}{2}m_{++/AA}(t-1)f_{B+/A+}(t-1)\Phi_F+m_{++/AA}(t-1)f_{BB/A+}(t-1)\Phi_F\\
&+\frac{1}{4}m_{B+/A+}(t-1)f_{B+/A+}(t-1)\Phi_F(1-\Phi_M)+\frac{1}{4}m_{B+/A+}(t-1)f_{BB/A+}(t-1)\Phi_F(1-\Phi_M)\\
&+\frac{1}{2}m_{B+/AA}(t-1)f_{B+/A+}(t-1)\Phi_F+\frac{1}{2}m_{B+/AA}(t-1)f_{BB/A+}(t-1)\Phi_F\\
&+\frac{1}{4}m_{BB/A+}(t-1)f_{B+/A+}(t-1)\Phi_F(1-\Phi_M)+\frac{1}{2}m_{BB/AA}(t-1)f_{B+/A+}(t-1)\Phi_F\\
&+\frac{1}{4}m_{B+/A+}(t-1)f_{++/A+}(t-1)\Phi_M+\frac{1}{2}m_{B+/A+}(t-1)f_{++/AA}(t-1)\Phi_M\\
&+\frac{1}{4}m_{B+/A+}(t-1)f_{B+/A+}(t-1)\Phi_M(1-\Phi_F)+\frac{1}{2}m_{B+/A+}(t-1)f_{B+/AA}(t-1)\Phi_M\\
&+\frac{1}{4}m_{B+/A+}(t-1)f_{BB/A+}(t-1)\Phi_M(1-\Phi_F)+\frac{1}{2}m_{B+/A+}(t-1)f_{BB/AA}(t-1)\Phi_M\\
&+\frac{1}{2}m_{BB/A+}(t-1)f_{++/A+}(t-1)\Phi_M+m_{BB/A+}(t-1)f_{++/AA}(t-1)\Phi_M\\
&+\frac{1}{4}m_{BB/A+}(t-1)f_{B+/A+}(t-1)\Phi_M(1-\Phi_F)+\frac{1}{2}m_{BB/A+}(t-1)f_{B+/AA}(t-1)\Phi_M\\
&+\frac{1}{2}m_{B+/A+}(t-1)f_{B+/A+}(t-1)\Phi_M\Phi_F+\frac{1}{2}m_{B+/A+}(t-1)f_{BB/A+}(t-1)\Phi_M\Phi_F\\
&+\frac{1}{2}m_{BB/A+}(t-1)f_{B+/A+}(t-1)\Phi_M\Phi_F\Big]/\tilde{\Omega},
\end{aligned}
$$

$$
\begin{aligned}
m_{BB/++}=\frac{\Omega_{BB/++}}{2}&\Big[\frac{1}{4}m_{B+/++}(t-1)f_{B+/++}(t-1)+\frac{1}{8}m_{B+/++}(t-1)f_{B+/A+}(t-1)(1-\Phi_F) \qquad (13)\\
&+\frac{1}{2}m_{B+/++}(t-1)f_{BB/++}(t-1)+\frac{1}{4}m_{B+/++}(t-1)f_{BB/A+}(t-1)(1-\Phi_F)\\
&+\frac{1}{8}m_{B+/A+}(t-1)(1-\Phi_M)f_{B+/++}(t-1)+\frac{1}{16}m_{B+/A+}(t-1)(1-\Phi_M)f_{B+/A+}(t-1)(1-\Phi_F)\\
&+\frac{1}{4}m_{B+/A+}(t-1)(1-\Phi_M)f_{BB/++}(t-1)+\frac{1}{8}m_{B+/A+}(t-1)(1-\Phi_M)f_{BB/A+}(t-1)(1-\Phi_F)\\
&+\frac{1}{2}m_{BB/++}(t-1)f_{B+/++}(t-1)+\frac{1}{4}m_{BB/++}(t-1)f_{B+/A+}(t-1)(1-\Phi_F)\\
&+m_{BB/++}(t-1)f_{BB/++}(t-1)+\frac{1}{2}m_{BB/++}(t-1)f_{BB/A+}(t-1)(1-\Phi_F)\\
&+\frac{1}{4}m_{BB/A+}(t-1)(1-\Phi_M)f_{B+/++}(t-1)+\frac{1}{8}m_{BB/A+}(t-1)(1-\Phi_M)f_{B+/A+}(t-1)(1-\Phi_F)\\
&+\frac{1}{2}m_{BB/A+}(t-1)(1-\Phi_M)f_{BB/++}(t-1)+\frac{1}{4}m_{BB/A+}(t-1)(1-\Phi_M)f_{BB/A+}(t-1)(1-\Phi_F)\Big]/\tilde{\Omega},
\end{aligned}
$$

$$
\begin{aligned}
m_{BB/A+} = \frac{\Omega_{BB/A+}}{2} \Bigg[ & \frac{1}{8} m_{B+/++}(t-1) f_{B+/A+}(t-1)(1-\Phi_F) + \frac{1}{4} m_{B+/++}(t-1) f_{B+/AA}(t-1) \\
& + \frac{1}{4} m_{B+/++}(t-1) f_{BB/A+}(t-1)(1-\Phi_F) + \frac{1}{2} m_{B+/++}(t-1) f_{BB/AA}(t-1) \\
& + \frac{1}{8} m_{B+/A+}(t-1)(1-\Phi_M) f_{B+/++}(t-1) + \frac{1}{8} m_{B+/A+}(t-1)(1-\Phi_M) f_{B+/A+}(t-1)(1-\Phi_F) \\
& + \frac{1}{8} m_{B+/A+}(t-1)(1-\Phi_M) f_{B+/AA}(t-1) + \frac{1}{4} m_{B+/A+}(t-1)(1-\Phi_M) f_{BB/++}(t-1) \\
& + \frac{1}{4} m_{B+/A+}(t-1)(1-\Phi_M) f_{BB/A+}(t-1)(1-\Phi_F) + \frac{1}{4} m_{B+/A+}(t-1)(1-\Phi_M) f_{BB/AA}(t-1) \\
& + \frac{1}{4} m_{B+/AA}(t-1) f_{B+/++}(t-1) + \frac{1}{8} m_{B+/AA}(t-1) f_{B+/A+}(t-1)(1-\Phi_F) \\
& + \frac{1}{2} m_{B+/AA}(t-1) f_{BB/++}(t-1) + \frac{1}{4} m_{B+/AA}(t-1) f_{BB/A+}(t-1)(1-\Phi_F) \\
& + \frac{1}{4} m_{BB/++}(t-1) f_{B+/A+}(t-1)(1-\Phi_F) + \frac{1}{2} m_{BB/++}(t-1) f_{B+/AA}(t-1) \\
& + \frac{1}{2} m_{BB/++}(t-1) f_{BB/A+}(t-1)(1-\Phi_F) + m_{BB/++}(t-1) f_{BB/AA}(t-1) \\
& + \frac{1}{4} m_{BB/A+}(t-1)(1-\Phi_M) f_{B+/++}(t-1) + \frac{1}{4} m_{BB/A+}(t-1)(1-\Phi_M) f_{B+/A+}(t-1)(1-\Phi_F) \\
& + \frac{1}{4} m_{BB/A+}(t-1)(1-\Phi_M) f_{B+/AA}(t-1) + \frac{1}{2} m_{BB/A+}(t-1)(1-\Phi_M) f_{BB/++}(t-1) \\
& + \frac{1}{2} m_{BB/A+}(t-1)(1-\Phi_M) f_{BB/A+}(t-1)(1-\Phi_F) + \frac{1}{2} m_{BB/A+}(t-1)(1-\Phi_M) f_{BB/AA}(t-1) \\
& + \frac{1}{2} m_{BB/AA}(t-1) f_{B+/++}(t-1) + \frac{1}{4} m_{BB/AA}(t-1) f_{B+/A+}(t-1)(1-\Phi_F) \\
& + m_{BB/AA}(t-1) f_{BB/++}(t-1) + \frac{1}{2} m_{BB/AA}(t-1) f_{BB/A+}(t-1)(1-\Phi_F) \\
& + \frac{1}{4} m_{B+/++}(t-1) f_{B+/A+}(t-1)\Phi_F + \frac{1}{2} m_{B+/++}(t-1) f_{BB/A+}(t-1)\Phi_F \\
& + \frac{1}{8} m_{B+/A+}(t-1) f_{B+/A+}(t-1)\Phi_F(1-\Phi_M) + \frac{1}{8} m_{B+/A+}(t-1) f_{BB/A+}(t-1)\Phi_F(1-\Phi_M) \\
& + \frac{1}{2} m_{BB/++}(t-1) f_{B+/A+}(t-1)\Phi_F + m_{BB/++}(t-1) f_{BB/A+}(t-1)\Phi_F \\
& + \frac{1}{4} m_{BB/A+}(t-1) f_{B+/A+}(t-1)\Phi_F(1-\Phi_M) + \frac{1}{2} m_{BB/A+}(t-1) f_{BB/A+}(t-1)\Phi_F(1-\Phi_M) \\
& + \frac{1}{4} m_{B+/A+}(t-1) f_{B+/++}(t-1)\Phi_M + \frac{1}{8} m_{B+/A+}(t-1) f_{B+/A+}(t-1)\Phi_M(1-\Phi_F) \\
& + \frac{1}{2} m_{B+/A+}(t-1) f_{BB/++}(t-1)\Phi_M + \frac{1}{4} m_{B+/A+}(t-1) f_{BB/A+}(t-1)\Phi_M(1-\Phi_F) \\
& + \frac{1}{2} m_{BB/A+}(t-1) f_{B+/++}(t-1)\Phi_M + \frac{1}{4} m_{BB/A+}(t-1) f_{B+/A+}(t-1)\Phi_M(1-\Phi_F) \\
& + m_{BB/A+}(t-1) f_{BB/++}(t-1)\Phi_M + \frac{1}{2} m_{BB/A+}(t-1) f_{BB/A+}(t-1)\Phi_M(1-\Phi_F) \Bigg] / \tilde{\Omega},
\end{aligned} \tag{14}
$$

$$
\begin{aligned}
m_{BB/AA} = \frac{\Omega_{BB/AA}}{2} \Bigg[ & \frac{1}{16} m_{B+/A+}(t-1)(1-\Phi_M) f_{B+/A+}(t-1)(1-\Phi_F) \\
& + \frac{1}{8} m_{B+/A+}(t-1)(1-\Phi_M) f_{B+/AA}(t-1) + \frac{1}{8} m_{B+/A+}(t-1)(1-\Phi_M) f_{BB/A+}(t-1)(1-\Phi_F) \\
& + \frac{1}{4} m_{B+/A+}(t-1)(1-\Phi_M) f_{BB/AA}(t-1) + \frac{1}{8} m_{B+/AA}(t-1) f_{B+/A+}(t-1)(1-\Phi_F) \\
& + \frac{1}{4} m_{B+/AA}(t-1) f_{B+/AA}(t-1) + \frac{1}{4} m_{B+/AA}(t-1) f_{BB/A+}(t-1)(1-\Phi_F) \\
& + \frac{1}{2} m_{B+/AA}(t-1) f_{BB/AA}(t-1) + \frac{1}{8} m_{BB/A+}(t-1)(1-\Phi_M) f_{B+/A+}(t-1)(1-\Phi_F) \\
& + \frac{1}{4} m_{BB/A+}(t-1)(1-\Phi_M) f_{B+/AA}(t-1) + \frac{1}{4} m_{BB/A+}(t-1)(1-\Phi_M) f_{BB/A+}(t-1)(1-\Phi_F)
\end{aligned} \tag{15}
$$

$$
\begin{aligned}
&+\frac{1}{2}m_{BB/A+}(t-1)(1-\Phi_M)f_{BB/AA}(t-1)+\frac{1}{4}m_{BB/AA}(t-1)f_{B+/A+}(t-1)(1-\Phi_F)\\
&+\frac{1}{2}m_{BB/AA}(t-1)f_{B+/AA}(t-1)+\frac{1}{2}m_{BB/AA}(t-1)f_{BB/A+}(t-1)(1-\Phi_F)\\
&+m_{BB/AA}(t-1)f_{BB/AA}(t-1)+\frac{1}{8}m_{B+/A+}(t-1)f_{B+/A+}(t-1)\Phi_F(1-\Phi_M)\\
&+\frac{1}{8}m_{B+/A+}(t-1)f_{BB/A+}(t-1)\Phi_F(1-\Phi_M)+\frac{1}{4}m_{B+/AA}(t-1)f_{B+/A+}(t-1)\Phi_F\\
&+\frac{1}{2}m_{B+/AA}(t-1)f_{BB/A+}(t-1)\Phi_F+\frac{1}{4}m_{BB/A+}(t-1)f_{B+/A+}(t-1)\Phi_F(1-\Phi_M)\\
&+\frac{1}{2}m_{BB/A+}(t-1)f_{BB/A+}(t-1)\Phi_F(1-\Phi_M)+\frac{1}{2}m_{BB/AA}(t-1)f_{B+/A+}(t-1)\Phi_F\\
&+m_{BB/AA}(t-1)f_{BB/A+}(t-1)\Phi_F+\frac{1}{8}m_{B+/A+}(t-1)f_{B+/A+}(t-1)\Phi_M(1-\Phi_F)\\
&+\frac{1}{4}m_{B+/A+}(t-1)f_{B+/AA}(t-1)\Phi_M+\frac{1}{4}m_{B+/A+}(t-1)f_{BB/A+}(t-1)\Phi_M(1-\Phi_F)\\
&+\frac{1}{2}m_{B+/A+}(t-1)f_{BB/AA}(t-1)\Phi_M+\frac{1}{4}m_{BB/A+}(t-1)f_{B+/A+}(t-1)\Phi_M(1-\Phi_F)\\
&+\frac{1}{2}m_{BB/A+}(t-1)f_{B+/AA}(t-1)\Phi_M+\frac{1}{2}m_{BB/A+}(t-1)f_{BB/A+}(t-1)\Phi_M(1-\Phi_F)\\
&+m_{BB/A+}(t-1)f_{BB/AA}(t-1)\Phi_M+\frac{1}{4}m_{B+/A+}(t-1)f_{B+/A+}(t-1)\Phi_M\Phi_F\\
&+\frac{1}{2}m_{B+/A+}(t-1)f_{BB/A+}(t-1)\Phi_M\Phi_F+\frac{1}{2}m_{BB/A+}(t-1)f_{B+/A+}(t-1)\Phi_M\Phi_F\\
&+m_{BB/A+}(t-1)f_{BB/A+}(t-1)\Phi_M\Phi_F\big]/\tilde{\Omega},
\end{aligned}
$$

and nine identical equations for females of these genotypes. Here $m$ and $f$ represent males and females of the genotype indicated in their respective subscripts and $\Omega$ is the fitness of a given genotype (in subscript) relative to wild-type. The overall fitness of the entire population $\tilde{\Omega}$ is calculated as the sum of all numerators in equations (7)-(15) and is used as a normalising factor to ensure genotype frequencies fill the entire range from zero to one.

To predict the results of the split drive treatment cages (main text) we compare the sum of squared errors for B, A and B/A carriers to model results for additive or multiplicative combinations of the relative fitness parameters obtained in Section 1. These produced very similar goodness of fit and so an additive combination of fitness parameters was utilized since this yields a simple and intuitive explanation of fitness cost interactions. The particular model scenario considered is that in which the initial cage setup consists of females heterozygous for both transgenic constructs and wild-type males, leading to initial conditions of the form

$$
\begin{gathered}
m_{++/++}(0)=0.5,\quad m_{++/A+}(0)=0,\quad m_{++/AA}(0)=0,\quad m_{B+/++}(0)=0,\quad m_{B+/A+}(0)=0,\\
m_{B+/AA}(0)=0,\quad m_{BB/++}(0)=0,\quad m_{BB/A+}(0)=0,\quad m_{BB/AA}(0)=0,\\
f_{++/++}(0)=0,\quad f_{++/A+}(0)=0,\quad f_{++/AA}(0)=0,\quad f_{B+/++}(0)=0,\quad f_{B+/A+}(0)=0.5,\\
f_{B+/AA}(0)=0,\quad f_{BB/++}(0)=0,\quad f_{BB/A+}(0)=0,\quad f_{BB/AA}(0)=0.
\end{gathered}
$$

All numerical simulations of this model were performed using Matlab (version R2019a; The MathWorks Inc., Natick, MA).

#### S4.5 Split-Drive Stochastic Model

The deterministic model outlined above was used to (1) estimate relative fitness parameters for each transgenic construct and (2) to provide simple, initial predictions around the outcome of the split drive treatment cages. While useful in planning cage trials and to some extent in analyzing the final outcomes, such models do not account for the stochasticity inherent in such experiments. Thus, we also developed a stochastic model framework capable of providing an estimated range within which we would expect the experimental results to fall. This is largely based on a previously developed model framework used for the study of novel multiplexing approaches for CRISPR-based gene drives[1] and has been altered to capture the dynamics of a split drive system. The main basis of this model has been discussed in depth previously and so here we provide only a brief outline of the adjustments made.

Firstly, since the initial model was formulated for the study of multiplexing strategies, it was already capable of managing genetic inheritance across multiple loci. Rather than allowing a free choice of number of loci, here we eliminate this and fix the model to two loci (those for the A and B elements of the split drive).

Perhaps the most important alteration here is the switch from the initial model of embryonic homing to one considering germline homing - as observed in the split drive developed in this study. In the initial model, a homing module altered the assigned genotypes of 'eggs' resulting from each mating pair. To move to a model of germline homing, this homing module was simply moved such that it acts between the selection of parental mating pairs and the assignment of genotypes to the resulting 'eggs' from those mating pairs.

The final, and rather trivial, change required here was in the calculation of transgene carrier frequencies. In the initial model these were calculated in the context of multiplexing approaches where the presence of any gene drive construct was the important factor - thus resulting in a single transgene carrier frequency no matter the multiplex number. However, here it is important to track carrier frequencies of both the B and A elements of the split drive. The relevant calculations were adjusted accordingly, producing separate transgene carrier frequencies for the B and A elements.

As with the deterministic modelling, all stochastic numerical simulations were performed using Matlab (version R2019a; The MathWorks Inc., Natick, MA).

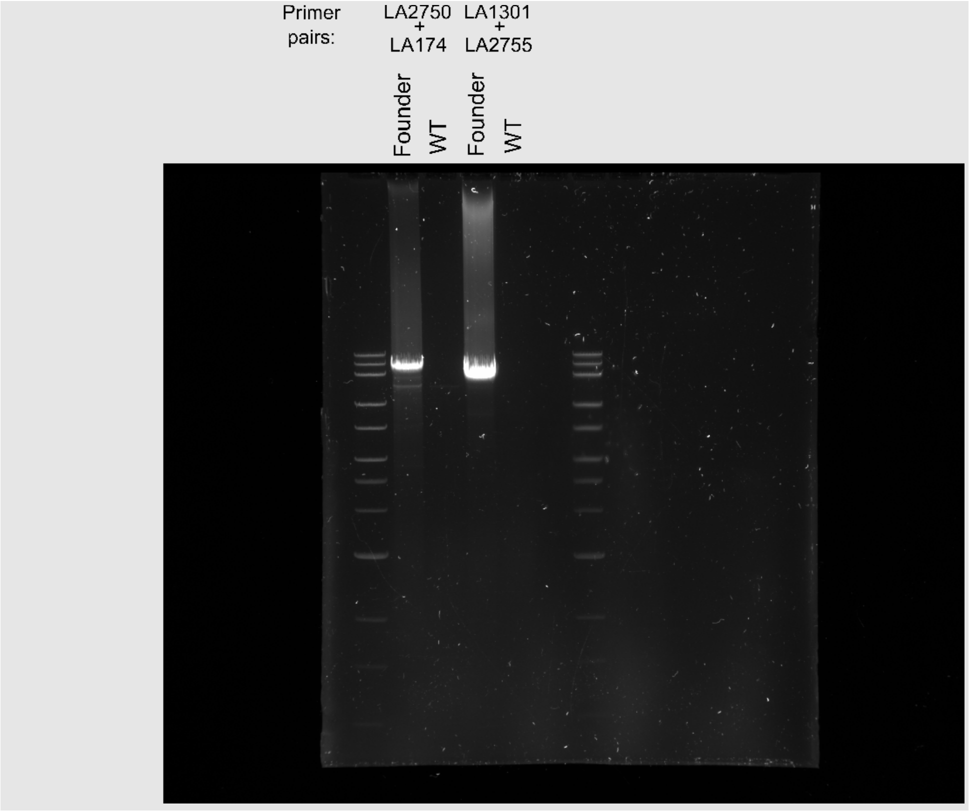


**Fig S4. Confirmation of *kmo*$^{sgRNAs}$ integration site by PCR.** Gel electrophoresis of PCRs to confirm the integration site of *kmo*$^{sgRNAs}$. The Quick Load 1kb Extend DNA ladder (NEB) was loaded into the first and last well.

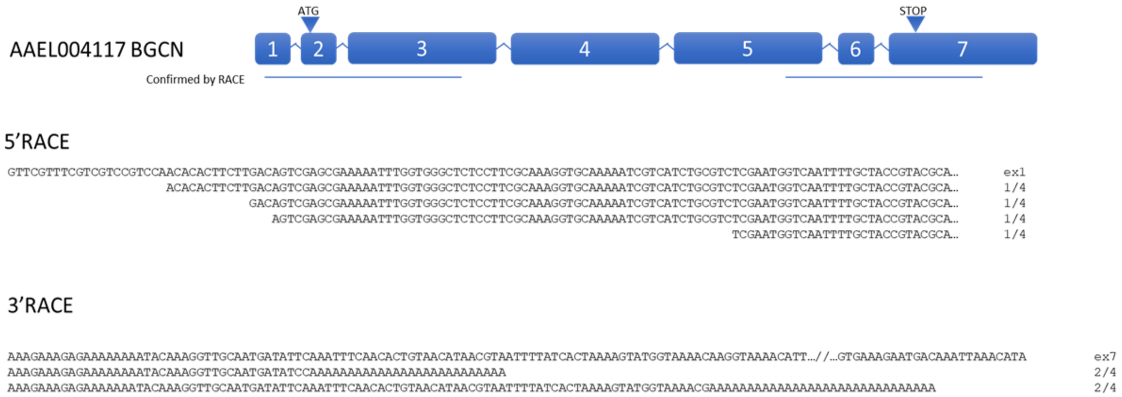


**Fig S5. RACE results for *bgcn*.** Diagram of the *bgcn* gene based on Vectorbase L5 assembly annotations (not to scale). RACE PCR products were cloned into the pJET vector and four clones were sequenced. The top line denotes the start of exon 1 or end of exon 7 as annotated, proceeding lines begin with the first base transcribed (pJET vector and RACE primer sequences trimmed). Right hand column denotes the number of clones with identical sequences.

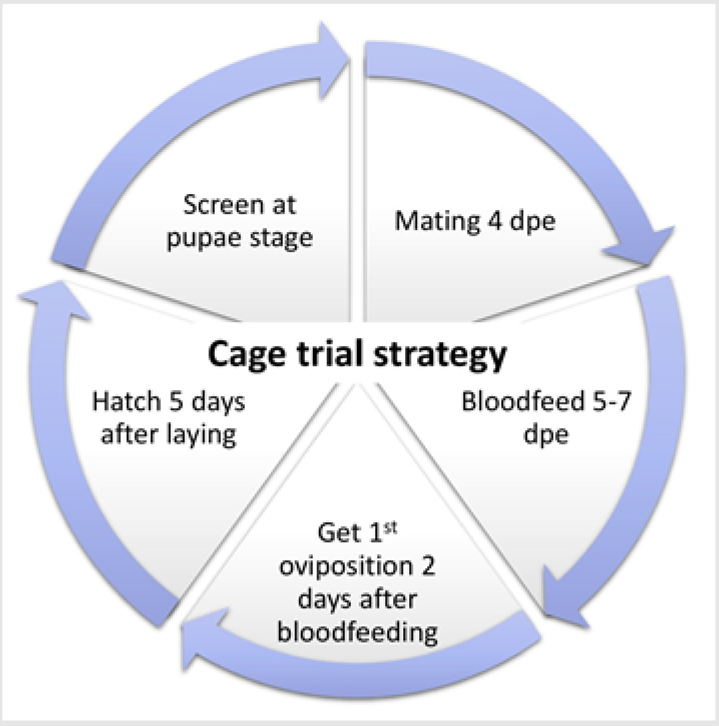


**Fig S6. Diagram of the cage trial strategy.** Steps followed during the 6 generations of the cage trial beginning with the mating of males and females 4 days post eclosion (dpe). Females were allowed to blood feed when they were 5-7 dpe and 2 days later eggs were collected. Five days after oviposition the eggs were hatched in degassed RO (reverse osmosis) water. 250 larvae for each cage were reared under standardized conditions. Finally, the progeny of each generation was screened at pupae stage for fluorescence and eye phenotype.

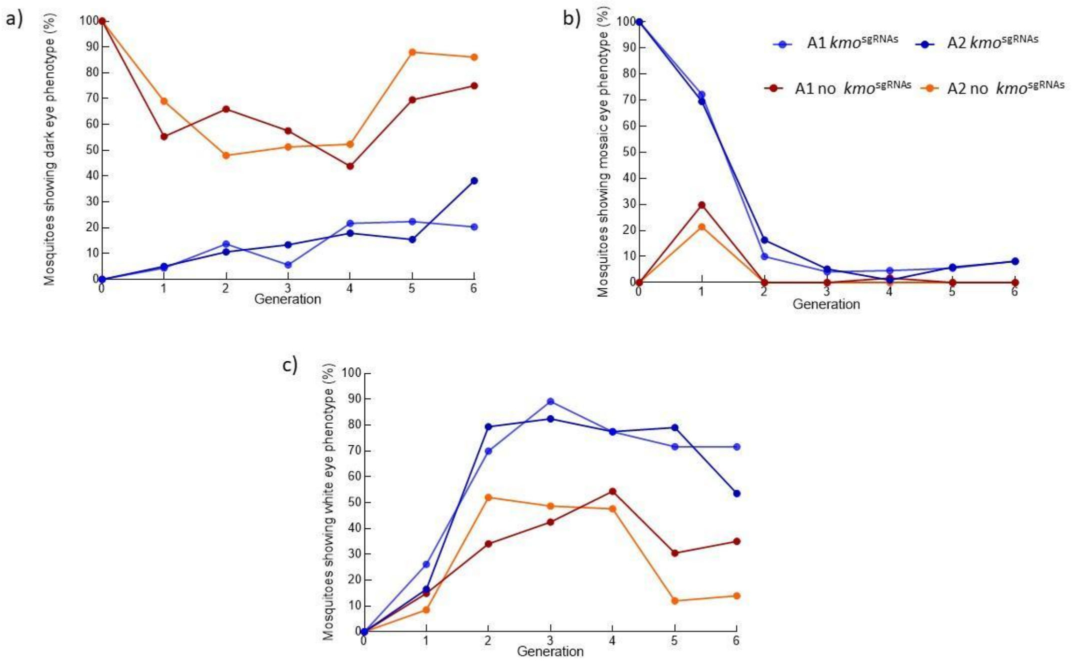


**Figure S7. Eye phenotypes tracked through cage trial.** Proportion of mosquitoes presenting dark eyes (a), mosaic eyes (b) or white eyes (c) screened from each generation of the cage trial experimental cages A1 and A2. Phenotypes have been separated based on the inheritance of the *kmo*$^{sgRNAs}$ element. Blue lines indicate those which inherited the *kmo*$^{sgRNAs}$ element; these would require disruption of one *kmo* allele to appear as mosaic (somatic disruption) or white eyed (germline disruption inherited from parents). Those which do not carry the *kmo*$^{sgRNAs}$ element are indicated in orange and can be either completely non-transgenic or carry the bgcn-Cas9. In these individuals two alleles of *kmo* would need to be disrupted to present a mosaic or white eyed phenotype.

**Table S1. Summary of microinjections performed to establish transgenic lines.**

| Construct | Embryos injected | $G_0$ survivors (%) | $G_1$s screened | Positive $G_1$s |
|---|---|---|---|---|
| ***kmo*$^{sgRNAs}$ (AGG1095)** | 1972 | 60 (3%) | 6345 | 46 (3/4 pools) |
| ***bgcn*-Cas9 (AGG1207)** | 1152 | 129 (11.2%) | 2238 | 38 (5/6 pools) |

**Table S2**. **Homozygous viability of *bgcn*-Cas9 lines.**

| ***bgcn*-Cas9 sibling cross** | **No. progeny inheriting *bgcn*-Cas9 (%)** | **No. WT progeny** | **TOTAL screened** | **Chi-squar e value** | **P value*** |
|---|---|---|---|---|---|
| **line B2** | 291 (64%) | 163 | 454 | 28.12 | <0.0001 |
| **line C** | 256 (74%) | 90 | 346 | 0.2476 | 0.6188 |
| **line D** | 202 (74%) | 69 | 271 | 0.0196 | 0.8886 |

WT: wild type

*Chi-square (df.1, p<0.0001) for deviation from expected Mendelian inheritance rate.

**Table S3. $F_1$ progeny from the *bgcn*-Cas9 x *kmo*$^{sgRNAs}$ $F_0$ cross.**

| $F_0$ Cross (male x female) | No. $F_1$ positive for *kmo*$^{sgRNAs}$ (mosaics) | No. $F_1$ positive for *kmo*$^{sgRNAs}$; *bgcn*-Cas9 (mosaics) | No. $F_1$ positive for *bgcn*-Cas9 (mosaics) | No. WT $F_1$ (mosaics) | Total F1 scored |
|---|---|---|---|---|---|
| ***kmo*$^{sgRNAs}$ x *bgcn*-Cas9D** | 29 (17) | 34 (27) | 23 (0) | 29 (0) | 105 (44) |

WT: wild type

**Table S4. Inheritance assessments of progeny from *kmo*$^{sgRNAs}$; *bgcn*-Cas9 transheterozygous $F_1$.**

| $F_1$ Cross (male x female) | No. $F_2$ positive for *kmo*$^{sgRNAs}$ | No. $F_2$ positive for *kmo*$^{sgRNAs}$; *bgcn*-Cas9 | No. $F_2$ positive for *bgcn*-Cas9 | No. WT $F_2$ | Total $F_2$ scored | Total $F_2$ inheriting *kmo*$^{sgRNAs}$ (%) | Total $F_2$ inheriting *bgcn*-Cas9 (%) |
|---|---|---|---|---|---|---|---|
| *kmo*$^{sgRNAs}$; *bgcn*-Cas9D x WT (exp 1) | 33 | 44 | 20 | 16 | 113 | 77 (68.1) | 64 (56.6) |
| *kmo*$^{sgRNAs}$; *bgcn*-Cas9D x WT (exp 2) | 431 | 367 | 155 | 618 | 1571 | 798 (50.8) | 522 (33.2) |
| *kmo*$^{sgRNAs}$; *bgcn*-Cas9D x WT (exp 3) | 213 | 198 | 154 | 190 | 755 | 411 (54.4) | 352 (46.8) |
| WT x *kmo*$^{sgRNAs}$; *bgcn*-Cas9D (exp 1) | 122 | 123 | 47 | 26 | 318 | 245 (77) | 170 (53.4) |
| WT x *kmo*$^{sgRNAs}$; *bgcn*-Cas9D (exp 2) | 66 | 45 | 17 | 19 | 147 | 111 (75.5) | 62 (42.1) |
| WT x *kmo*$^{sgRNAs}$; *bgcn*-Cas9D (exp 3) | 157 | 162 | 43 | 46 | 408 | 319 (78.7) | 205 (50.2) |

WT: wild type;

**Table S5. Mosaicism assessments of progeny from *kmo*$^{sgRNAs}$; *bgcn*-Cas9 trans-heterozygous $F_1$.**

| F1 Cross (male x female) | No. mosaics in the $F_2$ positive for *kmo*$^{sgRNAs}$ (%) | No. mosaics in the $F_2$ positive for *kmo*$^{sgRNAs}$; *bgcn*-Cas9 (%) | No. mosaics in the $F_2$ positive for *bgcn*-Cas9 (%) | No. mosaics in the WT $F_2$ (%) |
|---|---|---|---|---|
| ***kmo*$^{sgRNAs}$; *bgcn*-Cas9D x WT (exp 1)** | 2 (6) | 20 (45.5) | 0 | 0 |
| ***kmo*$^{sgRNAs}$; *bgcn*-Cas9D x WT (exp 2)** | 104 (21.4) | 235 (64) | 0 | 0 |
| ***kmo*$^{sgRNAs}$; *bgcn*-Cas9D x WT (exp 3)** | 25 (11.7) | 56 (28.3) | 0 | 0 |
| **WT x *kmo*$^{sgRNAs}$; *bgcn*-Cas9D (exp 1)** | 76 (62.3) | 87 (71) | 9 (19.1) | 1 (3.8) |
| **WT x *kmo*$^{sgRNAs}$; *bgcn*-Cas9D (exp 2)** | 65 (98.5) | 45 (100) | 3 (17.6) | 3 (15.8) |
| **WT x *kmo*$^{sgRNAs}$; *bgcn*-Cas9D (exp 3)** | 148 (94.2) | 157 (97) | 19 (44.1) | 15 (32.6) |

WT: wild type

**Table S6. G-test result data for $F_2$ rates of inheritance**. This is a maximum likelihood significance test. Individual crosses are analyzed for significant deviation from Mendelian inheritance. Individual crosses are pooled according to treatment for an overall Goodness of fit G-value and scored for whether there is heterogeneity among treatments.

| Line | df | G – goodness of fit | P-value |
|---|---|---|---|
| **D-Male** | 1 | 15.221 | 0.000096* |
| **D-Female** | 1 | 98.201 | 3.78E-23* |

| Pooled crosses | df | G – goodness of fit | P-value | df | Heterogeneity G-value | P-value |
|---|---|---|---|---|---|---|
| **D Line Mosaics only** | 1 | 110.03 | 2.2E-16 | 2 | 3.39 | 0.066 |

Table S7. Summary of the cutting assay screening from *kmo*$^{sgRNAs}$;*bgcn* Cas9 mosquitoes crossed to *kmo*$^{-/-}$ mosquitoes.

| Cross (male x female) | Replicate | No. $F_2$ positive for *kmo*$^{sgRNAs}$ | No. $F_2$ positive for *kmo*$^{sgRNAs}$; *bgcn*-Cas9 | No. $F_2$ positive for *bgcn*-Cas9 | No. WT $F_2$ | Total $F_2$ scored | Total $F_2$ inheriting *kmo*$^{sgRNAs}$ | Total $F_2$ non-inheriting *kmo*$^{sgRNAs}$ | Total $F_2$ non-inheriting *kmo*$^{sgRNAs}$ with WE |
|---|---|---|---|---|---|---|---|---|---|
| *kmo*$^{-/-}$ x (*kmo*$^{sgRNAs;}$ *bcgn*-Cas9 ) | 1 | 10 | 17 | 4 | 4 | 35 | 27 | 8 | 8 |
| | 2 | 16 | 6 | 8 | 12 | 42 | 22 | 20 | 20 |
| | 3 | 12 | 7 | 14 | 16 | 49 | 19 | 30 | 30 |
| | 4 | 23 | 22 | 13 | 8 | 66 | 45 | 21 | 5 |
| | 5 | 9 | 11 | 1 | 7 | 28 | 20 | 8 | 8 |
| | 6 | 22 | 35 | 6 | 6 | 69 | 57 | 12 | 12 |
| | 7 | 6 | 16 | 10 | 3 | 35 | 22 | 13 | 13 |
| | 8 | 26 | 24 | 17 | 10 | 77 | 50 | 27 | 13 |
| | 9 | 13 | 30 | 14 | 21 | 78 | 43 | 35 | 30 |
| | 10 | 33 | 38 | 4 | 1 | 76 | 71 | 5 | 5 |
| | 11 | 13 | 18 | 0 | 0 | 31 | 31 | 0 | 0 |
| | 12 | 25 | 24 | 3 | 1 | 53 | 49 | 4 | 4 |
| | 13 | 43 | 25 | 12 | 11 | 91 | 68 | 23 | 20 |
| | 14 | 22 | 20 | 2 | 3 | 47 | 42 | 5 | 4 |
| | 15 | 3 | 12 | 0 | 0 | 15 | 15 | 0 | 0 |
| | 16 | 16 | 19 | 1 | 0 | 36 | 35 | 1 | 0 |
| | 17 | 21 | 18 | 8 | 14 | 61 | 39 | 22 | 22 |

| | | | | | | | | | |
|---|---|---|---|---|---|---|---|---|---|
| | 18 | 4 | 10 | 1 | 4 | 19 | 14 | 5 | 5 |
| | 19 | 8 | 6 | 2 | 2 | 18 | 14 | 4 | 4 |
| | 20 | 16 | 14 | 2 | 7 | 39 | 30 | 9 | 9 |
| | 21 | 12 | 12 | 1 | 3 | 28 | 24 | 4 | 4 |
| | 22 | 5 | 7 | 3 | 9 | 24 | 12 | 12 | 12 |
| | 23 | 21 | 13 | 8 | 6 | 48 | 34 | 14 | 12 |
| | 24 | 4 | 7 | 10 | 3 | 24 | 11 | 13 | 9 |
| | 25 | 31 | 35 | 4 | 7 | 77 | 66 | 11 | 6 |
| | 26 | 9 | 4 | 1 | 1 | 15 | 13 | 2 | 2 |
| **(*kmo*$^{sgRNAs}$; *bcgn*-Cas9 ) x kmo$^{-/-}$** | 1 | 19 | 18 | 14 | 26 | 77 | 37 | 40 | 9 |
| | 2 | 14 | 7 | 10 | 6 | 37 | 21 | 16 | 15 |
| | 3 | 56 | 63 | 1 | 2 | 122 | 119 | 3 | 0 |
| | 4 | 10 | 17 | 17 | 8 | 52 | 27 | 25 | 15 |
| | 5 | 19 | 10 | 8 | 12 | 49 | 29 | 20 | 1 |
| | 6 | 27 | 20 | 22 | 20 | 89 | 47 | 42 | 7 |
| | 7 | 27 | 20 | 1 | 1 | 49 | 47 | 2 | 2 |
| | 8 | 16 | 22 | 18 | 20 | 76 | 38 | 38 | 6 |
| | 9 | 44 | 50 | 0 | 2 | 96 | 94 | 2 | 2 |
| | 10 | 19 | 27 | 18 | 9 | 73 | 46 | 27 | 7 |
| | 11 | 27 | 30 | 9 | 11 | 77 | 57 | 20 | 14 |
| | 12 | 30 | 26 | 23 | 29 | 108 | 56 | 52 | 2 |

| | | | | | | | | | |
|---|---|---|---|---|---|---|---|---|---|
| | 13 | 24 | 29 | 10 | 9 | 72 | 53 | 19 | 19 |
| | 14 | 19 | 11 | 18 | 15 | 63 | 40 | 33 | 8 |
| | 15 | 11 | 15 | 13 | 13 | 52 | 26 | 26 | 0 |
| | 16 | 2 | 3 | 1 | 3 | 9 | 5 | 4 | 1 |
| | 17 | 7 | 4 | 3 | 2 | 16 | 11 | 5 | 4 |
| | 18 | 2 | 5 | 5 | 2 | 14 | 7 | 7 | 1 |
| | 19 | 34 | 30 | 0 | 1 | 65 | 64 | 1 | 1 |
| | 20 | 50 | 33 | 0 | 0 | 83 | 83 | 0 | 0 |
| | 21 | 20 | 13 | 0 | 0 | 33 | 33 | 0 | 0 |
| | 22 | 16 | 14 | 6 | 10 | 46 | 30 | 16 | 16 |

WT: wild type; WE: white-eyed

**Table S8. Full model summaries for estimates of homing and cutting rates from *kmo*$^{sgRNAs}$; *bgcn* Cas9 mosquitoes crossed to *kmo*$^{-/-}$ mosquitoes. Coefficients, confidence intervals, and significance values are taken from a binomial glm or glmm with 'logit' link. Mixed-effects models included replicate as a nested random factor within the sex of the parent.**

| | **1.Mixed-effects model Homing rates** | **2.Mixed-effects model Combined male & female homing rates** | **3.Mixed-effects model Cutting rates** | **4.Pooled model for Homing rates** | **5.Pooled model for Cutting rates** |
|---|---|---|---|---|---|
| *Predictors* | *Log-Odds* | *Log-Odds* | *Log-Odds* | *Log-Odds* | *Log-Odds* |
| (Intercept) | 1.22 *** (0.70 – 1.74) | 1.18 *** (0.82 – 1.53) | 2.65 *** (1.57 – 3.72) | 0.92 *** (0.80 – 1.04) | 1.45 *** (1.32 – 1.59) |
| *kmo*$^{-/-}$ ♂ x *kmo*$^{sgRNAs}$; *bcgn*-Cas9♀ | -0.09 (-0.80 – 0.63) | N/A | 2.35 ** (0.77 – 3.93) | 0.07 (-0.10 – 0.25) | 1.61 *** (1.30 – 1.93) |
| **Random Effects** | | | | | |
| $\sigma^2$ | 4.64 | 4.64 | 8.38 | | |
| $\tau_{00}$ | 1.35 Replicate: Cross | 1.35 Replicate: Cross | 5.09 Replicate: Cross | | |
| | 0.00 Cross | 0.00 Cross | 0.00 Cross | | |
| N | 26 Replicate | 26 Replicate | 26 Replicate | | |
| | 2 Cross | 2 Cross | 2 Cross | | |
| Observations | 46 | 46 | 46 | 46 | 46 |
| Marginal $R^2$ / Conditional $R^2$ | 0.000 / NA | 0.000/ NA | 0.144 / NA | 0.007 | 0.073 |

* $p<0.05$ ** $p<0.01$ *** $p<0.001$

**Table S9. Percentage and types of *kmo* mutant alleles present in the *kmo*$^{-/-}$ line in the form of CIGAR strings. Cut sites of sgRNAs 447, 468, 499, and 519 are after nucleotides on positions 79, 100, 131, and 151, respectively.**

| **Genotype** | kmo$^{-/-}$ | | | |
|---|---|---|---|---|
| **No. of individuals** | 5 ♀ | 5 ♂ | 6 ♀ | 8 ♂ |
| **Total no. of reads** | 103,014 | 100,693 | 505,197 | 588,249 |
| **No. of reads mapped** | 99,796 | 97,987 | 498,107 | 568,133 |
| **Common CIGAR string obtained from sample** | | | | |
| **72M12D40M10D (%)** | 11,493 (11.5) | 31,805 (32.5) | 217,666 (43.7) | 147,901 (26.0) |
| **72M12D11M6D23M10D (%)** | 9,943 (10.0) | 6,809 (6.9) | 112,736 (22.6) | 46,052 (8.1) |
| **72M12D14M3D19M10D (%)** | 84 | 0 | 1 | 1,019 (0.2) |
| **98M3D24M5D (%)** | 2 | 11,093 (11.3) | 32,589 (6.5) | 6 |
| **98M3D19M10D (%)** | 14,463 (14.5) | 24 | 163 | 67,521 (11.9) |
| **98M3D24M7D (%)** | 7,841 (7.9) | 1 | 47 | 2 |

M: matches; D: deletions

**Table S10. Percentage of reads with deletions which may cause the loss of three or more sgRNA recognition sites.**

| **Parental ($F_1$) cross** | ♂ ***kmo***$^{-/-}$ **X** ♀ ***kmo***$^{sgRNAs}$**;** ***bgcn*****-Cas9** | | ♂ ***kmo***$^{sgRNAs}$**;** ***bgcn*****-Cas9 X** ♀ ***kmo***$^{-/-}$ | |
|---|---|---|---|---|
| **No. of individuals** | ***27*** | **24** | ***9*** | **11** |
| **Progeny ($F_2$) genotype/phenotype** | ***bgcn*****-Cas9/mosaic- or white-eyed** | **WT/mosaic- or white-eyed** | ***bgcn*****-Cas9/mosaic - or white-eyed** | **WT/mosaic- or white-eyed** |
| **Total no. of reads** | 3,538,107 | 3,266,941 | 1,657,916 | 2,292,377 |
| **No. of reads mapped** | 3,307,024 | 3,032,896 | 1,456,448 | 2,094,102 |
| **No. of reads with deletion sizes of:** | | | | |
| **46-50 nt (%)** | 126,120 (3.8) | 0 | 1,380 (0.09) | 13 (0.0006) |
| **51-55 nt (%)** | 425 (0.01) | 2,040 (0.07) | 439 (0.03) | 0 |
| **56-60 nt (%)** | 907 (0.03) | 13,759 (4.5) | 374 (0.03) | 1 |
| **61-65 nt (%)** | 3 | 1,937 (0.06) | 0 | 0 |
| **66-70 nt (%)** | 0 | 3 | 0 | 0 |
| **71-75 nt (%)** | 0 | 0 | 0 | 0 |

nt: nucleotides; WT: wild type

**Table S11. Complete primer list.**

| Name | Sequence (5'-3') |
|---|---|
| LA174 | AGGATGTCGAAGGAGAAGGCCAGG |
| LA182 | GGCGACTGAGATGTCCTAAATGCAC |
| LA184 | CAGACCGATAAAACACATGCGTCA |
| LA186 | CAGCGACGGATTCGCGCTATTTAG |
| LA187 | GTGTAGCGTGAAGACGACAGAA |
| LA518 | GCCTCCTGAATCCAAATATGCTTGC |
| LA924 | GCACCGAATCGGTGCCTGCCTTCCGGCATGATAACGGACTTGCCTTATTCCAAC TTGTCGTGCTGTTCCCAGCACGACTCTGGAAC |
| LA925 | GAAATTAATACGACTCACTATAGGGCCATATAATGTGGGCGGCAGTTCCAGAGTC GTGCTGG |
| LA926 | GAAATTAATACGACTCACTATAGGCACAGTACAATCCTCGAATCGTTCCAGAGTC GTGCTGG |
| LA927 | GAAATTAATACGACTCACTATAGGGGTTCCCTTCTACGGGCAGTTCCAGAGTCG TGCTGG |
| LA928 | GAAATTAATACGACTCACTATAGGCGGTGATCATTGGTGATGGTTCCAGAGTCGT GCTGG |
| LA1074 | CGGCCATTTACGATCGGTGGGTTTGG |
| LA1075 | GCGTTGTTCTGCGGTCCCCTGTTTTTG |
| LA1076 | CTGGCCAATGTTACTGTGGCCGGCG |
| LA1275 | TTATGATGATCGCCCTGCCC |
| LA1301 | TGGCCTTCTCCTTCGACATCCTGT |
| LA1352 | GATTACGCCAAGCTTGATGGTTCCTCATGACCTGCGCCGC |
| LA1725 | TTTTGCGGCCGCCAACGTTGGGGCGTCATAAG |
| LA1726 | TTTTCTCGAGGTTGGAGCTGTTTTCGTT |
| LA1737 | TTTTTTAATTAAATCTTGATACGTCTCTTCATCAAGC |
| LA1738 | TTTTGCGATCGCCCTCGAGCTATGTTTAATTTGTCATTCTTTCACATTG |
| LA2750 | TAAGTGTTTCGCAGACGGCTTCA |
| LA2755 | ACGCATGTGGGAGAACGATA |

**Table S12. Deep sequencing analysis for cutting assessment.**

| Sample name | Raw reads | Trimmed reads |
|---|---|---|
| G16_B_WE_S125_R1_001 | 13658 | 13637 |
| G18_B_WE_S126_R1_001 | 21127 | 20758 |
| G19_B_WE_S127_R1_001 | 30507 | 29195 |
| G19_WT_WE_S128_R1_001 | 20816 | 20749 |
| G1_B_WE_S123_R1_001 | 16446 | 16232 |
| G1_WT_WE_S124_R1_001 | 18647 | 18621 |
| G22_B_WE_S129_R1_001 | 23105 | 22879 |
| G22_WT_WE_S130_R1_001 | 18878 | 18865 |
| G23_WT_WE_S131_R1_001 | 196 | 103 |
| G24_B_WE_S132_R1_001 | 21857 | 21287 |
| G24_WT_WE_S133_R1_001 | 28354 | 28340 |
| G25_B_WE_S134_R1_001 | 26329 | 26156 |
| G25_WT_WE_S135_R1_001 | 30307 | 30271 |
| G27_B_WE_S136_R1_001 | 22552 | 22331 |
| G30_BE_WE_S137_R1_001 | 31352 | 31284 |
| G30_WT_WE_S138_R1_001 | 28403 | 28373 |
| G32_BE_WE_S139_R1_001 | 30415 | 30396 |
| G34_BE_WE_S140_R1_001 | 19080 | 19045 |
| G37_B_WE_S141_R1_001 | 29865 | 29812 |
| G37_WT_WE_S142_R1_001 | 20610 | 20594 |
| G38_B_WE_S143_R1_001 | 17684 | 17674 |
| G42_B_WE_S144_R1_001 | 20926 | 20879 |
| G42_WT_WE_S145_R1_001 | 25817 | 25716 |
| G45_WT_WE_S146_R1_001 | 42293 | 41635 |
| G46_B_WE_B_S148_R1_001 | 31922 | 31895 |
| G46_B_WE_S147_R1_001 | 20372 | 20352 |
| G46_WT_WE_S149_R1_001 | 25862 | 25476 |
| G50_B_WE_S150_R1_001 | 29954 | 29914 |
| G50_WT_WE_S151_R1_001 | 23440 | 23311 |
| G51_B_WE_S152_R1_001 | 25516 | 25492 |
| G6_B_WE_S153_R1_001 | 24602 | 24546 |
| G9_B_WE_S154_R1_001 | 22343 | 22313 |
| G9_WT_WE_S155_R1_001 | 19634 | 19596 |
| GM10_B_WE_S156_R1_001 | 18252 | 18230 |
| GM10_WT_WE_S157_R1_001 | 29860 | 29747 |
| GM12_B_WE_S158_R1_001 | 31970 | 31885 |
| GM14_B_WE_S159_R1_001 | 29932 | 29825 |
| GM14_WT_WE_S160_R1_001 | 18373 | 18368 |
| GM16_B_WE_B_S161_R1_001 | 20528 | 20402 |
| GM16_WT_WE_S162_R1_001 | 35259 | 35226 |

| | | |
|---|---|---|
| GM18_B_WE_B_S163_R1_001 | 29407 | 29267 |
| GM18_WT_WE_S164_R1_001 | 18495 | 18479 |
| GM24_B_WE_B_S167_R1_001 | 33939 | 33921 |
| GM24_WT_WE_B_S168_R1_001 | 35980 | 35094 |
| GM27_B_WE_S169_R1_001 | 30 | 30 |
| GM28_B_WE_S170_R1_001 | 29876 | 29633 |
| GM29_B_WE_S171_R1_001 | 31949 | 31916 |
| GM29_WT_WE_S172_R1_001 | 30186 | 30106 |
| GM2_B_WE_S165_R1_001 | 31926 | 31537 |
| GM2_WT_WE_S166_R1_001 | 23293 | 23218 |
| GM31_WT_WE_S173_R1_001 | 19803 | 19760 |
| GM32_B_WE_S174_R1_001 | 35815 | 35711 |
| GM35_B_WE_S175_R1_001 | 22807 | 22609 |
| GM5_B_WE_S176_R1_001 | 26327 | 26308 |
| GM9_B_WE_B_S177_R1_001 | 23906 | 23838 |
| GM9_WT_WE_B_S178_R1_001 | 25801 | 25709 |
| KMO_KO_5females_S179_R1_001 | 173303 | 12575 |
| KMO_KO_5males_S180_R1_001 | 199913 | 19701 |

**Table S13. Deep sequencing analysis of cage trial samples**

| Sample name | Raw reads | Trimmed reads |
|---|---|---|
| F2_1A_A_B_DE2_S21_R1_001 | 13784 | 13777 |
| F2_1A_A_B_DE3_S22_R1_001 | 29718 | 29400 |
| F2_1A_A_B_ME1_S23_R1_001 | 74605 | 74551 |
| F2_1A_A_B_ME2_S24_R1_001 | 37813 | 37767 |
| F2_1A_A_B_WE1_S25_R1_001 | 47942 | 47905 |
| F2_1A_A_B_WE2_S26_R1_001 | 27971 | 27580 |
| F2_1A_A_DE_S27_R1_001 | 45674 | 45636 |
| F2_1A_A_WE_S28_R1_001 | 39313 | 39132 |
| F2_1A_B_WE_S29_R1_001 | 49228 | 49196 |
| F2_1A_WT_WE1_S30_R1_001 | 46068 | 45992 |
| F2_1A_WT_WE2_S31_R1_001 | 46004 | 45467 |
| F2_1B_A_B_DE_S32_R1_001 | 48354 | 48307 |
| F2_1B_A_B_ME1_S33_R1_001 | 9424 | 9419 |
| F2_1B_A_B_ME2_S34_R1_001 | 42933 | 42899 |
| F2_1B_A_B_ME3_S35_R1_001 | 37963 | 37933 |
| F2_1B_A_DE1_S36_R1_001 | 10038 | 9965 |
| F2_1B_A_DE2_S37_R1_001 | 9314 | 9282 |
| F2_1B_A_WE1_S38_R1_001 | 4608 | 4606 |
| F2_1B_A_WE2_S39_R1_001 | 843 | 838 |
| F2_1B_WT_DE_S40_R1_001 | 30728 | 30683 |
| F2_1B_WT_WE1_S41_R1_001 | 9133 | 9087 |
| F4_1A_A_B_DE_B_S44_R1_001 | 49865 | 49826 |
| F4_1A_A_B_DE_S43_R1_001 | 9244 | 9225 |
| F4_1A_A_B_ME_S45_R1_001 | 44162 | 44113 |
| F4_1A_A_B_WE_S46_R1_001 | 58 | 41 |
| F4_1A_A_DE1_S47_R1_001 | 53557 | 53505 |
| F4_1A_A_DE2_S48_R1_001 | 42279 | 42259 |
| F4_1A_A_DE3_S49_R1_001 | 3609 | 3592 |
| F4_1A_A_DE4_S50_R1_001 | 4205 | 4137 |
| F4_1A_A_WE_S51_R1_001 | 2666 | 2664 |
| F4_1A_B_WE1_S52_R1_001 | 7083 | 6847 |
| F4_1A_B_WE2_S53_R1_001 | 149540 | 149415 |
| F4_1A_B_WE3_S54_R1_001 | 63642 | 63611 |
| F4_1A_WT_WE1_S55_R1_001 | 62630 | 62570 |
| F4_1A_WT_WE2_S56_R1_001 | 12980 | 12794 |
| F4_1B_A_B_DE1_S57_R1_001 | 48594 | 47963 |
| F4_1B_A_B_DE2_S58_R1_001 | 16930 | 16900 |
| F4_1B_A_B_DE3_S59_R1_001 | 11432 | 11321 |
| F4_1B_A_B_WE_S60_R1_001 | 172 | 171 |
| F4_1B_A_DE1_S61_R1_001 | 5616 | 5591 |

| | | |
|---|---|---|
| F4_1B_A_DE2_S62_R1_001 | 7598 | 7570 |
| F5_1A_A_ME1_S83_R1_001 | 29189 | 29130 |
| F5_1A_A_WE1_B_S85_R1_001 | 52947 | 52881 |
| F5_1A_A_WE1_S84_R1_001 | 27834 | 25559 |
| F5_1A_A_WE2_B_S87_R1_001 | 47585 | 47547 |
| F5_1A_A_WE2_S86_R1_001 | 30055 | 28165 |
| F5_1A_A_WE3_S88_R1_001 | 23860 | 23818 |
| F5_1A_B_WE_S89_R1_001 | 54126 | 53945 |
| F5_1A_WT_WE1_B_S91_R1_001 | 64447 | 64358 |
| F5_1A_WT_WE1_S90_R1_001 | 24712 | 24676 |
| F5_1A_WT_WE2_B_S93_R1_001 | 55185 | 55156 |
| F5_1A_WT_WE2_S92_R1_001 | 34089 | 33713 |
| F5_1A_WT_WE3_S94_R1_001 | 25393 | 25324 |
| F5_1B_A_B_DE1_B_S96_R1_001 | 84081 | 84026 |
| F5_1B_A_B_DE1_S95_R1_001 | 23633 | 23533 |
| F5_1B_A_B_DE2_B_S98_R1_001 | 50624 | 50579 |
| F5_1B_A_B_DE2_S97_R1_001 | 19088 | 19071 |
| F5_1B_A_B_DE3_B_S100_R1_001 | 37346 | 37305 |
| F5_1B_A_B_DE3_S99_R1_001 | 16791 | 16744 |
| F5_1B_A_B_DE4_B_S102_R1_001 | 12038 | 12028 |
| F5_1B_A_B_DE4_S101_R1_001 | 19171 | 18998 |
| F5_1B_A_B_DE5_S103_R1_001 | 55450 | 55412 |
| F5_1B_A_B_WE1_B_S105_R1_001 | 523 | 507 |
| F5_1B_A_B_WE1_S104_R1_001 | 265 | 123 |
| F5_1B_A_B_WE2_B_S107_R1_001 | 13449 | 13440 |
| F5_1B_A_B_WE2_S106_R1_001 | 9559 | 7747 |
| F5_1B_A_DE1_B_S109_R1_001 | 3 | 2 |
| F5_1B_A_DE1_S108_R1_001 | 18303 | 18060 |
| F5_1B_A_DE2_B_S111_R1_001 | 57524 | 57481 |
| F5_1B_A_DE2_S110_R1_001 | 25455 | 25414 |
| F5_1B_A_DE3_B_S113_R1_001 | 52922 | 52798 |
| F5_1B_A_DE3_S112_R1_001 | 22987 | 22974 |
| F5_1B_A_DE4_B_S115_R1_001 | 31165 | 31100 |
| F5_1B_A_DE4_S114_R1_001 | 23657 | 23645 |
| F5_1B_A_WE1_B_S117_R1_001 | 2274 | 428 |
| F5_1B_A_WE1_S116_R1_001 | 28242 | 28176 |
| F5_1B_A_WE2_B_S119_R1_001 | 4120 | 3049 |
| F5_1B_A_WE2_S118_R1_001 | 406 | 270 |
| F5_1B_A_WE3_B_S121_R1_001 | 4501 | 884 |
| F5_1B_A_WE3_S120_R1_001 | 20315 | 3229 |
| F5_1B_A_WE4_S122_R1_001 | 21592 | 21574 |

**Table S14. Closest potential off-targets as determined by CHOPCHOP, lowercase nucleotides indicate mismatches**

| Target: | sequence | Potential off targets |
|---|---|---|
| **kmo447** | GCCATATAATGTGGGCGGCAAGG | none |
| **kmo468** | GGCGGTGATCATTGGTGATGCGG | GGCcGTtgTCATTGGTGATGGGG<br>GGCGGTGATCATTGtgGATcCGG<br>GGCacTGATCATTGGTGATcTGG<br>GGgGGTGATCATTGGgGgTGGGG<br>GGCGGaGATCtTTGcTGATGTGG |
| **kmo499** | GGTTCCCTTCTACGGGCAGGG | GGTaCCCTTCTgaGGGCAGGG |
| **kmo519** | CACAGTACAATCCTCGAATCCGG | CACgccACAATCtTCGtATCGGG |

**Dataset S1 (separate file).** Cage trial results:

**-Cage trial screening. Screening results for each generation of the cage trial.** F1 to F6 progeny results classified by eye phenotype (DE,dark eyes; ME, mosaic eyes; WE, white eyes) for the multi-generational cage trial. TH (trans-heterozygous, *kmo*$^{sgRNAs}$;*bgcn*-Cas9D).

**-Eye phenotype. Number of individuals with mosaic/white eye phenotype in *kmo*$^{sgRNAs}$ and non-*kmo*$^{sgRNAs}$ mosquitoes in each generation.** *kmo*$^{sgRNAs}$ = all progeny that inherited the *kmo*$^{sgRNAs}$ element (*kmo*$^{sgRNAs}$ only as well as trans-heterozygotes); Non-*kmo*$^{sgRNAs}$ = progeny that did not inherit the *kmo*$^{sgRNAs}$ element (*bgcn*-Cas9 only and wild type).

**-Genotype. Phenotype frequencies for each generation of the cage trial assay.** Phenotype frequencies calculated from the F1-F6 generations (Number of mosquitoes showing a phenotype/total mosquitoes screened). TH: trans-heterozygous; WT: wild type.

**-Larvae. Number of larvae estimated in each experimental cage from F1 to F6.**
